# Ectosomes and exosomes are distinct proteomic entities that modulate spontaneous activity in neuronal cells

**DOI:** 10.1101/2021.06.24.449731

**Authors:** Inês Caldeira Brás, Mohammad Hossein Khani, Dietmar Riedel, Iwan Parfentev, Ellen Gerhardt, Christoph van Riesen, Henning Urlaub, Tim Gollisch, Tiago Fleming Outeiro

**Affiliations:** Department of Experimental Neurodegeneration, Center for Biostructural Imaging of Neurodegeneration, University Medical Center Göttingen, 37073 Göttingen, Germany.; Department of Ophthalmology, University Medical Center Göttingen, 37073 Göttingen, Germany.; Laboratory of Electron Microscopy, Max Planck Institute for Biophysical Chemistry, 37075 Göttingen, Germany.; Research Group Bioanalytical Mass Spectrometry, Max-Planck-Institute for Biophysical Chemistry, 37077 Göttingen, Germany.; Department of Neurology, University Medical Center Göttingen, 37075 Göttingen, Germany.; German Center for Neurodegenerative Diseases (DZNE), 53127 Bonn, Germany.; Bioanalytics, Institute of Clinical Chemistry, University Medical Center Göttingen, 37075 Göttingen, Germany.; Max Planck Institute for Experimental Medicine, 37075 Göttingen, Germany.; Translational and Clinical Research Institute, Faculty of Medical Sciences, Newcastle University, NE2 4HH, United Kingdom.; Scientific employee with an honorary contract at German Center for Neurodegenerative Diseases (DZNE), 37075 Göttingen, Germany.

**Author notes:** Lead contact: Prof. Dr. Tiago Fleming Outeiro Department of Experimental Neurodegeneration, University Medical Center Göttingen, 37073 Göttingen, Germany.

**Keywords:** Extracellular vesicles, ectosomes, microvesicles, exosomes, spreading, proteomics, multi-electrode array, neuronal activity

## Abstract

Extracellular vesicles (EVs) are important mediators in intercellular communication. However, understanding the biological origin and functional effects of EVs subtypes has been challenging due to the moderate differences in their physical properties and absence of reliable markers. Here, we characterize the proteomes of ectosomes and exosomes using an improved differential ultracentrifugation protocol and quantitative proteomics. Cytoskeleton and glycolytic proteins are distinctively present in ectosomes, while endosomal sorting complexes proteins and tetraspanins are enriched in exosomes. Furthermore, annexin-A2 was identified as a specific marker for ectosomes derived from cell media and human cerebrospinal fluid. Expression of EGFP as a cytosolic reporter leads to its incorporation in EVs and enables their imaging with higher resolution. Importantly, ectosomes and exosomes internalization in neuronal cells results in the modulation of neuronal spontaneous activity. Our findings suggest that EVs cargoes reflect core intracellular processes, and their functional properties might regulate basic biological and pathological processes.

## Introduction

Extracellular vesicles (EVs) are important vehicles in intercellular communication, mediating long range signaling events (You and Ikezu, 2019, Meldolesi, 2018, van Niel et al., 2018, Mathieu et al., 2019). These vesicles are important in the central nervous system (CNS), where they are secreted by diverse cell types, appearing also in the cerebrospinal fluid (CSF) (Chiasserini et al., 2014, Coleman and Hill, 2015). Proteins, RNA, and lipids are actively and selectively incorporated into EVs, justifying the importance of these vesicles not only in normal biology but also in disease, as they may report on pathological alterations (Choi et al., 2013, Bellingham et al., 2012, Thompson et al., 2016).

Size and mechanisms of biogenesis are the conventional classification approaches for EVs (Thery et al., 2018, Cocucci and Meldolesi, 2015, Kalra et al., 2016). Exosomes (30 -150nm in diameter) are derived from endosomes released from multivesicular bodies (MVBs) after fusion with the plasma membrane (Mathivanan et al., 2010, Willms et al., 2016). In contrast, microvesicles (also termed ectosomes, 100 -1000 nm in diameter) are larger EVs generated by direct shedding from the plasma membrane (Sadallah et al., 2011, Stein and Luzio, 1991).

While exosomes have been characterized in multiple studies, ectosomes remain understudied. Furthermore, disparities in purification strategies and lack of reliable protein markers that can discriminate between these EVs types limits our knowledge regarding ectosomes (Choi et al., 2013, Choi et al., 2015, van Niel et al., 2018). Therefore, further characterization of ectosomes may provide valuable information on biogenesis, cargo sorting and functional roles of these vesicles in physiological and pathological conditions.

Herein, we provide an indepth molecular and functional characterization of ectosomes and exosomes based on differential ultracentrifugation. Comprehensive proteomic analysis revealed specific protein composition and pathway enrichment for each EV subtype. Exosomes are composed of endosomal sorting proteins required for transport (ESCRT) and tetraspanins (Jeppesen et al., 2019). In contrast, ectosomes are enriched in cytoskeletal proteins, glycolytic enzymes, integrins and annexins. Interestingly, ectosomes isolated from human CSF and from cell media are enriched in annexin-A2, suggesting this protein can be exploited as an important marker for ectosomes characterization. EGFP incorporation in both ectosomes and exosomes enabled their imaging at higher resolution when compared with the use of the thiol-based dye Alexa Fluor 633 C5-maleimide. Remarkably, we demonstrate that EVs internalization affects the spontaneous activity of primary cortical neurons. Our work provides novel insight into the cell biology of intercellular communication via EVs, demonstrating they transfer cargoes that can modulate cellular function. Ultimately, our study also forms the foundation for future biomarker studies and for the understanding of the molecular basis of different diseases.

## Results

### Separation of ectosomes and exosomes by differential ultracentrifugation

Biological samples contain a heterogeneous mixture of EVs. To understand their composition and functional properties, ectosomes and exosomes were isolated from the media of human HEK cells using an improved differential ultracentrifugation protocol (Figure 1A) (Dujardin et al., 2014, Thery et al., 2006). To avoid possible contamination with EVs from fetal bovine serum (FBS) present in the cell media, cells were incubated with conditioned media (previously depleted of EVs) for 24 hours, as previously described (Thery et al., 2006, Shelke et al., 2014).

**Figure 1.**
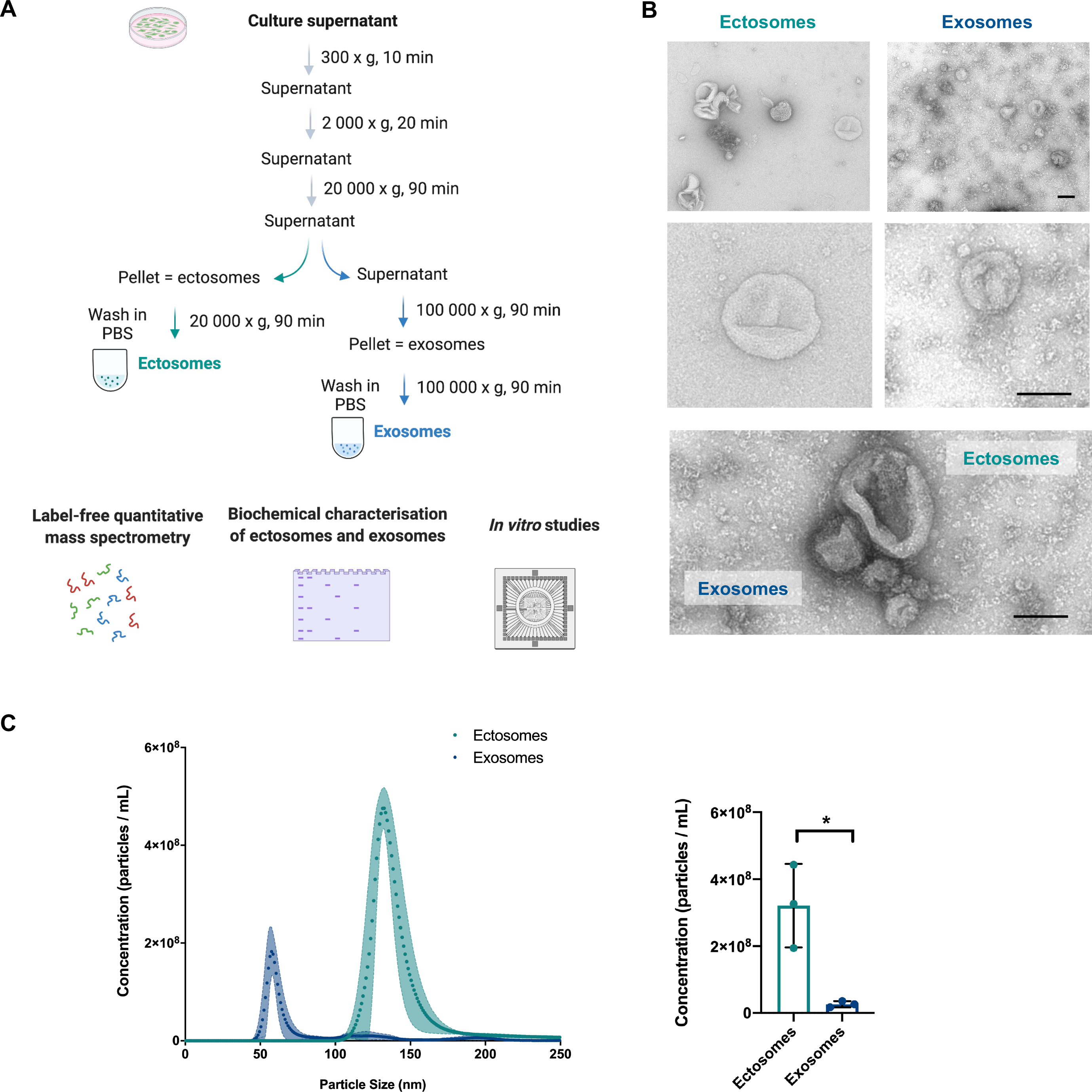
Purification and characterization of secreted EVs using differential centrifugation. **(A)** Schematic overview of the EVs purification protocol. Conditioned cell media was collected from HEK cells after 24 hours and subsequently centrifuged at different speeds. After isolation, EVs were used for label-free quantitative mass spectrometry, immunoblot and MEA recordings. **(B)** Whole-mount EM analysis of each pellet showing representative images of ectosomes and exosomes (scale bar 100nm). **(C)** NTA measurements of particle concentrations and average size distributions of ectosomes and exosomes. Data from at least three independent experiments. Significant differences were assessed by two-tailed unpaired t-test comparison and are expressed as mean ± SD, *p<0.05.

Both EVs subtypes exhibited a cap shaped morphology, typical for negative stained vesicular structures, and presented the expected size differences by electron microscopy (EM) (Figure 1B). Nanoparticle tracking analysis (NTA) was used to determine the size and to quantity of ectosomes and exosomes based on Brownian motion (Figure 1C). The diameter distribution for ectosomes was considerably larger and peaked at 140nm, while exosomes had an average diameter of 60nm (Figure 1C). Furthermore, the concentration of ectosomes was significantly greater when compared to exosomes (Figure 1C).

### Ectosomes and exosomes exhibit characteristic proteomic profiles

To determine the protein composition of ectosomes and exosomes, label-free quantitative mass spectrometry was performed (Figure 2). In total, 2216 proteins were identified in our study, with 371 proteins exclusively recognized in ectosomes, and 193 proteins enriched in exosomes (Figure 2A, Supplementary Tables 1 and 2).

**Figure 2.**
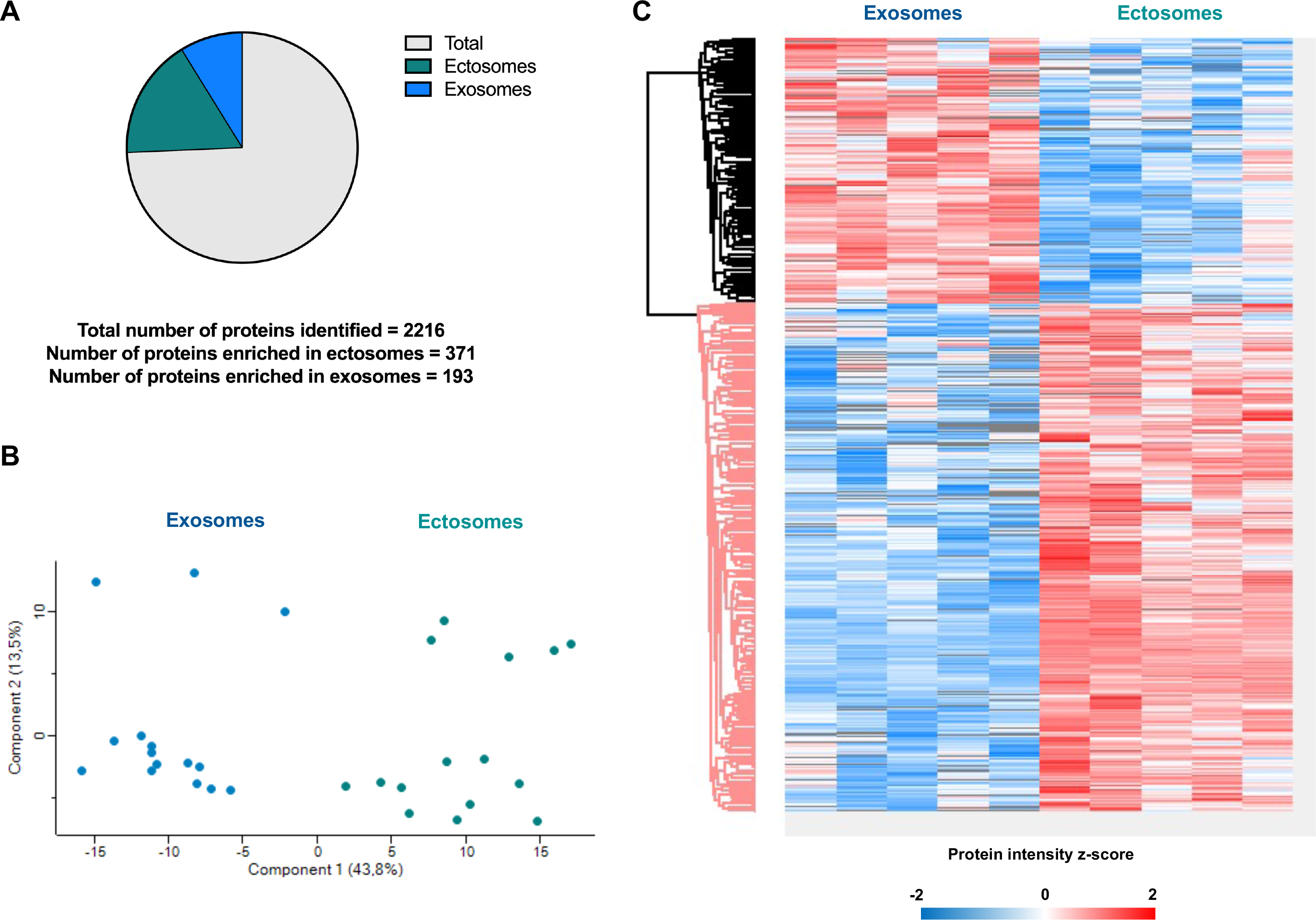
Proteomic analyses of ectosomes and exosomes using label-free quantitative mass spectrometry. **(A)** Diagram representing the number of unique proteins identified in each EV type and total number of proteins identified in the study. **(B)** PCA of the quantitative differences in spectral counts between ectosomes (turquoise) and exosomes (blue) in the biological replicates. **(C)** Heatmap of the significantly different identified proteins from proteomic profiling of in ectosomes and exosomes. High and low expression are shown in red and blue, respectively. Data represented from five independent samples. Data analyses were performed using Perseus software.

Principal component analysis (PCA) of protein composition revealed a clear separation of the two EVs subtypes (Figure 2B). Despite some overlap in identifiable proteins, which was to be expected, hierarchical clustering of the protein datasets uncovered two separate fractions with distinct proteomic profiles (Figure 2C).

These results indicate individual proteomic profiles of ectosomes and exosomes, supporting functional biological differences between the two EVs subtypes.

### Exosomes are enriched with tetraspanins and ESCRT-related proteins, while ectosomes contain cytoskeletal proteins and glycolytic enzymes

Next, we searched for specific markers of exosomes and ectosomes among the proteins uniquely present in each EV subtype (Kalra et al., 2016). Proteomic profiling identified a number of known exosomal proteins, including ESCRT and vacuolar protein sorting-associated (VPS) proteins (Supplementary Table 1). This included alix (*PDCD6IP*), syntenin-1 (*SDCBP*), vesicle-associated membrane protein 2 (*VAMP2*) and VPS25/ 28/ 37B (*VPS25/ 28/ 37B*) (Figure 3A, Table 1). Since the ESCRT machinery is important for the sorting of ubiquitinated cargo and regulation of exosomal biogenesis, their identification in our exosomal fraction is consistent with previous studies (Kowal et al., 2016, Zhang et al., 2018, Thery et al., 2006, Jeppesen et al., 2019, Vietri et al., 2020). Tetraspanins are highly enriched in exosomes and are associated with exosome biogenesis (Andreu and Yanez-Mo, 2014). Consistently, our exosomal fraction comprised CD63 antigen (*CD63*), CD81 antigen (*CD81*) and tetraspanin-4, 6, 7, 9 (*TSPAN4*/ 6/ 7/ 9) (Figure 3B, Supplementary Table 1). Furthermore, our proteomic analysis also highlighted the presence of lipid raft components in exosomes, such as flotillin-1 and 2 (*FLOT1*/ 2). However, their levels were not statistically different from the ones found in ectosomes. Similarly, TSG101 (*TSG101*) and CD9 (*CD9*) were also slightly enriched in exosomes, although this did not reach statistical significance in comparison to the levels found in ectosomes. Importantly, our results indicate that certain proteins commonly considered exosomal can also occur in ectosomes, emphasizing the need for using multiple protein markers to distinguish EVs.

**Figure 3.**
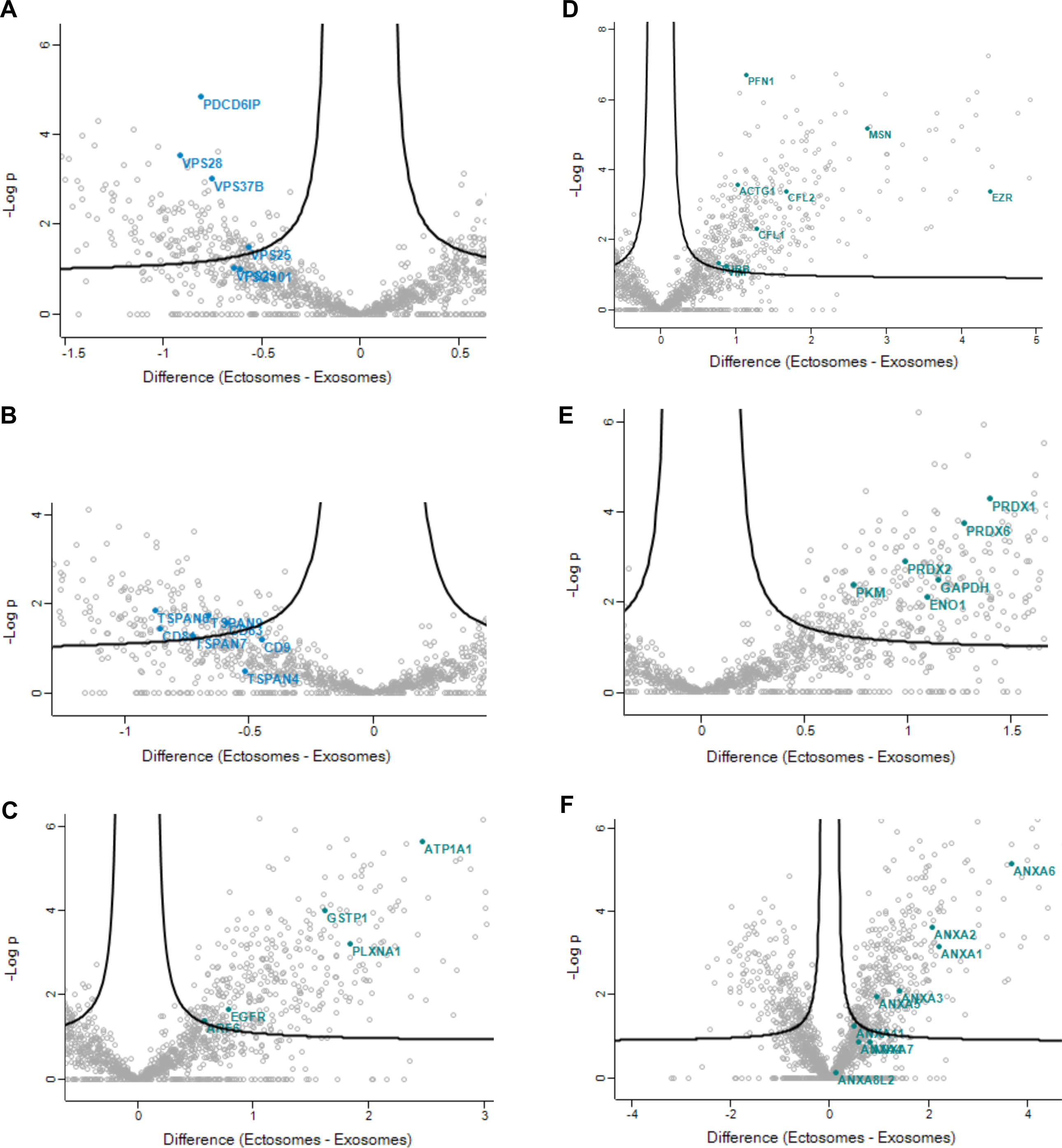
Protein hits enriched in ectosomes and exosomes. Volcano plots of quantitative differences in proteins in EVs fractions. **(A)** Endosomal sorting complexes required for transport (ESCRT) proteins present in exosomes - alix (*PDCD6IP*), tumor susceptibility 101 (*TSG101*) and vacuolar protein sorting-associated proteins 25/ 28/ 29/ 37B (*VPS25/ 28/ 29/ 37B*). **(B)** Tetraspanins proteins enriched in exosomes - CD9 antigen (*CD9*), CD63 antigen (*CD63*), CD81 antigen (*CD81*) and tetraspanin-4, 6, 7, 9 (*TSPAN4/ 6/ 7/ 9*). **(C)** Commonly identified proteins in larger vesicles, as ectosomes - plexin-A1 (*PLXA1*), glutathione S-transferase (*GSTP1*), sodium/potassium-transporting ATPase (*ATP1A1*), ADP-ribosylation factor 6 (*ARF6*) and Epidermal growth factor receptor (*EGFR*). **(D)** Cytoskeletal proteins enriched in ectosomes - profilin-1 (*PFN1*), cofilin-1/ 2 (*CFL1/ 2*), vimentin (*VIM*), ezrin (*EZR*), moesin (*MSN*) tubulin beta chain (*TUBB*) and actin (*ACTG1*). **(E)** Metabolic enzymes identified in ectosomes - peroxiredoxin-1/ 2/ 6 (*PRDX1/ 2/ 6*), pyruvate kinase (*PKM*), alpha-enolase (*ENO1*) and glyceraldehyde-3-phosphate dehydrogenase (*GAPDH*). **(F)** Annexin proteins enriched in ectosomes - annexin A1, A2, A3, A4, A5, A6, A7, A8, A11 (*ANXA1/ 2/ 3/ 4/ 5/ 6/ 7/ 8/ 11*). Blue dots represent the proteins enrichment in exosomes, while turquoise dots represent enrichment in ectosomes. Proteins in the plots are identified with their gene name. Dots above the dashed line represent proteins for which differences were significant (false discovery rate [FDR]<0.1). Data represented in “t-test Difference (Ectosomes - Exosomes)” vs. “-Log t-test p-value” from five independent samples for each group. Data analyses were performed using Perseus software. See also Figure S1.

**Table 1.** List of top 30 up-regulated proteins in exosomes identified in this study organized according to the significance values (full list in Supplementary Table 1).

| Protein names | Gene names | Biological process | $-\log_{10}(\text{p-value})$ |
| --- | --- | --- | --- |
| Alix | *PDCD6IP* | Cell cycle | 4,836 |
| Glycogen phosphorylase | *PYGM/ L* | Alcohol metabolic process | 4,300 |
| Proteasome subunit alpha/ beta | *PSMA/ B* | Protein catabolic process | 4,098 |
| Asporin | *ASPN* | Anatomical structure development | 3,734 |
| Homogentisate 1,2-dioxygenase | *HGD* | Amine metabolic process | 3,611 |
| Receptor-type tyrosine-protein phosphatase | *PTPRM* | Axon guidance | 3,610 |
| Vacuolar protein sorting-associated protein 28 homolog | *VPS28* | Catabolic process | 3,525 |
| Cation-independent mannose-6-phosphate receptor | *IGF2R* | Anatomical structure morphogenesis | 3,377 |
| Eukaryotic initiation factor 4A-II | *EIF4A2* | Biological regulation | 3,345 |
| Protein kinase C-binding protein NELL2 | *NELL2* | Cellular homeostasis | 3,340 |
| 3-mercaptopyruvate sulfurtransferase | *MPST* | Biosynthetic process | 3,327 |
| Mannose-1-phosphate guanylttransferase beta | *GMPPB* | Alcohol biosynthetic process | 3,264 |
| Cytosolic 10-formyltetrahydrofolate dehydrogenase | *ALDH1L1* | 10-formyltetrahydrofolate catabolic process | 3,226 |
| Cytoplasmic aconitate hydratase | *ACO1* | Acetyl-CoA catabolic process | 3,172 |
| S-adenosylmethionine synthase isoform type-1 | *MAT1A* | Amine biosynthetic process | 3,097 |
| Delta-aminolevulinic acid dehydratase | *ALAD* | Biosynthetic process | 3,039 |
| Tubulin alpha-4A chain | *TUBA4A* | Cell cycle | 3,033 |
| Vacuolar protein sorting-associated protein 37B | *VPS37B* | Cellular component assembly | 3,012 |
| Adenosylhomocysteinase | *AHCY* | Amine metabolic process | 2,989 |
| Regucalcin | *RGN* | Calcium ion homeostasis | 2,989 |
| Fructose-bisphosphate aldolase B | *ALDOB* | Alcohol biosynthetic process | 2,989 |
| 4-trimethylaminobutyraldehyde dehydrogenase | *ALDH9A1* | Amine biosynthetic process | 2,960 |
| Microtubule-associated protein RP/EB family member 2 | *MAPRE2* | Cell cycle | 2,933 |
| Uroporphyrinogen decarboxylase | *UROD* | Cellular metabolic process | 2,927 |
| Coronin-1A | *CORO1A* | Actin cytoskeleton organization | 2,918 |
| Haptoglobin | *HP* | Inflammatory response | 2,859 |
| Collagen alpha-3(VI) chain | *COL6A3* | Axon guidance | 2,814 |
| Argininosuccinate synthase | *ASS1* | Amide biosynthetic process | 2,791 |
| Galectin-3-binding protein | *LGALS3BP* | Biological adhesion | 2,774 |
| von Willebrand factor | *VWF* | Biological adhesion | 2,736 |

In our study, we identified 11 proteins in exosomes that are frequently identified in EVs, of the 100 most frequently reported proteins in Vesiclepedia and ExoCarta databases (Keerthikumar et al., 2016, Kalra et al., 2012) (Table 2, Supplementary Table 3). This indicates the specificity in proteins in exosomes and the existence of common exosomal protein markers across different samples.

**Table 2.** Top 11 proteins identified in this study and present in the Top 100 proteins list of often identified in EVs (in total our study found 11 proteins enriched in exosomes that are referred in the Top 100 list in vesiclepedia (Supplementary Table 3).

| Protein names | Gene names | Biological process | $-\log_{10}(\text{p-value})$ |
| --- | --- | --- | --- |
| Alix | *PDCD6IP* | Cell cycle | 4,836 |
| Adenosylhomocysteinase | *AHCY* | Amine metabolic process | 2,989 |
| Galectin-3-binding protein | *LGALS3BP* | Biological adhesion | 2,774 |
| Syntenin-1 | *SDCBP* | Actin cytoskeleton organization | 2,464 |
| GTP-binding nuclear protein Ran | *RAN* | Actin cytoskeleton organization | 1,877 |
| CD63 antigen | *CD63* | Biological adhesion | 1,589 |
| Talin-1 | *TLN1* | Actin cytoskeleton organization | 1,479 |
| Gelsolin | *GSN* | Actin cytoskeleton organization | 1,477 |
| CD81 antigen | *CD81* | Activation of MAPK activity | 1,454 |
| Complement C3 | *C3* | Activation of immune response | 1,344 |
| Alpha-2-macroglobulin | *A2M* | Biological regulation | 1,112 |

In ectosomes, we detected several proteins commonly identified in larger vesicles, such as plexin-A1 (*PLXA1*), glutathione S-transferase (*GSTP1*), the plasma membrane marker sodium/potassium-transporting ATPase (*ATP1A1*) and epidermal growth factor receptor (*EGFR*) (Figure 3C, Table 3 and Supplementary Table 2) (Zhang et al., 2018, Choi et al., 2013, Choi et al., 2015). Importantly, ADP-ribosylation factor 6 (*ARF6*) is thought to regulate ectosome biogenesis, and was also identified in our dataset (Figure 3C) (Muralidharan-Chari et al., 2009).

**Table 3.** List of top 30 up-regulated proteins in ectosomes identified in this study organized according to the significance values (full list in Supplementary Table 2).

| Protein names | Gene names | Biological process | $-\log_{10}(\text{p-value})$ |
| --- | --- | --- | --- |
| Band 4.1-like protein 2/3 | *EPB41L2/L3* | Actin cytoskeleton organization | 9,852 |
| Plasma membrane calcium-transporting ATPase 1 | *ATP2B1* | Ion transmembrane transport | 7,567 |
| Phosphatidylinositol 4-kinase alpha | *PI4KA* | Actin cytoskeleton organization | 7,239 |
| Tubulin-specific chaperone A | *TBCA* | Cellular component assembly | 6,731 |
| Profilin-1 | *PFN1* | Actin cytoskeleton organization | 6,708 |
| MARCKS-related protein | *MARCKSL1* | Regulation of cell proliferation | 6,644 |
| 4F2 cell-surface antigen heavy chain | *SLC3A2* | Amine transport | 6,412 |
| Cytoplasmic FMR1-interacting protein 1 | *CYFIP1* | Axon extension | 6,209 |
| Myelin protein zero-like protein 1 | *MPZL1* | Cell communication | 6,167 |
| Na(+)/H(+) exchange regulatory cofactor NHE-RF1 | *SLC9A3R1* | Actin cytoskeleton organization | 6,151 |
| Unconventional myosin-1c | *MYO1C* | Cellular component organization | 5,986 |
| Protein scribble homolog | *SCRIB* | Anatomical structure development | 5,985 |
| N(G),N(G)-dimethylarginine dimethylaminohydrolase 1 | *DDAH1* | Amine catabolic process | 5,900 |
| 2,3-cyclic-nucleotide 3-phosphodiesterase | *CNP* | Aging, synaptic transmission | 5,696 |
| Sodium/potassium-transporting ATPase subunit alpha-1 | *ATP1A1* | Cation homeostasis | 5,640 |
| FERM, RhoGEF and pleckstrin domain-containing protein 1 | *FARP1* | Actin cytoskeleton organization | 5,604 |
| Integrin alpha-5 | *ITGA5* | Cell projection organization | 5,589 |
| Catenin delta-1 | *CTNND1* | Axon guidance | 5,556 |
| Charged multivesicular body protein 6 | *CHMP6* | Adherens junction organization | 5,507 |
| Basigin | *BSG* | Cellular component organization | 5,349 |
| PDZ domain-containing protein GIPC1 | *GIPC1* | Anatomical structure morphogenesis | 5,304 |
| Plastin-3 | *PLS3* | Cell communication | 5,244 |
| Peripheral plasma membrane protein CASK | *CASK* | Anatomical structure development | 5,235 |
| Trifunctional purine biosynthetic protein adenosine-3 | *GART* | Calcium ion transmembrane transport | 5,214 |
| 14-3-3 protein epsilon | *YWHAE* | Protein binding | 5,213 |
| Moesin | *MSN* | Cell migration | 5,169 |
| Annexin-A6 | *ANXA6* | Calcium ion binding | 5,150 |
| Myristoylated alanine-rich C-kinase substrate | *MARCKS* | Calcium ion transport | 5,105 |
| Guanine nucleotide-binding protein subunit alpha-13 | *GNA13* | Activation of adenylate cyclase | 5,054 |
| Ubiquitin-40S ribosomal protein S27a | *RPS27A* | Activation of MAPK activity | 5,027 |

Actin, tubulin, and keratins are highly abundant cellular cytoskeletal proteins observed in EVs preparations (Kowal et al., 2016, Mathivanan et al., 2010, Zhang et al., 2018). In our proteomic dataset for ectosomes, we found an enrichment in cytoskeletal proteins as profilin-1 (*PFN1*), cofilin-1/ 2 (*CFL1*/ 2), vimentin (*VIM*), ezrin (*EZR*), moesin (*MSN*), tubulin beta chain (*TUBB*) and actin (*ACTG1*) (Figure 3D, Table 3 and Supplementary Table 2). The absence of other cytoskeletal-associated proteins in our ectosomes and exosomes, such as those involved in actin filament polymerization, associated with microtubules, or intermediate filaments, may be related with their exclusion as potential contaminants in our hit list using Perseus software. The removal of these proteins avoided the identification of proteins that might be present in the cell media but not specifically in our vesicle fractions (Supplementary Table 1 and 2).

Interestingly, our analyses also revealed a significant enrichment in metabolic enzymes in ectosomes, such as peroxiredoxin-1/ 2/ 6 (*PRDX1*/ 2/ 6), pyruvate kinase (*PKM*), alpha-enolase (*ENO1*) and glyceraldehyde-3-phosphate dehydrogenase (*GAPDH*) (Figure 3E, Supplementary Table 2). The presence of annexin family members in EVs fractions has been previously documented and, importantly, perturbations in the levels of their secretion are associated with disease (Popa et al., 2018). Remarkably, we found several annexin proteins enriched in ectosomes, including annexin-A1, A2, A3, A4, A5, A6, A7, A8, A11 (*ANXA1/ 2/ 3/ 4/ 5/ 6/ 7/ 8/ 11*) (Figure 3F, Table 3 and Supplementary Table 2). Integrins have been reported in microvesicles (Hoshino et al., 2015, Muralidharan-Chari et al., 2009) and we found them abundantly in ectosomes [alpha-1/ 2/ 5/ 6 (*ITGA1/ 2/ 5/ 6*)] (Supplementary Figure 1A, Supplementary Table 2). Furthermore, other proteins were enriched in ectosomes as clathrin (*CLTC*), MHC class I (*HLA-A/ B*), cell division control protein 42 (*CDC42*), 14-3-3 proteins (YWHAZ/ E/ B/ G/ H/ Q), histones H1.4 / H1.0/ H2A (*HIST1H1E / H1F0/ HIST2H2AC*), heat shock protein HSP90 (*HSP90AB1/ AA1*), heat shock 70 kDa protein (*HSPA1A/ B, HSPA4*) and heat shock protein 105 kDa (*HSPH1*) (Supplementary Figure 1A, Supplementary Table 2). Matrix metalloproteinase 2 (MMP2) is a protease in the extracellular matrix and it was previously described as an ectosomal marker (Keerthikumar et al., 2015, Taraboletti et al., 2002). However, in our study we did not find an enrichment in matrix metalloproteinases proteins (Supplementary Table 2).

Rabs are small GTPases that regulate numerous vesicle docking and fusion events, including the sorting and trafficking of MVBs to the plasma membrane (van Niel et al., 2018). Rabs form complexes with proteins involved in membrane trafficking through the endocytic system and are usually used as markers of different endocytic compartments. Despite being frequently identified in exosomes, we also identified several Rab proteins in ectosomes (Supplementary Figure 1B, Supplementary Tables 1 and 2). In particular, Rab-1A/ 1B/ 4A/ 5B/ 27B (*RAB1A/ 1B/ 4A/ 5B/ 27B*) were enriched in exosomes, while Rab-2A/ 5C/ 6A/ 6B/ 7A/ 8A/ 8B/ 9A/ 10/ 11B/ 13/ 21/ 23/ 35 (*RAB2A/ 5C/ 6A/ 6B/ 7A/ 8A/ 8B/ 9A/ 10/ 11B/ 13/ 21/ 23/ 35*) were present in ectosomes (Supplementary Figure 1B, Supplementary Tables 1-2). Our findings suggest that Rab proteins may play unique roles in the biogenesis of distinct types of EVs.

We also assessed if the proteins we identified in ectosomes are among the 100 most commonly identified EVs proteins, reported in the Vesiclepedia and ExoCarta databases (Supplementary Table 2 and 3) (Keerthikumar et al., 2016, Pathan et al., 2019). Interestingly, we found 52 proteins in ectosomes that are common to this list (Table 4, Supplementary Table 2). Strikingly, cytoskeleton proteins and cytosolic glycolytic enzymes were absent from our exosomal fraction and were only identified in ectosomes (Supplementary Table 2).

**Table 4.** Top 20 proteins identified in this study and present in the Top 100 proteins list of often identified in EVs (in total our study found 52 proteins enriched in ectosomes that are referred in the Top 100 list in vesiclepedia (Supplementary Table 3).

| Protein names | Gene names | Biological process | $-\log_{10}(\text{p-value})$ |
| --- | --- | --- | --- |
| Profilin-1 | *PFN1* | Actin cytoskeleton organization | 6,708 |
| 4F2 cell-surface antigen heavy chain | *SLC3A2* | Amine transport | 6,412 |
| Sodium/potassium-transporting ATPase subunit alpha-1 | *ATP1A1* | Cation homeostasis | 5,640 |
| 14-3-3 protein epsilon, beta/alpha, theta, zeta/delta | *YWHAE/AB/Q/Z* | Amine metabolic process | 5,213 |
| Moesin | *MSN* | Cell migration | 5,169 |
| Annexin-A6 | *ANXA6* | Calcium ion binding | 5,150 |
| Peroxiredoxin-1 | *PRDX1* | Anatomical structure homeostasis | 4,263 |
| Fatty acid synthase | *FASN* | Acetyl-CoA metabolic process | 4,138 |
| Guanine nucleotide-binding protein G | *GNAS* | Activation of adenylate cyclase | 4,055 |
| Ras GTPase-activating-like protein | *IQGAP1* | Biological regulation | 3,751 |
| Annexin-A2 | *ANXA2* | Calcium binding | 3,628 |
| Integrin beta-1 | *ITGB1* | Actin cytoskeleton organization | 3,579 |
| Ezrin | *EZR* | Actin cytoskeleton organization | 3,375 |
| Heat shock protein HSP 90-beta | *HSP90AB1* | Cellular response to stress | 3,294 |
| Ras-related protein Rab-10 | *RAB10* | Anatomical structure morphogenesis | 3,255 |
| T-complex protein 1 subunit alpha | *TCP1* | Cell recognition | 3,167 |
| Annexin-A1 | *ANXA1* | Innate immune response | 3,162 |
| Ras-related protein Rab-7a | *RAB7A* | Antigen processing and presentation | 3,137 |
| L-lactate dehydrogenase A chain | *LDHA* | Alcohol metabolic process | 2,965 |
| Elongation factor 1-alpha 1 | *EEF1A1* | Biosynthetic process | 2,761 |

In summary, proteomic profiling revealed a diverse protein content in both ectosomes and exosomes but enabled us to identify a unique protein signature that enables their distinction.

### Gene ontology enrichment terms are unique for ectosomes and exosomes

To further understand the distinct biological roles of exosomes and ectosomes, we performed (GO) enrichment analysis using Perseus software (Figures 4 and 5). Signaling processes, immune response, as well as proteasomal and ubiquitination biological processes were enriched in exosomes (Figures 4A). Furthermore, most of the proteins identified were associated with proteasome, organelle and vesicle cellular components, correlating with cytosolic molecular functions of these proteins (Figure 4B and C). Using STRING, we next assessed the physical and functional protein association networks, in order to identify known and predicted protein-protein interactions (Szklarczyk et al., 2019, von Mering et al., 2005). These analyses exhibited a cluster of protein-protein interactions involving proteasomal proteins, as highlighted in the KEGG pathway analyses (Figure 4D and E, Supplementary Figure 2).

**Figure 4.**
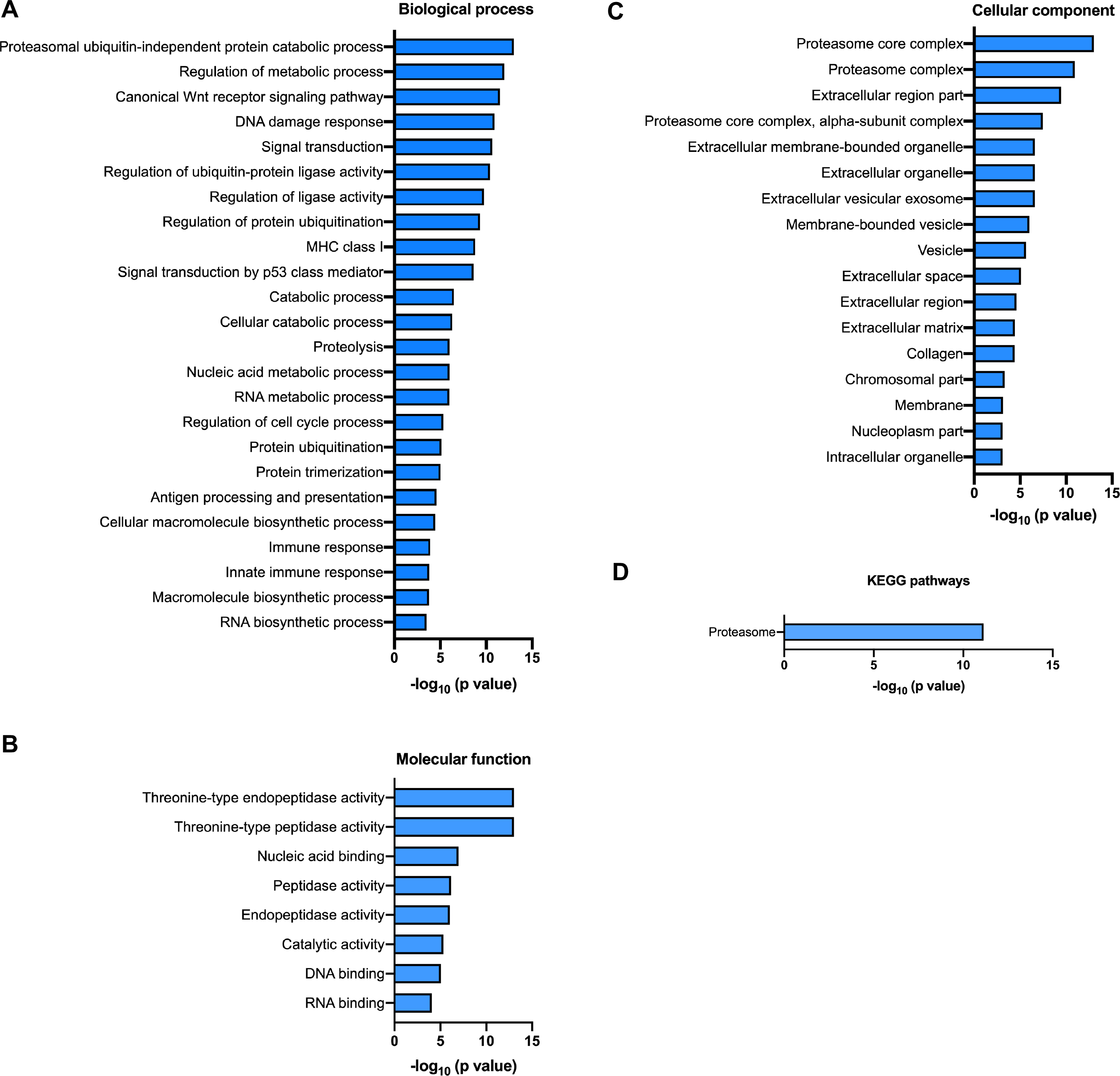
GO terms for proteins enriched in exosomes. The following categories were evaluated: biological process **(A)**, molecular function **(B)**, cellular component **(C)**, and KEGG pathways **(D)**. Data from five independent samples for each group was analyzed using Perseus software. See also Figure S2.

In ectosomes, several of the enriched biological processes were related with the regulation of biological and cellular processes, ion transport, actin regulation related pathways and signal transduction (Figure 5A). As expected, our molecular function hits included protein binding, enzymes and cytoskeletal proteins (Figure 5B). Also, we found a significant enrichment of both glycolysis and gluconeogenesis, key processes of energy metabolism. A large fraction of the proteins identified were associated with plasma membrane, vesicles, cytoplasm and cell interaction and communication processes, thus enabling ectosomes to act as putative intercellular transporters (Figure 5C). Furthermore, KEGG pathway analyses revealed an enrichment of ectosomes in actin cytoskeleton processes and tight junctions, compatible with their plasma membrane origin and enrichment in membrane and cytoplasmic proteins (Figure 5D). STRING association network analyses revealed a large protein-protein interaction network, showing the diversity of proteins enclosed in the vesicles (Supplementary Figure 3).

**Figure 5.**
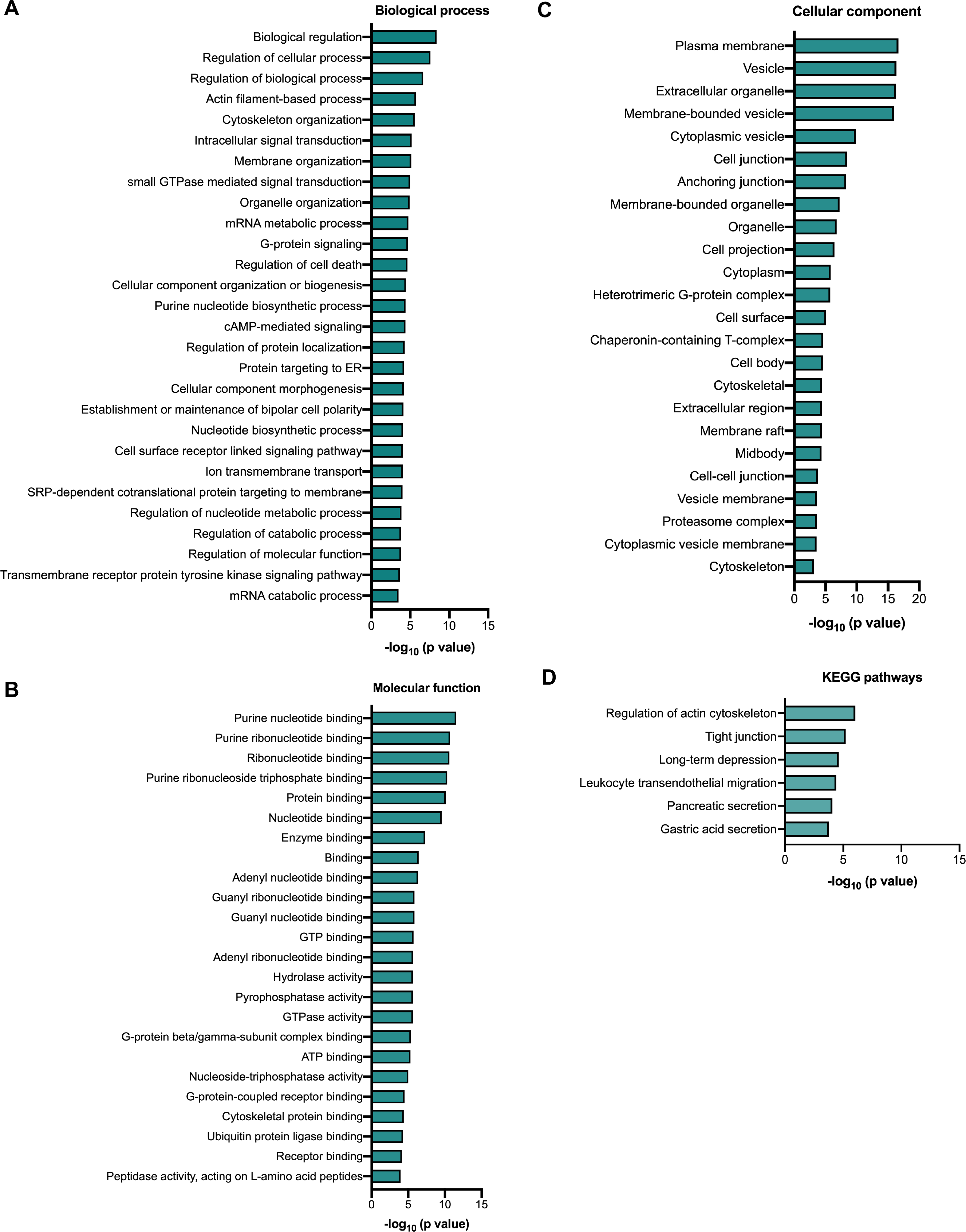
GO terms for proteins enriched in ectosomes. The following categories were evaluated: biological process **(A)**, molecular function **(B)**, cellular component **(C)**, and KEGG pathways **(D)**. Data from five independent samples for each group was analyzed using Perseus software. See also Figure S3.

Overall, these findings suggest that ectosomes may be involved in cell communication, regulating cellular metabolism and organization, and transferring immune or pathological signals between cells.

### Validation of protein markers for ectosomes and exosomes

To confirm the protein markers identified in the proteomic datasets, ectosomes and exosomes fractions were applied into immunoblots and analyzed for several of the protein hits. Staining of ectosomes and exosomes fractions showed a similar total protein profile (Figure 6A, Supplementary Figure 4A). Notably, this was markedly different from what we observed in whole the cell lysates, and we also confirmed that the conditioned media used to collect EVs did not carry residual EVs from the FBS (Supplementary Figure 4B).

**Figure 6.**
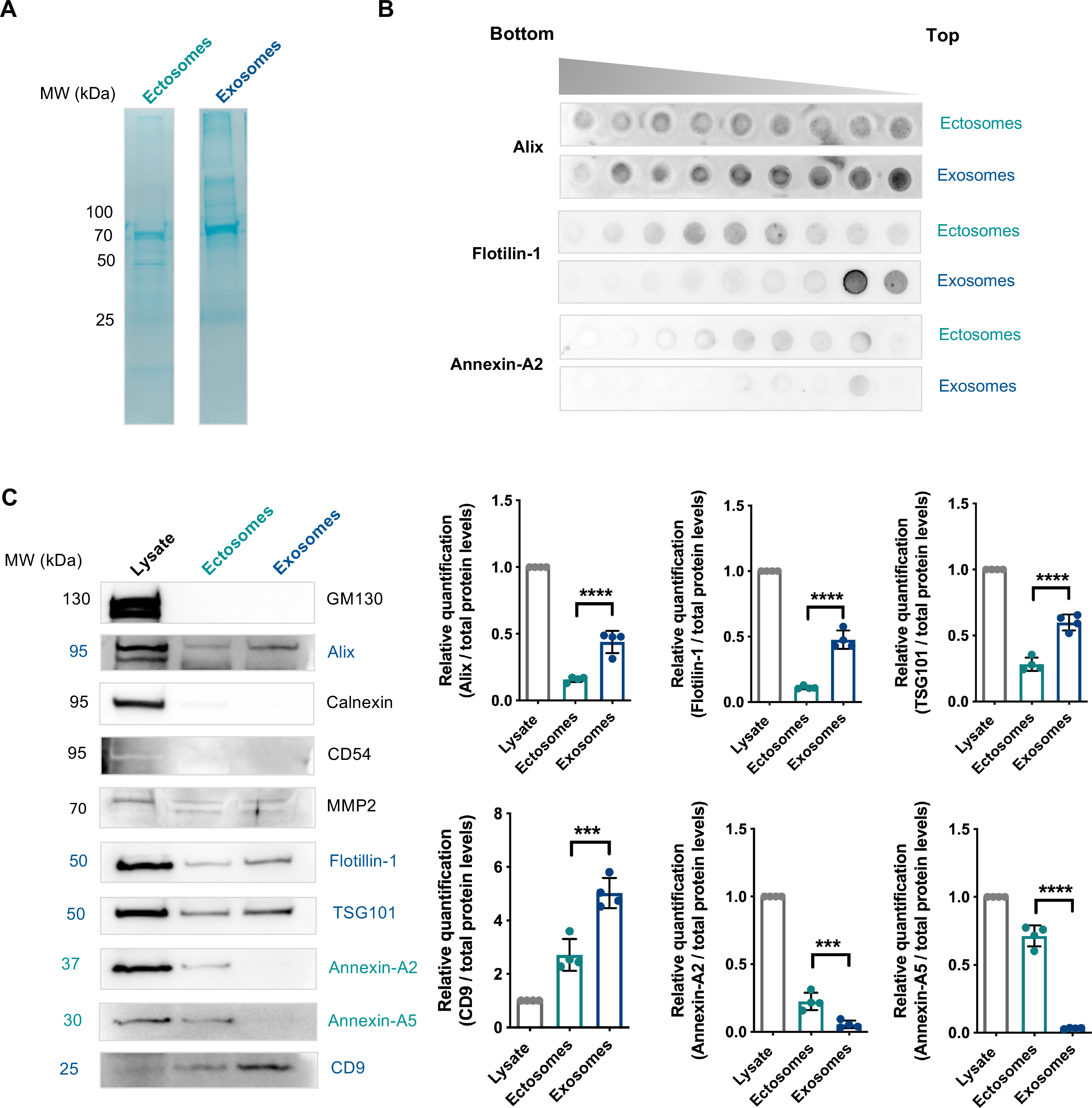
Validation of the proteomic profiling in ectosomes and exosomes. **(A)** MemCode staining demonstrates the total protein levels present in each fraction. **(B)** Ectosomes and exosomes were covered with a discontinuous sucrose step-gradient and fractions were applied into a Dot-Blot system and incubated with antibodies for alix, flotillin-1 and annexin-A2. **(C)** Immunoblots of HEK whole-cell lysates, ectosomes and exosomes fractions. Equal quantities of protein were separated on SDS-PAGE gels, and membranes were blotted with the indicated antibodies. Protein levels were normalized to total protein levels. Data from at least three independent experiments. Significant differences were assessed by one-way ANOVA followed by multiple comparisons with significance between groups corrected by Bonferroni procedure. Differences were considered to be significant for values of p<0.05 and are expressed as mean ± SD, ***p<0.001, ****p<0.0001. See also Figure S4.

In order to evaluate the EVs diversity in each fraction, the vesicles were applied into a sucrose gradient (Figure 6B). This enabled us to separate the EVs according to their floatation speed and equilibrium density, and to uncover the protein markers in each EVs type (Colombo et al., 2014). We found alix was uniformly distributed in the EVs, although in higher levels in exosomes. Interestingly, flotillin-1 was present in higher levels in light exosomes, and was slightly enriched in heavier ectosomes. Interestingly, annexin-A2 was specifically incorporated in lighter ectosomes, and was almost not detected in exosomes (Figure 6B).

Additionally, we assessed the presence of several proteins in the proteomic analyses by immunoblot and compared their distribution among EVs (Figure 6C). Consistent with previous data, alix, flotillin-1, TSG101 and CD9 were significantly enriched in exosomes (Figure 6C) (Thery et al., 2006, Jeppesen et al., 2019, Kowal et al., 2016). Conversely, annexin-A2 and annexin-A5 were significantly increased in ectosomes (Jeppesen et al., 2019) (Figure 6C). MMP2 and CD54 are commonly identified in EVs, but in our samples we did not detect significant differences in their levels in the immunoblot analyses or in the proteomic results (Keerthikumar et al., 2015, Taraboletti et al., 2002, Clayton et al., 2001). Importantly, calnexin and GM130, endoplasmic reticulum and Golgi apparatus markers, respectively, were absent from our preparations, confirming no cross-contamination in the fractions due to cell death (Figure 6C) (Lotvall et al., 2014).

### Markers of cell-derived EVs distinguish ectosomes and exosomes in human CSF

CSF, the fluid bathing the central nervous system, is considered a relevant and accessible window into brain function and homeostasis. Therefore, we next asked whether the markers identified in our cell-derived EVs would enable us to distinguish EVs present in human CSF. Interestingly, ectosomes and exosomes purified from human CSF showed the presence of the same protein markers identified in EVs purified from the cell media (Figure 7). These results indicate that the protocol we used can be applied to different biofluids and confirm that the protein markers we identified are valid for EVs of different origins.

**Figure 7.**
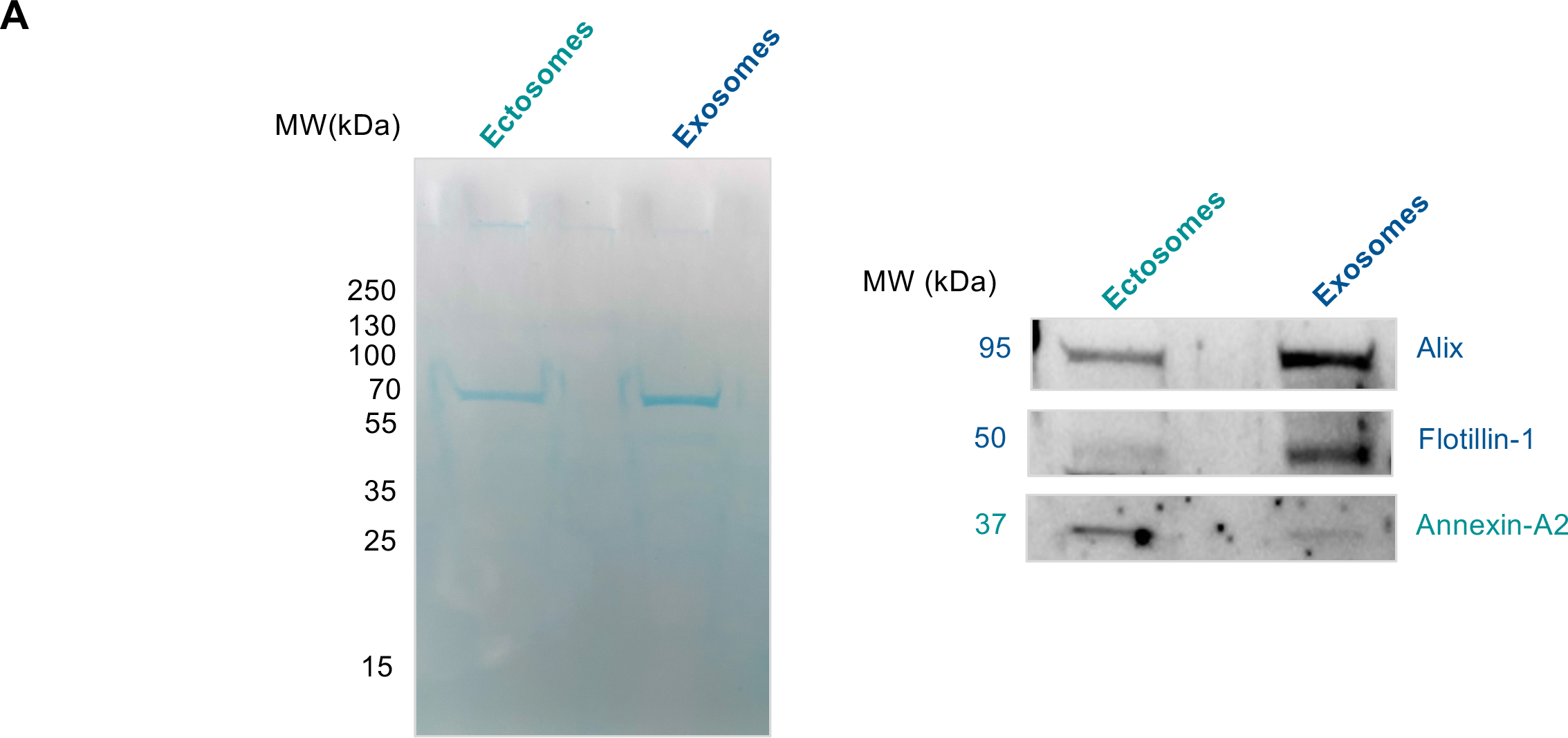
Presence of ectosomal and exosomal markers in human CSF. **(A)** Total protein levels were visualized by staining with MemCode. **(B)** Immunoblot validation of annexin-A2 as ectosomes marker. Alix and flotillin-1 were also evaluated in the blot.

### Purification and visualization of EVs using high-resolution microscopy

Several strategies have been established to enable the visualization of EVs in internalization experiments. These include incubation with dyes that bind to the membrane, or the expression of EVs-related proteins fused to fluorescent tags (Escrevente et al., 2011, Tario et al., 2012, Temchura et al., 2008, Suetsugu et al., 2013, Nakase et al., 2015). However, these strategies present several limitations, such as the leakage of the dye to the plasma membrane after labelled-EVs are internalized, or the incomplete labelling of fluorescent vesicles due to the heterogeneity of protein markers in EVs.

Therefore, to overcome these obstacles, we developed a labelling strategy by stably expressing the green fluorescent protein (EGFP) in HEK cells and purifying ectosomes and exosomes from the cell media. As comparison, we labelled EVs using the thiol-based dye Alexa Fluor 633 C5-maleimide (Roberts-Dalton et al., 2017). As expected, the EGFP signal co-localized with the signal of labelled-EVs (Figure 8A). Interestingly, EGFP-labelled EVs could be imaged at higher resolution when compared with dye-labelled EVs.

**Figure 8.**
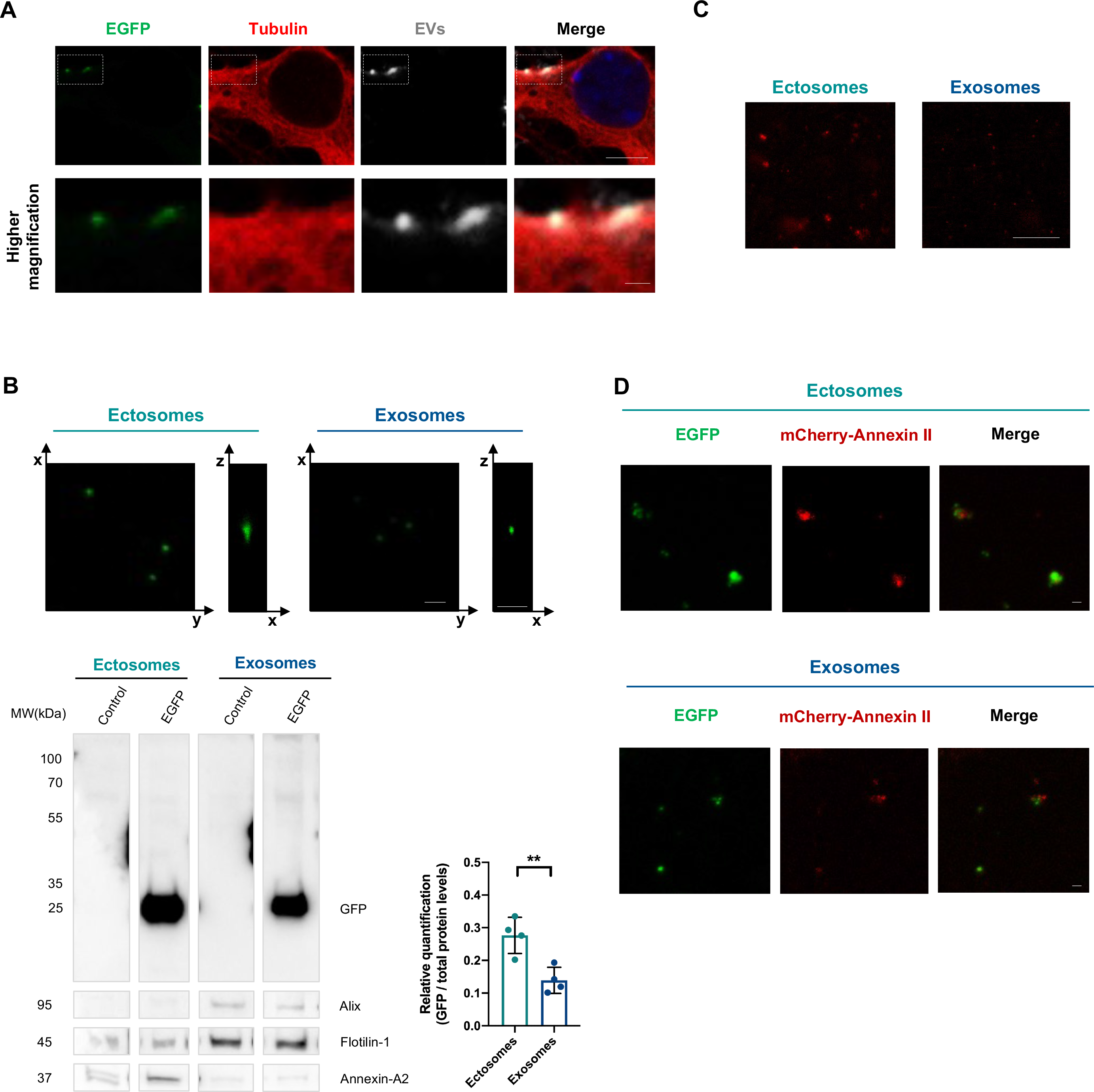
Visualization of ectosomes and exosomes using STED microscopy. **(A)** Representative image of primary cortical neurons treated with exosomes containing EGFP and labelled with thiol-based dye Alexa Fluor 633 C5-maleimide (scale bar 5µm). Higher magnification images of the vesicles on the cell plasma membrane (dashed boxes in white) show the colocalization signal of the double labelling strategies (scale bar 1µm). (**B)** On the top, XY and ZX axis STED imaging of purified ectosomes and exosomes from HEK cells expressing EGFP tag (scale bars represent 1µm in XY and ZX axis). On the bottom, immunoblot of the vesicles displaying the incorporated levels of EGFP and respective vesicle protein markers. The immunoblot lanes represent a montage from the same blot. **(C)** STED imaging of purified ectosomes and exosomes from HEK cells expressing mCherry-Annexin-A2 (scale bar 5µm). **(D)** STED images of purified ectosomes and exosomes containing EGFP and mCherry-Annexin-A2 (scale bar 1µm). See also Figure S5.

When the EGFP positive vesicles were visualized by stimulated emission depletion (STED) microscopy, ectosomes exhibited stronger fluorescence than exosomes (Figure 8B). Additionally, ectosomes incorporated higher levels of EGFP compared with exosomes, independent of their size or total protein levels (quantifications were normalized to total protein levels) (Figure 8B, Supplementary Figure 5A). The higher incorporation of EGFP in ectosomes might be related with its cytosolic expression, making it more available for incorporation into ectosomes. This is consistent with our finding of a greater number of cytoplasmatic proteins in ectosomes than in exosomes.

Next, we further evaluated this labelling strategy and constructed an mCherry-annexin-A2 stable cell line. Ectosomes showed stronger fluorescence intensity for annexin-A2 when compared with exosomes, further validating the enrichment of annexin-A2 in these vesicles (Figure 8C, Supplementary Figure 5B). Additionally, ectosomes purified from cells co-expressing EGFP and mCherry-annexin-A2 exhibited the co-localization of both signals (Figure 8D). mCherry-annexin-A2 was also detected in exosomes using STED microscopy, albeit at lower levels (Figure 8D, Supplementary Figure 5B).

In summary, stable expression of EGFP in cells allows the purification of fluorescently labelled EVs from media of cultured cells for high-resolution imaging studies without changing the levels of specific protein markers.

### Annexin-A2 as specific marker for ectosomes

Annexins are abundant membrane associated proteins that have been identified as EVs constituents (Kowal et al., 2016, van Niel et al., 2018, Zhang et al., 2018, Jeppesen et al., 2019). In our proteomic analyses, we found several annexin proteins enriched in ectosomes, such as annexin-A2 (Figure 3F, Supplementary Table 2). Using sucrose gradients and immunoblot analyses, we observed that annexin-A2 is markedly present in ectosomes (Figure 6B and C). Expression of mCherry-annexin-A2 in cells showed its higher incorporation in ectosomes using STED microscopy (Figure 8C-D, Supplementary Figure 5). Remarkably, ectosomes purified from human CSF also demonstrated an enrichment in annexin-A2 (Figure 7). Together, these results indicate that annexin-A2 is a specific marker of ectosomes from different origins.

### Neuronal cells take up ectosomes and exosomes at similar levels

The uptake of EVs by recipient cells is a key step for intercellular communication. Therefore, we next investigated the internalization and intercellular transfer of both ectosomes and exosomes *in vitro*. Primary cortical neurons were seeded in microfluidic devices in order to separate neuronal cell bodies from their axons and from second-order neurons that would be contacted by those axons (Figure 9A, Supplementary Figure 6A and B). Fluidic isolation was achieved using a volume difference between the soma and axon chambers (Takeda et al., 2015). At DIV14, neurons were treated with an equal amount of exosomes or ectosomes enriched with EGFP (20μg/mL) in the upper left well (input) (Figure 9A, Supplementary Figure 6A and B). Both ectosomes and exosomes were taken up and transferred between neurons. Visualization of the axons in the microgrooves revealed the colocalization of EGFP from the EVs with tubulin staining, demonstrating their internalization and transport in the cells (Figure 9A).

**Figure 9.**
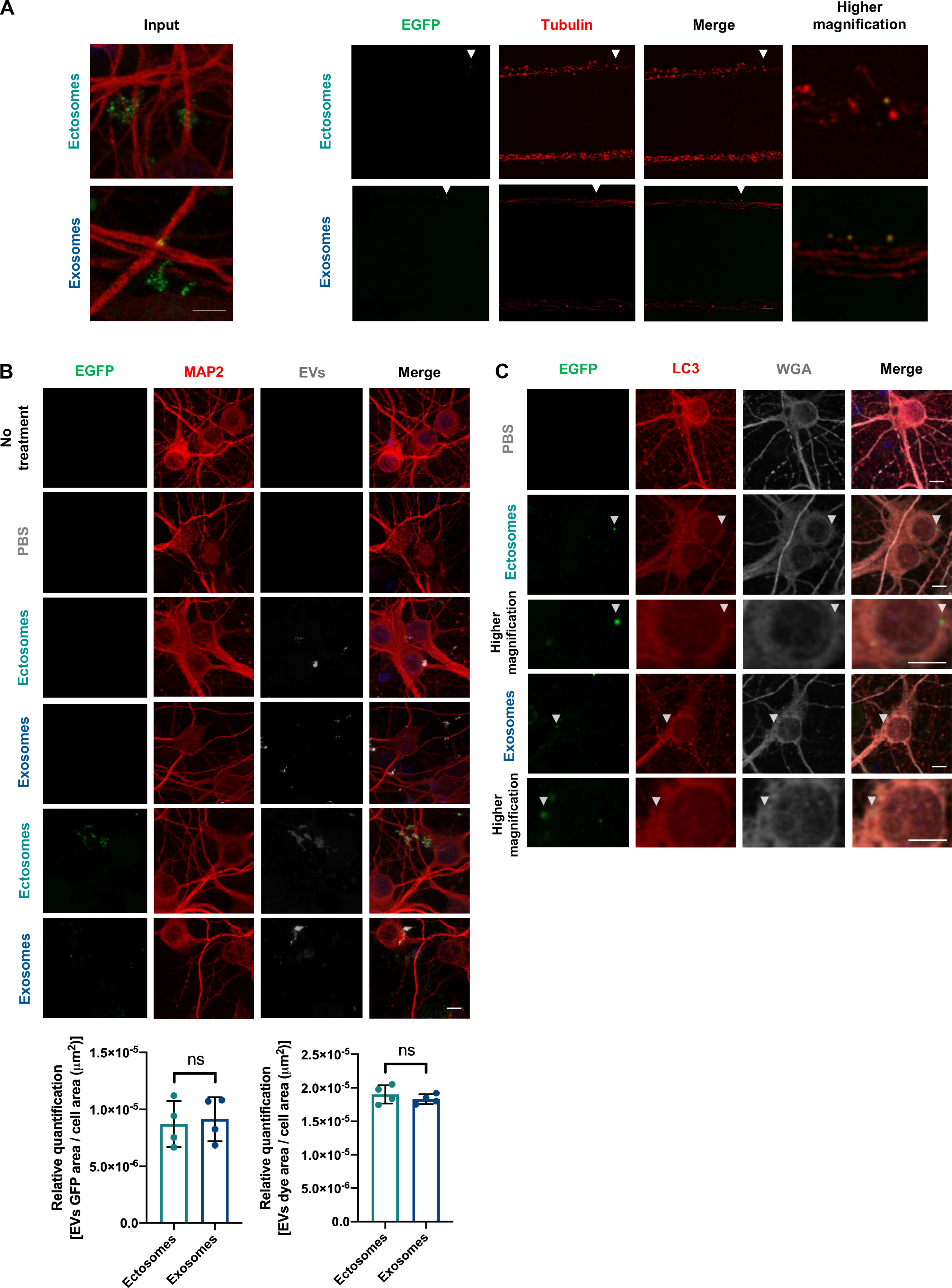
Internalization and axonal transport of EVs by primary cortical neurons. **(A)** Primary cortical neurons were seeded in microfluidic devices to separate neuronal cell bodies. On the left, EVs containing EGFP were added to primary cortical neurons at DIV14 for 24 hours in the upper left input well (20μg/mL). On the right, representative images of cortical neurons immunostained for tubulin demonstrating cytoplasmic internalization and axonal transport of ectosomes and exosomes. White arrowheads show the fluorescent signals within the cell (scale bar 5µm). **(B)** Primary cortical neurons were treated with 20μg/mL EVs at DIV14 for 24 hours. Ectosomes and exosomes without and with EGFP were previously labelled with the fluorescent dye Alexa Fluor 633 C5-maleimide. As a control, dye was incubated with PBS and monitored in the same manner. Representative images of cortical neurons immunostained for MAP2 to demonstrate cytoplasmic internalization of different EVs subtypes (scale bar 5µm). Quantification of the internalized vesicles was performed by measuring the EVs dye or EGFP signal area per cell area internalized by neuronal cells. Data from at least three independent experiments. Significant differences were assessed by two-tailed unpaired t-test comparison and are expressed as mean ± SD. **(C)** Primary cortical neurons were treated with 20μg/mL of EGFP-enriched EVs at DIV14 for 24 hours. Representative images of cortical neurons immunostained for LC3 and cell membrane was labelled using WGA dye. White arrowhead indicates the fluorescent signals within the cell and higher magnification images are presented below each panel (scale bar 5µm). See also Figure S6.

To determine the internalization levels of ectosomes and exosomes in primary cortical neurons, Alexa Fluor 633 C5-maleimide-labelled EVs, with and without EGFP, were added to the medium at DIV14 to a final concentration of 20μg/mL. After 24 hours, cells were fixed, immunostained for MAP2 and analyzed by confocal microscopy. Labelled EVs were observed as puncta predominantly in the cytoplasm of cells, confirming the internalization of ectosomes and exosomes (Figure 9B). PBS-treated neurons did not show positive signal for the dye, indicating that the dye efficiently bound to EVs (Figure 9B). Interestingly, ectosomes and exosomes were internalized at similar levels and the signal of EGFP and dye co-localized for double-labelled EVs (Figure 9B). Additionally, EVs treatment was not toxic to the neurons (Supplementary Figure 6C). Interestingly, after internalization, the EVs-EGFP signal was surrounded by LC3 staining, suggesting a possible engulfment and degradation of EVs in primary cortical neurons (Figure 9C).

### Ectosomes modulate spontaneous activity of cortical neuronal networks

Finally, we assessed the functional relevance of EVs. Since several cell types in the brain actively release EVs to the extracellular space, we hypothesized that neuronal function might be modulated by ectosomes and/or exosomes (Frohlich et al., 2014, You et al., 2020, Sharma et al., 2019, Antonucci et al., 2012). Therefore, we used multi-electrode arrays (MEAs) to evaluate the effect of EVs internalization on spontaneous activity in primary cortical neuronal cultures (Sun et al., 2010). Cells were cultured in MEA chambers until DIV14 to allow the establishment of mature neuronal networks, and spontaneous firing activity was recorded 24 hours after incubation with 20μg/mL of EVs (Figure 10, Supplementary Figure 7). Representative raster plots and voltage traces show the typical firing activity and bursts events in neuronal cultures treated with PBS, ectosomes or exosomes (Figure 10A). Interestingly, assessment of bursting activity parameters showed that under EVs treatment, spikes bursts came more irregularly and showed increased duration (Figure 10B). Exosomes internalization also resulted in longer inter-burst intervals, decreased intra-burst spiking frequency, and a smaller percentage of spikes within the bursts, in contrast to ectosomes (Figure 10B). Although not significant, EVs treatment caused a slight reduction in the burst rate (Supplementary Figure 7C). In our cultures, we also observed a reduction in the mean firing rate after EVs internalization, with decrease of the average spike amplitude for the neurons treated with ectosomes (Supplementary Figure 7D).

**Figure 10.**
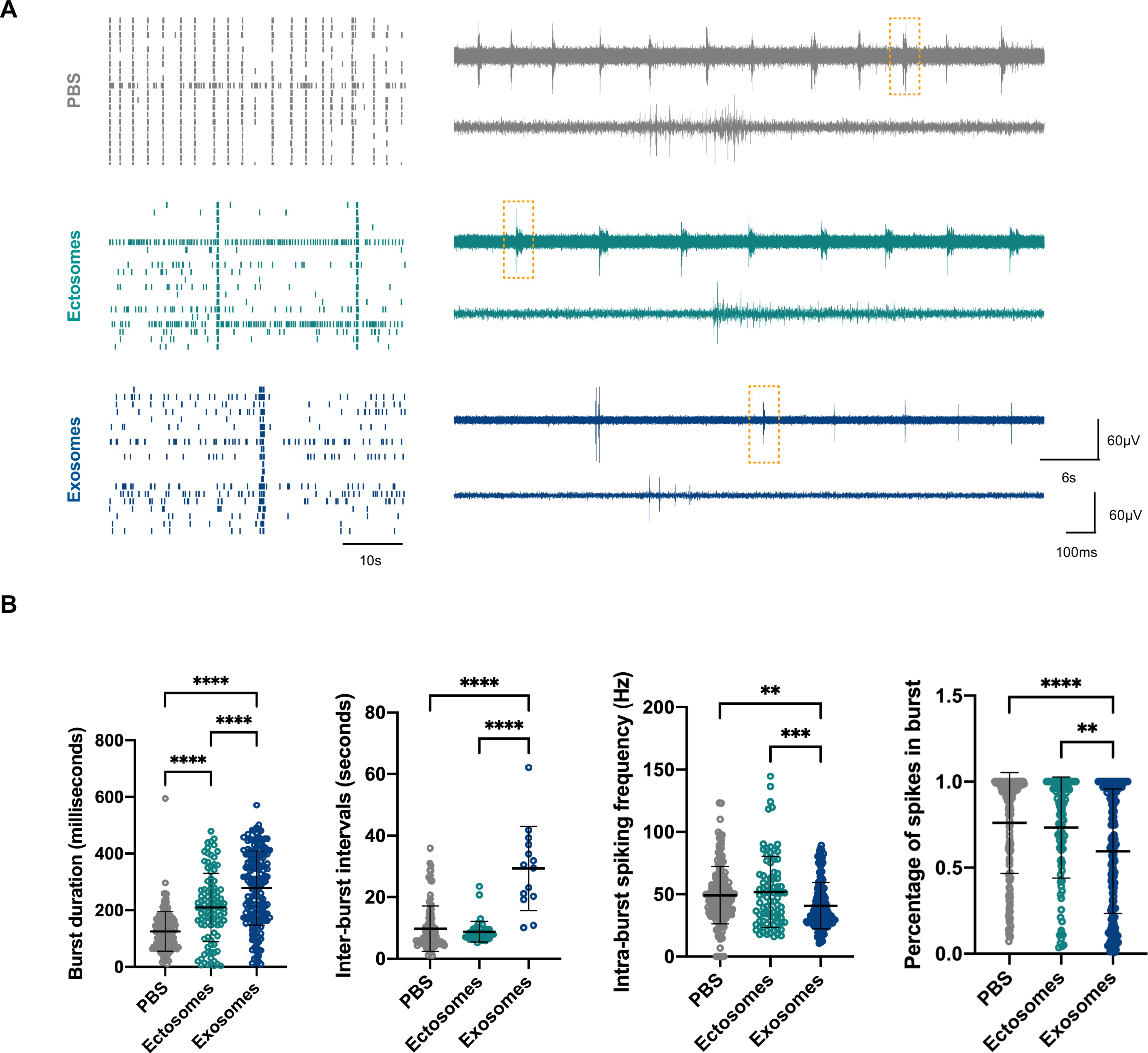
EVs modulate spontaneous activity in primary cortical neurons. **(A)** On the left, representative raster plots of the spontaneous firing activities recorded from cortical neurons after incubation with 20µg/mL of EVs for 24 hours at DIV14, recorded using 60-electrode MEAs. Each row represents one single cell (20 cells shown) and each vertical line represents a single spike obtained on DIV15 [scale bar represents 10 seconds (s)]. On the right, representative voltage traces showing the typical firing activity and bursts events in neuronal cultures treated with PBS, ectosomes or exosomes (upper traces, scale bars represent 60µV and 6s). Closeups of the dashed boxes represent the spikes occurring within a burst ([ower traces, scale bars represent 60µV and 100 milliseconds (ms)]. **(B)** Bursting properties of the cortical neurons treated with EVs (burst duration, inter-burst intervals, intra-burst spiking frequency and percentage of spikes in bursts). Data from at least three independent experiments for each condition. Significant differences were assessed by one-way ANOVA followed by multiple comparisons with significance between groups corrected by Bonferroni procedure. Differences were considered to be significant for values of p<0.05 and are expressed as mean ± SD, **p<0.01, ***p<0.001****, p<0.0001. See also Figure S7.

Altogether, these results suggest that ectosomes and exosomes can modulate important parameters of neuronal spontaneous activity and make the cultured neurons fire in a less synchronized and more irregular fashion.

## Discussion

EVs are important vehicles for remote intercellular communication and signaling. These vesicles encapsulate proteins, RNAs, lipids and signaling molecules that reproduce the cellular content of the origin cell and can modulate several processes in the recipient cells (Basso and Bonetto, 2016, Paolicelli et al., 2019, Hessvik and Llorente, 2018). It is unclear how many different types of EVs are secreted by each cell type. The diversity in their size and content suggests that cells may secrete a large number of different types of vesicles, reflecting distinct physiological roles (Kalra et al., 2016).

The use of EVs as biomarkers has attracted significant interest in several areas of study. However, the use of different EVs purification methods, the absence of reliable markers, and the lack of comprehensive characterization, caused an accumulation of contradictory data and challenges in the study of EVs biology (Raposo and Stoorvogel, 2013, Kowal et al., 2016, van Niel et al., 2018, Zhang et al., 2018). Numerous studies have focused on the characterization and study of the biological function of exosomes. However, ectosomes remain largely understudied (Basso and Bonetto, 2016, Gangoda et al., 2015, Mulcahy et al., 2014, Hessvik and Llorente, 2018). Thus, understanding the role of ectosomes in intercellular communication, the mechanisms involved in their biogenesis, and the characterization of their content will shed light into their biological function and into how they may be used as disease biomarkers.

In our study, we developed a differential ultracentrifugation protocol to efficiently and reproducibly isolate ectosomes and exosomes from diverse biofluids, including human CSF. We observed that ectosomes are larger and are released from cells in higher quantities when compared to exosomes.

Inclusion or exclusion of cellular proteins into EVs appears to be based on controlled protein-sorting mechanisms during their biogenesis, rather than simply on protein abundance in the cell, in agreement with unique protein signatures for each EV subtype. For example, it is known that exosomes are distributed in subpopulations displaying distinct compositions and/or functions (Kowal et al., 2016, Willms et al., 2016). Slight differences in protein content are also expected depending on the isolation protocol and cell type/tissue of origin. Our comprehensive proteomic analyses revealed singular proteomic profiles for ectosomes and exosomes that enabled us to establish protein markers that can be used for their biochemical distinction (Jeppesen et al., 2019, Kowal et al., 2016, Choi et al., 2015). We have identified several membrane-associated proteins, such as annexin-A2 and annexin-A5, as markers for ectosomes. Furthermore, we confirmed that human CSF-derived ectosomes are also enriched in annexin-A2. As previously shown, CD9 and alix are specific markers for cell and CSF-derived exosomes (Kowal et al., 2016, Jeppesen et al., 2019, Choi et al., 2013).

EVs in any biofluid comprise diverse subpopulations that can differ in composition and biogenic mechanisms. Therefore, the variety of machineries involved in the generation of EVs will result in differences in protein composition, lipids and RNA, thereby determining their function and fate. In our study, we demonstrate that ectosomes and exosomes reflect distinct biological processes in the cells. The assessment of ectosomal content highlighted proteins and enzymes involved in processes occurring in the cytosol and plasma membrane. In contrast, exosomal content revealed proteins involved in precise biological processes, such as proteasomal activity. These results indicate that evaluation of different EVs from the same biofluid will provide comprehensive information on biological processes and on the status of the cells. Also, their composition might differ under normal and stress conditions, informing on the signals and cargoes transferred between cells, and reflecting physiological and pathological statuses.

Fluorescent labelling of EVs to investigate their internalization in cells remains challenging due to possible changes in their functionality. The majority of labelling strategies include the use of dyes which bind non-covalently to the membrane bilayer. However, dyes can aggregate or transfer signal to the cell plasma membrane after treatment with labelled-EVs, resulting in misleading results from uptake experiments (Escrevente et al., 2011, Tario et al., 2012, Temchura et al., 2008). Other labelling methods include the use of stable cell lines that fuse GFP to protein markers in EVs (Suetsugu et al., 2013, Nakase et al., 2015). Subsequently, only a subpopulation of EVs will be fluorescent due to EVs heterogeneity. Therefore, we fluorescently labelled EVs through the stable expression of EGFP in cells. EGFP signal co-localized with the signal of labelled-EVs using a thiol-based dye (Roberts-Dalton et al., 2017). Interestingly, the EGFP signal allowed the imaging of EVs internalization with high-resolution, which will be useful for future studies of EVs biology. Furthermore, EGFP is more incorporated in ectosomes than in exosomes, likely as a result of EGFP availability in the cytoplasm and due to the distinct origins of the EVs. We also observed that stable expression of mCherry-annexin-A2 resulted in its incorporation in higher levels in ectosomes than in exosomes, further confirming the use of annexin-A2 as an ectosomal marker. In summary, in our study we developed a simple and practical tool to fluorescently label EVs derived from cultured cells. This strategy takes advantage of the cytosolic expression of EGFP, without compromising the protein composition of the EVs.

After the detailed biochemical characterization of both ectosomes and exosomes, we evaluated their internalization and functional implications in neuronal activity by treating primary cortical neurons with the different EVs. Neurons take up and transport both ectosomes and exosomes between cells and do not display any detectable signs of neurotoxic effects. The LC3 staining surrounding the EVs after internalization suggests at least a fraction enters in the cells as intact vesicles and are sorted for degradation by autophagy.

Ectosomes and exosomes carry a complex assortment of biomolecules with distinct activities that become internalized by receptor cells, most probably producing distinct functional outputs. EVs have been hypothesized to play important roles in the nervous system (van Niel et al., 2018). However, the potential role of ectosomes and exosomes in neural function is unclear. Assessment of neuronal network activity using MEA recordings demonstrated that spontaneous neuronal activity can be modulated by ectosomes and exosomes. Most importantly, EVs treatment disrupted the regular synchronized bursting activity neuronal cultures, resulting in overall lower and more disorganized spiking activity. We hypothesize that this reorganization of concerted neuronal activity might result from EVs acting on the presynaptic site, where they may decrease the release of synaptic vesicles. Previous studies showed that exosomes purified from microglial and oligodendroglial cells can also modulate neuronal activity (Frohlich et al., 2014, Antonucci et al., 2012). However, the impact of EVs on neuronal excitability *in vivo* remains understudied.

Altogether, our study forms the foundation for future studies of EVs biology, demonstrates distinct biogenesis and, importantly, confirms distinct functional effects of exosomes and ectosomes in neuronal networks *in vitro*. Moreover, our work will aid efforts to discover future biomarkers for different human pathological conditions.

## Acknowledgments

We thank Sabine König and Uwe Plessmann from Max Planck Institute for Biophysical Chemistry (Göttingen) and Christof Lenz from the Core Facility Proteomics at University Medical Center Göttingen (Göttingen) for helping with mass spectrometry analysis. TFO is supported by European Union’s Horizon 2020 research and innovation program under grant agreement No. 721802 (SynDegen), and by the Deutsche Forschungsgemeinschaft (DFG, German Research Foundation) under Germany’s Excellence Strategy - EXC 2067/1-390729940, by SFB1286 (B8). TG is supported by the European Research Council (ERC) under the European Union’s Horizon 2020 research and innovation programme (grant agreement number 724822). This work was partly supported by the Göttingen Graduate School for Neurosciences, Biophysics, and Molecular Biosciences (DFG grant GSC 226/4). Annexin-A2-GFP plasmid was kindly provided by Volker Gerke & Ursula Rescher (Plasmid #107196, Addgene) (Rescher, Zobiack et al. 2000). Figures were created with BioRender.com.

## Author contributions

T.F.O. and I.C.B. conceived the study. I.C.B performed all the cell culture, molecular biology and imaging experiments. M.H.K and I.C.B stablished the spike sorting framework, performed MEA experiments and data analysis. D.R performed the EM experiments. I.P. evaluated the mass spectrometry data using MaxQuant. E.G prepared the lentiviral vectors used in the study. C.V.R provided the CSF material for the study. T.G. provided methodology and resources for the MEA experiments. I.C.B analyzed and interpreted the data. I.C.B generated the graphs and figures. I.C.B. and T.F.O. wrote the manuscript. T.F.O. supervised the work.

## Methods

### Resource availability

#### Lead contact for reagent and resource sharing

Further information and requests for resources and reagents should be directed to and will be fulfilled by the Lead Contact, Tiago F. Outeiro.

#### Data and code availability

Original data generated by this study are available upon request.

### Experimental model and subject details

#### Human cerebrospinal fluid samples

Human cerebrospinal fluid (CSF) samples were obtained with informed consent from adult volunteers and all experiments were performed in accordance with relevant guidelines and regulations. Samples were obtained from the CSF bank of the University Medical Center Göttingen (Göttingen, Germany). Samples were classified according to the disease diagnostics and information about sex and age were anonymized. CSF samples were stored in0.5, 1.0 and 1.5mL aliquots at -80°C prior to analysis and were pooled for extracellular vesicles (EVs) isolation (10mL final volume).

#### Neuronal culture

Primary cortical neuronal cultures were prepared from C57BL6/J#00245 wild-type E15.5 embryos from the animal facility of the University Medical Center Göttingen (Göttingen, Germany), as previously described (Szego et al., 2019). C57BL6/J#00245 background mice were bred and maintained under specific pathogen free conditions in the animal facility of the University Medical Center Göttingen (Göttingen, Germany). Animal procedures were performed in accordance with the European Community (Directive 2010/63/EU), institutional and national guidelines (license number 19.3213). In detail, pregnant animals were sacrificed by carbon dioxide intoxication and the embryos extracted from the uterus. The meninges were removed, and the cortex was dissected under a stereomicroscope, and afterwards the tissue was transferred to ice-cold 1x Hanks balanced salt solution (CaCl_2_ and MgCl_2_ free; HBSS) (Gibco Invitrogen, CA, USA) supplemented with 0.5% sodium bicarbonate solution (Sigma-Aldrich, MO, USA). After trypsinization at 37°C for 15 minutes (min) (100μL of 0.25% trypsin; Gibco Invitrogen, CA, USA), the reaction was stopped by adding 100μL DNase I (0.5mg/mL, Roche, Basel, Switzerland) and 100μL fetal bovine serum (FBS) (Anprotec, Bruckberg, Germany). The tissue was gently dissociated using a glass pipette. After centrifugation at 300x*g* for 5 min, cells were resuspended in pre-warmed neurobasal medium (Gibco Invitrogen, CA, USA) supplemented with 2% B27 supplement (Gibco Invitrogen, CA, USA), 1% penicillin-streptomycin (PAN Biotech, Aidenbach, Germany) and 0.25% GlutaMAX (Gibco Invitrogen, CA, USA). Cells were seeded on coverslips coated with poly-L-ornithine (0.1mg/mL in borate buffer; PLO) (Sigma-Aldrich, MO, USA) or culture plates (Corning, Merck, Darmstadt, Germany) for immunocytochemistry and western blot experiments. Cells were maintained at 37°C with 5% CO^2^, and fresh medium was added every 3-4 days.

#### Human embryonic kidney cells

Human embryonic kidney (HEK) 293 cells (ATTC, VA, USA) were maintained in DMEM medium (PAN Biotech, Aidenbach, Germany) supplemented with 10% FBS (Anprotec, Bruckberg, Germany) and 1% penicillin-streptomycin (PAN Biotech, Aidenbach, Germany) at 37°C in a 5% CO^2^ atmosphere. Stable cell lines expressing pRRL-CMV-EGFP-PRE-SIN, FUGW-mCherry-Annexin-A2 or FUGW-mCherry were developed by lentiviral infection of HEK 293 cells. Cells were incubated during 5 days with the virus and after three passages the infection rate was confirmed by microscopy (more than 90% of positive cells).

### Method details

#### Lentivirus production protocol

Production of lentivirus was performed as previously described (Tiscornia et al., 2006). HEK

293 cells were seeded in culture plates (Corning, Merck, Darmstadt, Germany) and incubated overnight at 37°C with 5% CO^2^ in DMEM (PAN Biotech, Aidenbach, Germany) supplemented with 10% FBS (Anprotec, Bruckberg, Germany) and 1% penicillin-streptomycin (PAN Biotech, Aidenbach, Germany). On the next day, cells were incubated with transfection medium containing DMEM with 2% FBS (Anprotec, Bruckberg, Germany) for 5 hours (h) before transfection. Calcium phosphate (CaPO4) precipitation method was used for transfection with a plasmid mix [144μg of Delta 8.9 packaging virus, 57.9μg vesicular stomatitis virus glycoprotein (VSVG) packing virus and 160μg of the plasmid of interest]. DNA mix was added to 6 mL of 1x BBS (50 mM BES, 280 mM NaCl, 1.5 mM Na2HPO4) and 0.36 mL CaCl_2_ (2.5M CaCl_2_) was added to the mixture in a vortex shaker. Solution was incubated 20min in the dark before adding to the cells. On the next day, the medium was changed to Panserin (PAN Biotech, Aidenbach, Germany) supplemented with 1% of non-essential amino acids (MEM, Gibco Invitrogen, CA, USA) and 1% penicillin-streptomycin (PAN Biotech, Aidenbach, Germany). Viruses were harvested 72h after transfection, centrifuged at 3000x*g* for 15min at 4°C. The supernatant was cleared of cell debris by filtering through a 0.45µm filter (Corning, Merck, Darmstadt, Germany) and mixed with 1x PEG solution (SBI System Bioscience, CA, USA) to pellet the virus. After 2 days of incubation at 4°C, viruses were spin down by centrifugation at 1500x*g* during 30min at 4 °C. Supernatant was discarded and the pellet was resuspended in Panserin (PAN Biotech, Aidenbach, Germany).

#### Microfluidic chambers

Triple compartment neuron silicone devices (TCND1000) with 2 microgroove barriers of 500μm with a 1000μm central compartment were obtained from Xona microfluidics and prepared for neuronal cell culture as previously described (Harris et al., 2007, Takeda et al., 2015) and following the manufacturer’s instructions (Xona microfluidics, NC, USA). Experiments were performed as previously described (Freundt et al., 2012). Cover glass 24mm x 40mm (Corning, Merck, Darmstadt, Germany) and microfluidic devices were rinsed with 70% ethanol (Sigma-Aldrich, MO, USA) and water under sterile conditions. Cover glass were coated with poly-L-lysine (0.5 mg/ mL in borate buffer; PLL) (Sigma-Aldrich, MO, USA). After bonding the microfluidic devices to glass coverslips, neurons were seeded in the soma channels and incubated at 37°C, 5% CO^2^, in a humidified incubator. Furthermore, the second and third chambers were filled with more neuronal cell media than in the first chambers. The volume difference between the chambers resulted in continuous hydrostatic pressure barrier, which also prevented diffusion of EVs from the treated chamber into others. Treatment was performed at days *in vitro* (DIV) 14 and the cells fixed for immunocytochemistry at DIV15 (see immunocytochemistry section).

#### Immunocytochemistry experiments

After treatment, primary and cell lines cultures were first washed with 1x PBS (PAN Biotech, Aidenbach, Germany) and fixed with 4% of paraformaldehyde solution (PFA) for 20min at room temperature (RT). In order to quench PFA autofluorescence, samples were incubated with 50 mM of ammonium chloride (NH4Cl) solution for 30min. In order to label the plasma membrane, cells were incubated during 10 min with wheat germ agglutinin (WGA) Alexa FluorTM 633 conjugate in 1x HBSS at RT (5.0 µg/mL; Invitrogen, CA, USA). Cells were washed with 1x PBS (PAN Biotech, Aidenbach, Germany). Cells were permeabilized with 0.1% Triton X-100 (Sigma-Aldrich, MO, USA) for 10min. Following permeabilization, cells were blocked with 2% bovine serum albumin (BSA) in 1x PBS (Sigma-Aldrich, MO, USA) for 1h at RT and then incubated with the primary antibodies overnight at 4°C. Afterwards, the cells were washed with 1x PBS (PAN Biotech, Aidenbach, Germany) and then with fluorescence conjugated secondary antibodies for 2h at RT. Finally, nuclei were counter-stained with DAPI (Carl Roth, Karlsruhe, Germany) and mounted with mowiol for microscopy.

#### Western blots

Cultured cells were collected and lysed in RIPA buffer (50 mM Tris, pH 8.0, 0.15 M NaCl, 0.1% SDS, 1.0% NP-40, 0.5% Na-Deoxycholate, 2mM EDTA, supplemented with protease and phosphatase inhibitors cocktail (cOmplete^TM^ protease inhibitor and PhosSTOP^TM^ phosphatase inhibitor; Roche, Basel, Switzerland). Protein concentrations in the lysates were determined by the Bradford protein assay (Bio-Rad, CA, USA). 30µg of protein were denaturated 5min at 95°C, loaded into 12% SDS-PAGE gels and transferred to nitrocellulose membranes using iBlot2 (Invitrogen, CA, USA). Membranes were blocked with 5% skim milk (Sigma-Aldrich, MO, USA) in Tris-buffered saline (pH 8) with 0.05% Tween 20 (TBS-T) and then incubated with the appropriate primary antibody overnight in 5% BSA (Sigma-Aldrich, MO, USA) in TBS-T at 4°C. After three washes with TBS-T, membranes were incubated for 2h with horseradish peroxidase (HRP) conjugated secondary antibodies. After incubation with the secondary antibody, membranes were washed three times with TBS-T and developed in a chemiluminescence system (Fusion FX Vilber Lourmat, Vilber, France) using chemiluminescent HRP substrate (Millipore, MA, USA). Intensities of specific bands were normalized to a protein loading control or to the total protein levels marked using MemCode™ Reversible Protein (Thermo Fisher Scientific, MA, USA).

#### Isolation of extracellular vesicles

Isolation of ectosomes and exosomes was performed using an adapted protocol from previous studies (Thery et al., 2006, Dujardin et al., 2014). HEK 293 cells were grown in conditioned medium (depleted of FBS-derived exosomes), produced as previously described (Thery et al., 2006). Briefly, DMEM (PAN Biotech, Aidenbach, Germany) supplemented with 20% FBS (Anprotec, Bruckberg, Germany) and 2% penicillin-streptomycin (PAN Biotech, Aidenbach, Germany) was centrifuged in polypropylene tubes (Optiseal; Beckman Coulter, CA, USA) in a fixed rotor (type 70 Ti; Beckman Coulter, CA, USA) during 16h at 100 000x*g*, 4**°**C. The depleted medium was then diluted with DMEM medium (PAN Biotech, Aidenbach, Germany) in order to reach the final supplements concentration required to make the conditioned medium. The cells were seeded in T75 cm^2^ flasks (Corning, Merck, Darmstadt, Germany) and grew until 80% confluency, then were washed with warm 1x PBS (PAN Biotech, Aidenbach, Germany) and incubated with 14mL of fresh conditioned media during 24h. The media was collected and placed on ice before centrifuging for 10min at 300x*g* at 4**°**C to pellet cell debris. The supernatant was collected for new tubes and frozen in 13mL aliquots at -80**°**C for further analyses. To isolate EVs, the supernatant was thawed in ice and centrifuged 20min at 2 000x*g*, 4**°**C. All the isolation protocol was performed in the cold room at 4**°**C and the samples were maintained in ice. 12mL of the supernatant was transferred into ultra-clear tubes (Beckman Coulter, CA, USA) and centrifuged in a swing rotor (TH-641 Sorvall; Thermo Fisher Scientific, MA, USA) during 90min at 20 000x*g*, 4**°**C. 11mL of the medium was carefully transferred into a new centrifuge tube with a sterile pipette and the pellet containing ectosomes was resuspended in ice cold 1x PBS (PAN Biotech, Aidenbach, Germany). The medium was again centrifuged in a swing rotor (TH-641 Sorvall; Thermo Fisher Scientific, MA, USA) during 90min at 100 000x*g* (4**°**C) to generate exosomes. The supernatant was discarded and the pellet containing exosomes was resuspended in ice cold 1x PBS (PAN Biotech, Aidenbach, Germany) with protease and phosphatase inhibitors (cOmplete^TM^ protease inhibitor and PhosSTOP^TM^ phosphatase inhibitor; Roche, Basel, Switzerland). Both ectosomes and exosomes pellets were again centrifuged at the correspondent velocities in order to concentrate and clean the pellets. Afterwards the supernatant was removed completely by inverting the tubes and the pellets were resuspended in 100µl of 1x PBS (PAN Biotech, Aidenbach, Germany). Protein concentrations were determined by the BCA Protein assay (Thermo Fisher Scientific, MA, USA).

#### Labelling of the EVs

Labelling of EVs was performed as previously described (Roberts-Dalton et al., 2017). Briefly, Alexa Fluor 633 C5-maleimide (200μg/mL; Invitrogen, Carlsbad, California, CA, USA) was added to an aliquot of EVs purified after ultracentrifugation containing 60-100µg of total protein for a final volume of 50μL in 1x PBS (PAN Biotech, Aidenbach, Germany). Samples were incubated for 1h with no agitation in the dark at RT. Non-incorporated and excess of dye was removed by washing the sample in 1x PBS (PAN Biotech, Aidenbach, Germany) and centrifugation during 90min at 20 000x*g* (4**°**C) for ectosomes and at 100 000x*g* for 90min for exosomes (4**°**C) in a swing rotor (TH-641 Sorvall; Thermo Fisher Scientific, MA, USA). As a control, 1x PBS (PAN Biotech, Aidenbach, Germany) without EVs was also incubated with the dye and performed in parallel to confirm dye removal by washing with 1x PBS (PAN Biotech, Aidenbach, Germany) and ultracentrifugation. For microscopy analysis, labelled EVs were gently mixed with mowiol and applied into PLO-coated coverslips and glass slides.

#### Sucrose gradient

Sucrose gradient centrifugation was performed from crude vesicle pellets adapted from previous studies (Chiou, 2016, Thery et al., 2006). Sucrose stock solutions (10, 16, 22, 28, 34, 40, 46, 52, 58, 64, 70 and 90%) (Sigma-Aldrich, MO, USA) were prepared in 1x PBS (PAN Biotech, Aidenbach, Germany). The crude exosome and ectosome preparations (in 100μL PBS) were resuspended in 1mL of 90% sucrose stock solution (82% final sucrose concentration). Each sample was transferred to an ultra-clear centrifuge tube (Beckman Coulter, CA, USA) and overlayed with the rest of the gradient stocks on top starting with 1mL of 70% sucrose solution and the last with 10% solution, in order to make a gradient going from the highest to lowest sucrose concentration. Samples were centrifuged for 16h at 100 000x*g* at 4°C (TH-641 Sorvall; Thermo Fisher Scientific, MA, USA) and afterwards 1mL fractions were collected starting from the top to bottom. Fractions were resuspended in 1x PBS (PAN Biotech, Aidenbach, Germany) and centrifuged 100 000x*g* at 4°C for 90min (TH-641 Sorvall; Thermo Fisher Scientific, MA, USA). The final pellets were resuspended in 1x PBS (PAN Biotech, Aidenbach, Germany) and applied into a DotBlot system with a 0.2µm nitrocellulose membrane (Bio-Rad, CA, USA).

#### NTA analysis

Particle number and size distribution in ectosomes and exosomes samples were determined by nanoparticle tracking analysis (NTA) using a NanoSight LM10 instrument and a LM14 viewing unit equipped with a 532 nm laser (NanoSight, Salisbury, UK). Total EVs samples were diluted in 1x PBS (PAN Biotech, Aidenbach, Germany) to a final volume of 400mL prior to analysis, according to the manufacturer recommendations. Data was recorded using NTA 2.3 software with a detection threshold of 5, captured with a camera level of 16 at 25°C, in videos of 5 x 60 seconds (s) repeated 4 times and averaged for each biological replicate.

#### Electron microscopy

Electron microscopy images from ectosomes and exosomes was performed following a protocol previously described (Thery et al., 2006). Isolated EVs were bound to a glow discharged carbon foil covered grids. After staining with 1% uranyl acetate_*(aq.)*_, the samples were evaluated at RT with a Talos L120C transmission electron microscope (Thermo Fisher Scientific, MA, USA).

#### Proteomic analyses of EVs

Samples were resuspended in Laemmli sample buffer and separated by SDS-PAGE. The entire lane was subsequently cut in 23 gel pieces and tryptically digested, as previously described (Shevchenko et al., 2006). Peptides extracted from the gel pieces were analyzed in technical triplicates by liquid chromatography coupled to mass spectrometry (LC-MS) on a Dionex UltiMate 3000 RSLCnano system connected to an Orbitrap Fusion mass spectrometer (Thermo Fisher Scientific, MA, USA). Peptides were separated by a 43min gradient ranging from 8% to 37% acetonitrile on an in-house packed C18 column (75µm x 30cm, Reprosil-Pur 120C18-AQ, 1.9µm, Dr. Maisch GmbH, Ammerbuch, Germany) at 300 nl/min flow rate. MS1 spectra were acquired with 120 000 resolution (full width at half maximum, FWHM) and a scan range from 350 to 1,600 m/z. Within a cycle time of 3s, precursor ions with a charge state between +2 and +7 were selected individually with a 1.6 m/z isolation window and were fragmented with a normalized collision energy of 35 by higher energy collisional dissociation (HCD). MS2 spectra were acquired in the ion trap with 20% normalized AGC and dynamic injection time. Once selected precursors were excluded from another fragmentation event for 30s. Raw acquisition files were subjected to database search with Maxquant (version 1.6.2.10) (Cox and Mann, 2008) against the reference proteome of *Homo sapiens* (downloaded on 19/2019). Default settings were used unless stated differently below. Fractions were defined according to the cutting of the gel lanes and experiments were defined on the level of technical replicates. Unique and razor peptides were used for label-free quantification.

Data analysis was performed with Perseus (version and 1.6.15.0) (Tyanova et al., 2016). Reverse hits, potential contaminants and hits only identified by site were filtered out. Quantitative values were averaged across technical replicates ignoring missing values. A two-sample t-test was performed on biological replicates of ectosomes and exosomes with an artificial within groups variance (s0) of 0.1 and a permutation-based FDR of 0.1 for multiple testing correction. Results were visualized by volcano plots. To assess the reproducibility of the experiments within the biological replicates of each cell extracellular vesicle type, we employed principal component analysis (PCA), which was performed using the Perseus built-in tool. The EVs subtype-signature proteins were selected by extracting in Perseus the proteins with the most distinct expression profiles between the different subtypes. For hierarchical clustering, normalized intensities were first z-scored and then clustered using Euclidean as a distance measure for column and row clustering. GO enrichment analyses were performed using Perseus software and the human proteome as background (”mainAnnot.homosapiens.txt.gz, downloaded in 04/ 2021). Enrichment was considered statistically significant using Fisher exact test and corrected for multiple testing by the Benjamini-Hochberg FDR method with adjusted 0.02 threshold value (GO terms were plotted as -log10 of the p value). Gene ontology analyses were performed using the hit proteins identified in each EVs type.

#### Internalization experiment with EVs

Primary cortical neurons were treated with 20μg/mL of ectosomes or exosomes resuspended in 1x PBS (Sigma-Aldrich, MO, USA). EVs were added to cortical neurons at DIV14 and collected for further analyses at DIV15, 24h after the treatment.

#### Lactate DeHydrogenase assay

Cytotoxicity in primary and cell line cultures was assessed using the cytotoxicity Lactate DeHydrogenase (LDH) detection kit according to the manufacturer’s instructions (Roche, Basel, Switzerland). The culture medium was centrifuged at 500x*g* for 5min to pellet cell debris before used in the experiments.

#### Multi-electrode array

Multi-electrode array (MEA) experiments were performed following standard protocols for MEA recordings (Wagenaar et al., 2004, Khani and Gollisch, 2021, Frohlich et al., 2014, You et al., 2020, Sharma et al., 2019, Antonucci et al., 2012). Primary cortical neuronal cultures were plated directly on 60MEA200/30iR-Ti-gr planar arrays (60 electrodes, 30µm electrode diameter and 200µm electrode spacing; MultiChannel Systems, Reutlingen, Germany). The arrays were coated with PLL (500µg/mL in borate buffer; Sigma-Aldrich, MO, USA) overnight at 4°C. The arrays were rinsed three times with distilled water before coating with laminin (5µg/mL in distilled water; Sigma-Aldrich, MO, USA) for at least 1h at RT. The solution was aspirated, and the cells were directly plated on top of the electrodes. Neurons were treated with 20µg/mL of EVs at DIV14 and recorded at DIV15, 24h after the treatment. The neuronal activity over the array was recorded using the MultiChannel MEA2100 system (MultiChannel Systems, Reutlingen, Germany) and the cells were kept in their culture medium with temperature maintained at 35-37°C during recordings. To avoid movement-induced artifacts, recordings started 10min after translocation of arrays from the incubator to the recording stage. The spontaneous activity of the cultured neurons was recorded for 5 - 10 min. The electrode signals were amplified, band-pass filtered (200 Hz to 3 kHz) and recorded digitally at 25 kHz, using the MultiChannel Experimenter software (version 2.17.7.0, MultiChannel Systems, Reutlingen, Germany).

Spike sorting was carried out using a modified version of the Kilosort automatic sorting software (Pachitariu et al., 2016b, Pachitariu et al., 2016a) (available at: https://github.com/MouseLand/Kilosort and https://github.com/dimokaramanlis/KiloSortMEA). The output of Kilosort was visually inspected and manually curated with the “Phy 2” software (https://github.com/cortex-lab/phy). Only those clusters of spikes (“units”) with a well-separated spike waveform and a clear refractory period were included in the final analysis and considered as coming from individual cells. The spike clusters were pre-processed and analyzed using custom-made MATLAB scripts (Version: 9.7.0, R2019b; Mathworks, MA, USA). The raster plots, average firing rate and spike amplitude were measured from the spontaneous activity of each recorded cell. In all our recordings, we observed frequent “bursts” - groups of spikes occurring rapidly and consecutively with short inter-spike intervals [less than few tens of milliseconds (ms)], followed by quiescent periods much longer than typical inter-spike intervals (in our recordings generally several seconds). The bursts typically occurred synchronously for multiple cells over the array and in our analysis, we focused on such population-wide synchronized bursts. To detect the concurrent bursts, we computed the population firing rate as a histogram (100ms bin size) of array-wide spiking activity. The peaks of the firing rate histogram were used to detect synchronous, array-wide bursts with at least 500ms distance between two consecutive peaks. The peaks that were smaller than 1/5 of the largest peak were excluded as they do not correspond to array-wide, synchronous activity. A time window of 650ms around each peak (150ms before to 500ms after) was defined as the burst window (onset and offset of each burst). For each recorded cell, the spikes belonging to bursts were measured during the defined burst windows, and cells with fewer than six spikes across all their detected bursts were excluded from this analysis. From the detected bursts, the following parameters were calculated: (1) burst rate of the culture as the number of bursts per time over the duration of each recording; (2) inter-burst-interval as the time between the measured offset of a burst and the onset of the following burst, calculated for each pair of successive bursts in a recording; (3) burst duration for each cell as the time between the cell’s first and last spike during the burst window; (4) intra-burst-frequency as the rate of spikes occurring within a burst, averaged over all the detected bursts for each cell; and (5) percentage of spikes in bursts as the ratio of spikes occurring during bursts relative to the total number of spikes for each cell.

#### Confocal imaging

Imaging was performed on a Leica SP5 confocal laser scanning microscope equipped with hybrid detectors using Application Suite X software with 100x immersion objective lenses (Leica Biosystems, Wetzlar, Germany). Samples were excited using 405 Diode, argon and helium–neon 633 lasers, pinhole = 1, 0.2 µm thickness Z stacks and 2 averaging line-by-line. The acquisition settings were optimized to avoid underexposure and oversaturation effects and kept equal throughout image acquisition of the samples.

#### STED microscopy

Samples were imaged using an inverse 4-channel Expert Line easy3D STED setup (Abberior Instruments, Göttingen, Germany). The setup was based on an Olympus IX83 microscope body equipped with a plan apochromat 100x 1.4 NA oil-immersion objective (Olympus). Fluorescence lasers 595 nm and 775 nm (Abberior Instruments, Göttingen, Germany) were utilized for the imaging. Fluorescence signal was detected using avalanche photodiodes detectors (Abberior Instruments, Göttingen, Germany) in predefined channels. The operation of the setup and the recording of images were performed with the Imspector software, version 0.14 (Abberior Instruments, Göttingen, Germany).

#### Quantification and statistical analysis

Images were analyzed using ImageJ software (National Institutes of Health) (Schindelin et al., 2012). To analyze the degree of EVs uptake in neuronal cells, measurement of EVs and cell area was performed from isolated areas chosen randomly within regions containing EVs signal. All data are presented as mean ± SD. Data from at least three independent experiments and each replicate represents one independent experiment. To assess differences between two groups, two-tailed unpaired student t-test was performed using GraphPad Prism 9 software (GraphPad, CA, USA). To assess differences between more than two groups, significant differences were assessed by one-way ANOVA followed by multiple comparisons with significance between groups corrected by Bonferroni procedure using GraphPad Prism 9 software (GraphPad, CA, USA). Differences were considered to be significant for values of p<0.05 and are expressed as mean ± SD. For mass spectrometry, the spectral count differences between samples were considered to be significant for FDR values<0.1 (see proteomic analyses section).

## Supplemental Information

**Supplementary Figure 1. Protein hits enriched in ectosomes and exosomes. Volcano plots of quantitative differences in proteins in EVs fractions. (A)** Other proteins enriched in ectosomes - clathrin (*CLTC*), cell division control protein 42 (*CDC42*), integrin alpha-1/ 2/ 5/ 6 (*ITGA1/ 2/ 5/ 6*), 14-3-3 proteins (*YWHAZ/ E/ B/ G/ H/ Q*), histones H1.4 / H1.0/ H2A (*HIST1H1E / H1F0/ HIST2H2AC*), heat shock protein HSP90 (*HSP90AB1/ AA1*), heat shock 70 kDa protein (*HSPA1A/ B, HSPA4*) and heat shock protein 105 kDa (*HSPH1*). **(B)** Ras-related proteins (Rabs) identified in EVs - in exosomes Rab-1A/ 1B/ 4A/ 5B/ 27B (*RAB1A/ 1B/ 4A/ 5B/ 27B*), in ectosomes Rab-2A/ 5C/ 6A/ 6B/ 7A/ 8A/ 8B/ 9A/ 10/ 11B/ 13/ 21/ 23/ 35 (*RAB2A/ 5C/ 6A/ 6B/ 7A/ 8A/ 8B/ 9A/ 10/ 11B/ 13/ 21/ 23/ 35*). Blue dots represent the proteins enrichment in exosomes, while turquoise dots represent enrichment in ectosomes. Proteins in the plots are identified with their gene name. Dots above the dashed line represent proteins for which differences were significant (false discovery rate [FDR] <0.1). Data represented in “t-test Difference (Ectosomes - Exosomes)” vs. “-Log t-test p-value” from five independent samples for each group. Data analyses were performed using Perseus software.

**Supplementary Figure 2. Functional protein association networks were plotted by using the STRING database.** The exosomal proteins significantly identified in our study show robust networks and significant overlap. The line thickness represents the confidence of the association (https://version-11-0b.string-db.org/cgi/network?networkId=br68rv2GDNdl).

**Supplementary Figure 3. Functional protein association networks were plotted by using the STRING database.** The ectosome proteins significantly identified in our study show robust networks and significant overlap. The line thickness represents the confidence of the association (https://version-11-0b.string-db.org/cgi/network?networkId=bwoR4jPEXhpC).

**Supplementary Figure 4. Conditioned media used for the experiments does not contain residual EVs. (A)** Memcode staining from four different replicates from ectosomes and exosomes pellets purified from different media collections showing the efficiency in the isolation protocol. **(B)** Immunoblot of ectosomes and exosomes released by HEK cells, EVs that are present in the conditioned cell media without incubation with the cells and whole-cell lysate. Equal quantities of protein were separated on SDS-PAGE gels, and membranes were blotted for alix, flotillin-1 and Annexin-A2.

**Supplementary Figure 5. Characterization of EGFP and annexin-A2 levels in EVs. (A)** Immunoblot showing the levels of EGFP in ectosomes and exosomes and several EVs markers. **(B)** Immunoblot displaying the levels of mCherry-Annexin-A2 and flotillin-1 in EVs.

**Supplementary Figure 6. Cellular uptake of EVs by primary cortical neurons. (A)** Primary cortical neurons were seeded in microfluidic devices to separate neuronal cell bodies. EVs containing EGFP were added to primary cortical neurons at DIV14 with 20μg/mL of ectosomes or exosomes for 24 hours in the upper left well of the device. **(B)** Top panel shows contrast light images of the primary cortical neurons growing in microfluidic devices. Bottom panel shows neurons cell bodies and axons labelled with membrane cell dye Alexa Fluor 633 C5-maleimide (scale bar 100µm). **(C)** Lactate DeHydrogenase (LDH) measurements show that EVs enriched with EGFP or labelled with dye show similar low toxicity for the primary cortical neurons.

**Supplementary Figure 7. EVs modulate spontaneous activity in primary cortical neurons. (A)** MEA recordings were used to monitor neural spike firing after treating primary cortical neurons with 20μg/mL of ectosomes or exosomes at DIV14 for 24 hours. Schematic representation shows the steps performed for the data analysis. **(B)** Phase-contrast image of primary cortical neurons cultured on an MEA chamber. **(C)** Bursts rate (per minute) of the cortical neurons treated with EVs. **(D)** Quantification of the mean firing rate and average spike amplitude of primary cortical neurons incubated with PBS, ectosomes or exosomes. Data from at least three independent experiments for each condition. Significant differences were assessed by one-way ANOVA followed by multiple comparisons with significance between groups corrected by Bonferroni procedure. Differences were considered to be significant for values of p < 0.05 and are expressed as mean ± SD, *p<0.05, **p<0.01.

## Supplementary Table legends

**Supplementary Table 1.** List of significantly altered proteins identified in exosomes. Common proteins (11) in the list of Top 100 proteins often identified in EVs are highlighted in bold (see Supplementary Table 3).

**Supplementary Table 2.** List of significantly altered proteins identified in ectosomes. Common proteins (52) in the list of Top 100 proteins often identified in EVs are highlighted in bold (see Supplementary Table 3).

**Supplementary Table 3.** Table of the Top 100 proteins often identified in EVs (source: vesiclepedia, http://microvesicles.org/extracellular_vesicle_markers).

## Resources Table

| REAGENT or RESOURCE | SOURCE | IDENTIFIER |
| --- | --- | --- |
| **Antibodies** |  |  |
| Mouse monoclonal anti- alix | Abcam | Cat#ab117600 |
| Mouse monoclonal anti-alpha-tubulin | Sigma-Aldrich | Cat#T9026 |
| Mouse monoclonal anti- β-Actin | Sigma-Aldrich | Cat#A5441 |
| Rabbit monoclonal anti- annexin-A2 | Abcam | Cat#ab178677 |
| Rabbit polyclonal anti- annexin-A5 | Cell signalling | Cat#8555 |
| Rabbit polyclonal anti- calnexin | Abcam | Cat#ab22595 |
| Rabbit polyclonal anti- CD54 | Cell Signalling | Cat#4915 |
| Rabbit monoclonal anti- CD9 | Cell Signalling | Cat#13174 |
| Rabbit monoclonal anti- flotillin-1 | Cell Signalling | Cat#18634 |
| Mouse monoclonal anti- GFP | Roche | Cat#11814460001 |
| Rabbit monoclonal anti-GM130 | Cell Signalling | Cat#12480 |
| Rabbit polyclonal anti- LC3 | MBL International | Cat#PM036 |
| Mouse monoclonal anti- MAP2 | Abcam | Cat#ab11267 |
| Rabbit polyclonal anti- mCherry | Abcam | Cat#ab167453 |
| Rabbit polyclonal anti- MAP2 | Proteintech | Cat#17490-1-AP |
| Mouse monoclonal anti- MMP-2 | Santa Cruz | Cat#sc-13595 |
| Rabbit polyclonal anti- TSG101 | Abcam | Cat#ab30871 |
| Alexa Fluor 555 donkey anti-rabbit | Invitrogen | Cat#A31572 |
| Alexa Fluor 555 donkey anti-mouse | Invitrogen | Cat#A31570 |
| ECL™ Mouse IgG, HRP-linked whole antibody | Amersham | Cat#NXA931 |
| ECL™ Rabbit IgG, HRP-linked whole antibody | Amersham | Cat#NA934V |
| **Biological samples** |  |  |
| Human CSF | University Medical Center<br>Göttingen (Göttingen,<br>Germany) | N/A |
| **Critical commercial assays** |  |  |
| MemCode™ Reversible Protein | ThermoScientific | 24580 |
| **Experimental models: Cell lines** |  |  |
| Human embryonic kidney (HEK) 293 cells | ATCC | CRL-11268 |
| **Experimental models: Organisms/strains** |  |  |
| Primary cortical neurons (C57BL6/J#00245 wild-type E15.5 embryos) | University Medical Center<br>Göttingen (Göttingen,<br>Germany) | N/A |
| **Recombinant DNA** |  |  |
| Annexin-A2-GFP | (Rescher et al., 2000) | Plasmid #107196,<br>Addgene |
| FUGW-mCherry-Annexin-A2 | Home-made | N/A |
| pRRL-CMV-EGFP-PRE-SIN | Home-made | N/A |
| **Software and algorithms** |  |  |
| Perseus (version and 1.6.15.0) | (Tyanova et al., 2016) | https://maxquant.net/perseus/ |
| MultiChannel Experimenter software (version 2.17.7.0) | N/A | https://www.multichannelsystems.com/downloads/software |
| Kilosort | (Pachitariu et al., 2016) | https://github.com/MouseLand/Kilosort |
| Phy2 | N/A | https://github.com/cortex-lab/phy |
| MATLAB R2019b (Version: 9.7.0) | Mathworks | https://de.mathworks.com/products/matlab.html |
| Prism 9 | GraphPad | https://www.graphpad.com/scientific-software/prism/ |
| Fiji | (Schindelin et al., 2012) | https://imagej.net/Downloads |
| Other |  |  |
| Wheat germ agglutinin (WGA), Alexa Fluor™ 633 conjugate | Invitrogen | W21404 |
| Alexa Fluor™ 633 C5-maleimide | Molecular probes | A20342 |
| DAPI | Roth | 6335.1 |

- PACHITARIU, M., STEINMETZ, N., KADIR, S., CARANDINI, M. & HARRIS, K. Fast and accurate spike sorting of high-channel count probes with KiloSort. NIPS Proceedings, 2016. Neural Information Systems Foundation, Inc. - RESCHER, U., ZOBIACK, N. & GERKE, V. 2000. Intact Ca(2+)-binding sites are required for targeting of annexin 1 to endosomal membranes in living HeLa cells. *J Cell Sci*, 113 ( Pt 22), 3931-8. - SCHINDELIN, J., ARGANDA-CARRERAS, I., FRISE, E., KAYNIG, V., LONGAIR, M., PIETZSCH, T., PREIBISCH, S., RUEDEN, C., SAALFELD, S., SCHMID, B., TINEVEZ, J. Y., WHITE, D. J., HARTENSTEIN, V., ELICEIRI, K., TOMANCAK, P. & CARDONA, A. 2012. Fiji: an open-source platform for biological-image analysis. *Nat Methods*, 9, 676-82. - TYANOVA, S., TEMU, T., SINITYN, P., CARLSON, A., HEIN, M. Y., GEIGER, T., MANN, M. & COX, J. 2016. The Perseus computational platform for comprehensive analysis of (prote)omics data. *Nat Methods*, 13, 731-40.

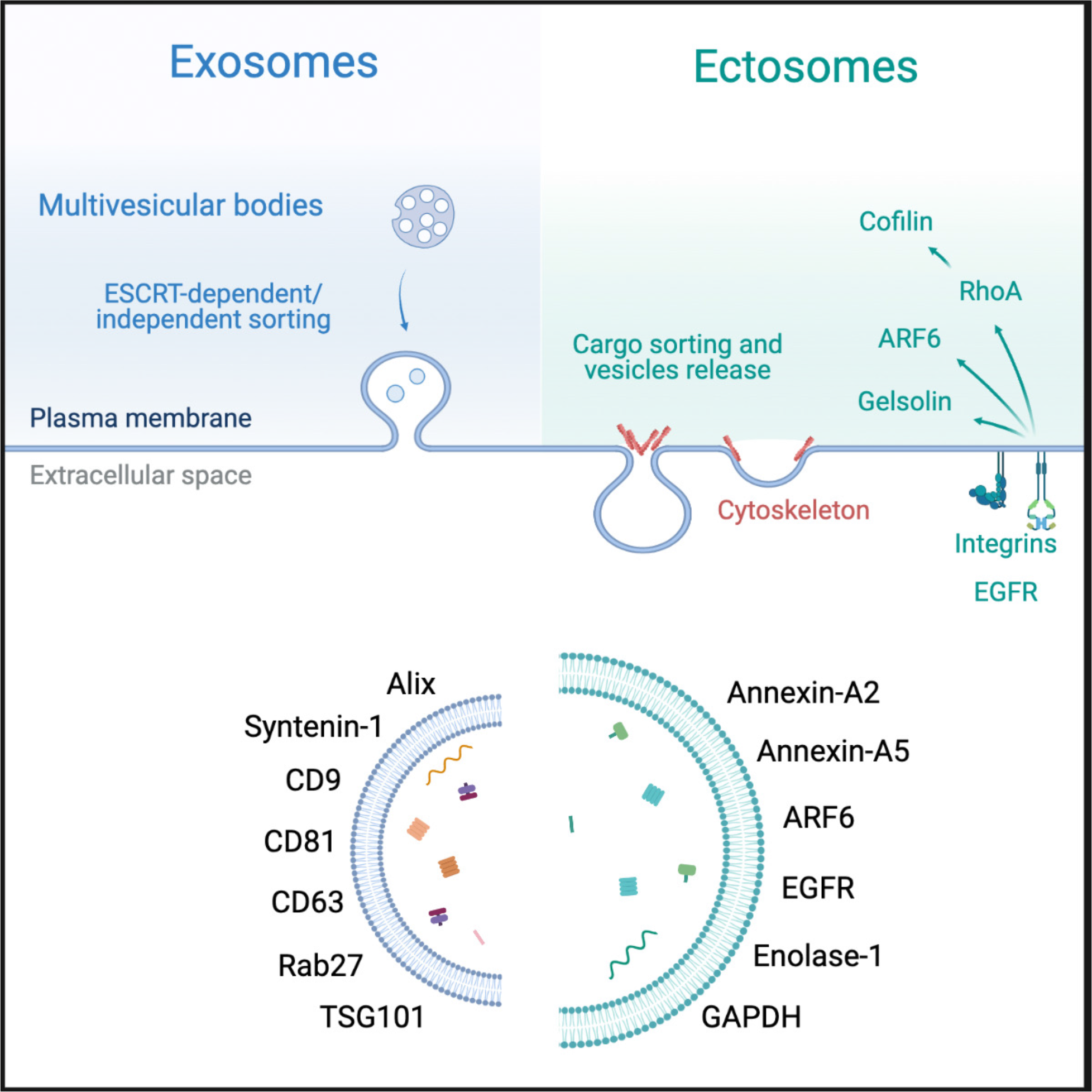

## References

1. Andreu, Z. & Yanez-Mo, M. 2014. Tetraspanins in extracellular vesicle formation and function. Front Immunol, 5, 442.

2. Antonucci, F., Turola, E., Riganti, L., Caleo, M., Gabrielli, M., Perrotta, C., Novellino, L., Clementi, E., Giussani, P., Viani, P., Matteoli, M. & Verderio, C. 2012. Microvesicles released from microglia stimulate synaptic activity via enhanced sphingolipid metabolism. Embo J, 31, 1231–40.

3. Basso, M. & Bonetto, V. 2016. Extracellular Vesicles and a Novel Form of Communication in the Brain. Front Neurosci, 10, 127.

4. Bellingham, S. A., Guo, B. B., Coleman, B. M. & Hill, A. F. 2012. Exosomes: vehicles for the transfer of toxic proteins associated with neurodegenerative diseases? Front Physiol, 3, 124.

5. Chiasserini, D., Van Weering, J. R., Piersma, S. R., Pham, T. V., Malekzadeh, A., Teunissen, C. E., De Wit, H. & Jimenez, C. R. 2014. Proteomic analysis of cerebrospinal fluid extracellular vesicles: a comprehensive dataset. J Proteomics, 106, 191–204.

6. Chiou, N.-T. A., K. Mark 2016. Improved exosome isolation by sucrose gradient fractionation of ultracentrifuged crude exosome pellets. protocolexchange.

7. Choi, D. S., Kim, D. K., Kim, Y. K. & Gho, Y. S. 2013. Proteomics, transcriptomics and lipidomics of exosomes and ectosomes. Proteomics, 13, 1554–71.

8. Choi, D. S., Kim, D. K., Kim, Y. K. & Gho, Y. S. 2015. Proteomics of extracellular vesicles: Exosomes and ectosomes. Mass Spectrom Rev, 34, 474–90.

9. Clayton, A., Court, J., Navabi, H., Adams, M., Mason, M. D., Hobot, J. A., Newman, G. R. & Jasani, B. 2001. Analysis of antigen presenting cell derived exosomes, based on immuno-magnetic isolation and flow cytometry. J Immunol Methods, 247, 163–74.

10. Cocucci, E. & Meldolesi, J. 2015. Ectosomes and exosomes: shedding the confusion between extracellular vesicles. Trends Cell Biol, 25, 364–72.

11. Coleman, B. M. & Hill, A. F. 2015. Extracellular vesicles--Their role in the packaging and spread of misfolded proteins associated with neurodegenerative diseases. Semin Cell Dev Biol, 40, 89–96.

12. Colombo, M., Raposo, G. & Thery, C. 2014. Biogenesis, secretion, and intercellular interactions of exosomes and other extracellular vesicles. Annu Rev Cell Dev Biol, 30, 255–89.

13. Cox, J. & Mann, M. 2008. MaxQuant enables high peptide identification rates, individualized p.p.b.-range mass accuracies and proteome-wide protein quantification. Nat Biotechnol, 26, 1367–72.

14. Dujardin, S., Begard, S., Caillierez, R., Lachaud, C., Delattre, L., Carrier, S., Loyens, A., Galas, M. C., Bousset, L., Melki, R., Auregan, G., Hantraye, P., Brouillet, E., Buee, L. & Colin, M. 2014. Ectosomes: a new mechanism for non-exosomal secretion of tau protein. PloS One, 9, e100760.

15. Escrevente, C., Keller, S., Altevogt, P. & Costa, J. 2011. Interaction and uptake of exosomes by ovarian cancer cells. Bmc Cancer, 11, 108.

16. Freundt, E. C., Maynard, N., Clancy, E. K., Roy, S., Bousset, L., Sourigues, Y., Covert, M., Melki, R., Kirkegaard, K. & Brahic, M. 2012. Neuron-to-neuron transmission of alpha-synuclein fibrils through axonal transport. Ann Neurol, 72, 517–24.

17. Frohlich, D., Kuo, W. P., Fruhbeis, C., Sun, J. J., Zehendner, C. M., Luhmann, H. J., Pinto, S., Toedling, J., Trotter, J. & Kramer-Albers, E. M. 2014. Multifaceted effects of oligodendroglial exosomes on neurons: impact on neuronal firing rate, signal transduction and gene regulation. Philos Trans R Soc Lond B Biol Sci, 369.

18. Gangoda, L., Boukouris, S., Liem, M., Kalra, H. & Mathivanan, S. 2015. Extracellular vesicles including exosomes are mediators of signal transduction: are they protective or pathogenic? Proteomics, 15, 260–71.

19. Harris, J., Lee, H., Vahidi, B., Tu, C., Cribbs, D., Cotman, C. & Jeon, N. L. 2007. Non-plasma bonding of Pdms for inexpensive fabrication of microfluidic devices. J Vis Exp, 410.

20. Hessvik, N. P. & Llorente, A. 2018. Current knowledge on exosome biogenesis and release. Cell Mol Life Sci, 75, 193–208.

21. Hoshino, A., Costa-Silva, B., Shen, T. L., Rodrigues, G., Hashimoto, A., Tesic Mark, M., Molina, H., Kohsaka, S., Di Giannatale, A., Ceder, S., Singh, S., Williams, C., Soplop, N., Uryu, K., Pharmer, L., King, T., Bojmar, L., Davies, A. E., Ararso, Y., Zhang, T., Zhang, H., Hernandez, J., Weiss, J. M., Dumont-Cole, V. D., Kramer, K., Wexler, L. H., Narendran, A., Schwartz, G. K., Healey, J. H., Sandstrom, P., Labori, K. J., Kure, E. H., Grandgenett, P. M., Hollingsworth, M. A., De Sousa, M., Kaur, S., Jain, M., Mallya, K., Batra, S. K., Jarnagin, W. R., Brady, M. S., Fodstad, O., Muller, V., Pantel, K., Minn, A. J., Bissell, M. J., Garcia, B. A., Kang, Y., Rajasekhar, V. K., Ghajar, C. M., Matei, I., Peinado, H., Bromberg, J. & Lyden, D. 2015. Tumour exosome integrins determine organotropic metastasis. Nature, 527, 329–35.

22. Jeppesen, D. K., Fenix, A. M., Franklin, J. L., Higginbotham, J. N., Zhang, Q., Zimmerman, L. J., Liebler, D. C., Ping, J., Liu, Q., Evans, R., Fissell, W. H., Patton, J. G., Rome, L. H., Burnette, D. T. & Coffey, R. J. 2019. Reassessment of Exosome Composition. Cell, 177, 428–445 e18.

23. Kalra, H., Drummen, G. P. & Mathivanan, S. 2016. Focus on Extracellular Vesicles: Introducing the Next Small Big Thing. Int J Mol Sci, 17, 170.

24. Kalra, H., Simpson, R. J., Ji, H., Aikawa, E., Altevogt, P., Askenase, P., Bond, V. C., Borras, F. E., Breakefield, X., Budnik, V., Buzas, E., Camussi, G., Clayton, A., Cocucci, E., Falcon-Perez, J. M., Gabrielsson, S., Gho, Y. S., Gupta, D., Harsha, H. C., Hendrix, A., Hill, A. F., Inal, J. M., Jenster, G., Kramer-Albers, E. M., Lim, S. K., Llorente, A., Lotvall, J., Marcilla, A., Mincheva-Nilsson, L., Nazarenko, I., Nieuwland, R., Nolte-’T Hoen, E. N., Pandey, A., Patel, T., Piper, M. G., Pluchino, S., Prasad, T. S., Rajendran, L., Raposo, G., Record, M., Reid, G. E., Sanchez-Madrid, F., Schiffelers, R. M., Siljander, P., Stensballe, A., Stoorvogel, W., Taylor, D., Thery, C., Valadi, H., Van Balkom, B. W., Vazquez, J., Vidal, M., Wauben, M. H., Yanez-Mo, M., Zoeller, M. & Mathivanan, S. 2012. Vesiclepedia: a compendium for extracellular vesicles with continuous community annotation. PloS Biol, 10, e1001450.

25. Keerthikumar, S., Chisanga, D., Ariyaratne, D., Al Saffar, H., Anand, S., Zhao, K., Samuel, M., Pathan, M., Jois, M., Chilamkurti, N., Gangoda, L. & Mathivanan, S. 2016. ExoCarta: A Web-Based Compendium of Exosomal Cargo. J Mol Biol, 428, 688–692.

26. Keerthikumar, S., Gangoda, L., Liem, M., Fonseka, P., Atukorala, I., Ozcitti, C., Mechler, A., Adda, C. G., Ang, C. S. & Mathivanan, S. 2015. Proteogenomic analysis reveals exosomes are more oncogenic than ectosomes. Oncotarget, 6, 15375–96.

27. Khani, M. H. & Gollisch, T. 2021. Linear and nonlinear chromatic integration in the mouse retina. Nat Commun, 12, 1900.

28. Kowal, J., Arras, G., Colombo, M., Jouve, M., Morath, J. P., Primdal-Bengtson, B., Dingli, F., Loew, D., Tkach, M. & Thery, C. 2016. Proteomic comparison defines novel markers to characterize heterogeneous populations of extracellular vesicle subtypes. Proc Natl Acad Sci U S A, 113, E968–77.

29. Lotvall, J., Hill, A. F., Hochberg, F., Buzas, E. I., Di Vizio, D., Gardiner, C., Gho, Y. S., Kurochkin, I. V., Mathivanan, S., Quesenberry, P., Sahoo, S., Tahara, H., Wauben, M. H., Witwer, K. W. & Thery, C. 2014. Minimal experimental requirements for definition of extracellular vesicles and their functions: a position statement from the International Society for Extracellular Vesicles. J Extracell Vesicles, 3, 26913.

30. Mathieu, M., Martin-Jaular, L., Lavieu, G. & Thery, C. 2019. Specificities of secretion and uptake of exosomes and other extracellular vesicles for cell-to-cell communication. Nat Cell Biol, 21, 9–17.

31. Mathivanan, S., Ji, H. & Simpson, R. J. 2010. Exosomes: extracellular organelles important in intercellular communication. J Proteomics, 73, 1907–20.

32. Meldolesi, J. 2018. Exosomes and Ectosomes in Intercellular Communication. Curr Biol, 28, R435–R444.

33. Mulcahy, L. A., Pink, R. C. & Carter, D. R. 2014. Routes and mechanisms of extracellular vesicle uptake. J Extracell Vesicles, 3.

34. Muralidharan-Chari, V., Clancy, J., Plou, C., Romao, M., Chavrier, P., Raposo, G. & D’souza-Schorey, C. 2009. Arf6-regulated shedding of tumor cell-derived plasma membrane microvesicles. Curr Biol, 19, 1875–85.

35. Nakase, I., Kobayashi, N. B., Takatani-Nakase, T. & Yoshida, T. 2015. Active macropinocytosis induction by stimulation of epidermal growth factor receptor and oncogenic Ras expression potentiates cellular uptake efficacy of exosomes. Sci Rep, 5, 10300.

36. Pachitariu, M., Steinmetz, N., Kadir, S. & Carandini, M. 2016a. Kilosort: realtime spike-sorting for extracellular electrophysiology with hundreds of channels. BioRxiv, 061481.

37. Pachitariu, M., Steinmetz, N., Kadir, S., Carandini, M. & Harris, K. Fast and accurate spike sorting of high-channel count probes with KiloSort. Nips Proceedings, 2016b. Neural Information Systems Foundation, Inc.

38. Paolicelli, R. C., Bergamini, G. & Rajendran, L. 2019. Cell-to-cell Communication by Extracellular Vesicles: Focus on Microglia. Neuroscience, 405, 148–157.

39. Pathan, M., Fonseka, P., Chitti, S. V., Kang, T., Sanwlani, R., Van Deun, J., Hendrix, A. & Mathivanan, S. 2019. Vesiclepedia 2019: a compendium of Rna, proteins, lipids and metabolites in extracellular vesicles. Nucleic Acids Res, 47, D516–D519.

40. Popa, S. J., Stewart, S. E. & Moreau, K. 2018. Unconventional secretion of annexins and galectins. Semin Cell Dev Biol, 83, 42–50.

41. Raposo, G. & Stoorvogel, W. 2013. Extracellular vesicles: exosomes, microvesicles, and friends. J Cell Biol, 200, 373–83.

42. Roberts-Dalton, H. D., Cocks, A., Falcon-Perez, J. M., Sayers, E. J., Webber, J. P., Watson, P., Clayton, A. & Jones, A. T. 2017. Fluorescence labelling of extracellular vesicles using a novel thiol-based strategy for quantitative analysis of cellular delivery and intracellular traffic. Nanoscale, 9, 13693–13706.

43. Sadallah, S., Eken, C. & Schifferli, J. A. 2011. Ectosomes as modulators of inflammation and immunity. Clin Exp Immunol, 163, 26–32.

44. Schindelin, J., Arganda-Carreras, I., Frise, E., Kaynig, V., Longair, M., Pietzsch, T., Preibisch, S., Rueden, C., Saalfeld, S., Schmid, B., Tinevez, J. Y., White, D. J., Hartenstein, V., Eliceiri, K., Tomancak, P. & Cardona, A. 2012. Fiji: an open-source platform for biological-image analysis. Nat Methods, 9, 676–82.

45. Sharma, P., Mesci, P., Carromeu, C., Mcclatchy, D. R., Schiapparelli, L., Yates, J. R., 3rd, Muotri, A. R. & Cline, H. T. 2019. Exosomes regulate neurogenesis and circuit assembly. Proc Natl Acad Sci U S A, 116, 16086–16094.

46. Shelke, G. V., Lasser, C., Gho, Y. S. & Lotvall, J. 2014. Importance of exosome depletion protocols to eliminate functional and Rna-containing extracellular vesicles from fetal bovine serum. J Extracell Vesicles, 3.

47. Shevchenko, A., Tomas, H., Havlis, J., Olsen, J. V. & Mann, M. 2006. In-gel digestion for mass spectrometric characterization of proteins and proteomes. Nat Protoc, 1, 2856–60.

48. Stein, J. M. & Luzio, J. P. 1991. Ectocytosis caused by sublytic autologous complement attack on human neutrophils. The sorting of endogenous plasma-membrane proteins and lipids into shed vesicles. Biochem J, 274 (Pt 2), 381–6.

49. Suetsugu, A., Honma, K., Saji, S., Moriwaki, H., Ochiya, T. & Hoffman, R. M. 2013. Imaging exosome transfer from breast cancer cells to stroma at metastatic sites in orthotopic nude-mouse models. Adv Drug Deliv Rev, 65, 383–90.

50. Sun, J. J., Kilb, W. & Luhmann, H. J. 2010. Self-organization of repetitive spike patterns in developing neuronal networks in vitro. Eur J Neurosci, 32, 1289–99.

51. Szego, E. M., Dominguez-Meijide, A., Gerhardt, E., Konig, A., Koss, D. J., Li, W., Pinho, R., Fahlbusch, C., Johnson, M., Santos, P., Villar-Pique, A., Thom, T., Rizzoli, S., Schmitz, M., Li, J., Zerr, I., Attems, J., Jahn, O. & Outeiro, T. F. 2019. Cytosolic Trapping of a Mitochondrial Heat Shock Protein Is an Early Pathological Event in Synucleinopathies. Cell Rep, 28, 65–77 e6.

52. Szklarczyk, D., Gable, A. L., Lyon, D., Junge, A., Wyder, S., Huerta-Cepas, J., Simonovic, M., Doncheva, N. T., Morris, J. H., Bork, P., Jensen, L. J. & Mering, C. V. 2019. String v11: protein-protein association networks with increased coverage, supporting functional discovery in genome-wide experimental datasets. Nucleic Acids Res, 47, D607–D613.

53. Takeda, S., Wegmann, S., Cho, H., Devos, S. L., Commins, C., Roe, A. D., Nicholls, S. B., Carlson, G. A., Pitstick, R., Nobuhara, C. K., Costantino, I., Frosch, M. P., Muller, D. J., Irimia, D. & Hyman, B. T. 2015. Neuronal uptake and propagation of a rare phosphorylated high-molecular-weight tau derived from Alzheimer’s disease brain. Nat Commun, 6, 8490.

54. Taraboletti, G., D’ascenzo, S., Borsotti, P., Giavazzi, R., Pavan, A. & Dolo, V. 2002. Shedding of the matrix metalloproteinases Mmp-2, Mmp-9, and Mt1-Mmp as membrane vesicle-associated components by endothelial cells. Am J Pathol, 160, 673–80.

55. Tario, J. D., Jr., Humphrey, K., Bantly, A. D., Muirhead, K. A., Moore, J. S. & Wallace, P. K. 2012. Optimized staining and proliferation modeling methods for cell division monitoring using cell tracking dyes. *J Vis Exp*, e4287.

56. Temchura, V. V., Tenbusch, M., Nchinda, G., Nabi, G., Tippler, B., Zelenyuk, M., Wildner, O., Uberla, K. & Kuate, S. 2008. Enhancement of immunostimulatory properties of exosomal vaccines by incorporation of fusion-competent G protein of vesicular stomatitis virus. Vaccine, 26, 3662–72.

57. Thery, C., Amigorena, S., Raposo, G. & Clayton, A. 2006. Isolation and characterization of exosomes from cell culture supernatants and biological fluids. Curr Protoc Cell Biol, Chapter 3, Unit 3 22.

58. Thery, C., Witwer, K. W., Aikawa, E., Alcaraz, M. J., Anderson, J. D., Andriantsitohaina, R., Antoniou, A., Arab, T., Archer, F., Atkin-Smith, G. K., Ayre, D. C., Bach, J. M., Bachurski, D., Baharvand, H., Balaj, L., Baldacchino, S., Bauer, N. N., Baxter, A. A., Bebawy, M., Beckham, C., Bedina Zavec, A., Benmoussa, A., Berardi, A. C., Bergese, P., Bielska, E., Blenkiron, C., Bobis-Wozowicz, S., Boilard, E., Boireau, W., Bongiovanni, A., Borras, F. E., Bosch, S., Boulanger, C. M., Breakefield, X., Breglio, A. M., Brennan, M. A., Brigstock, D. R., Brisson, A., Broekman, M. L., Bromberg, J. F., Bryl-Gorecka, P., Buch, S., Buck, A. H., Burger, D., Busatto, S., Buschmann, D., Bussolati, B., Buzas, E. I., Byrd, J. B., Camussi, G., Carter, D. R., Caruso, S., Chamley, L. W., Chang, Y. T., Chen, C., Chen, S., Cheng, L., Chin, A. R., Clayton, A., Clerici, S. P., Cocks, A., Cocucci, E., Coffey, R. J., Cordeiro-Da-Silva, A., Couch, Y., Coumans, F. A., Coyle, B., Crescitelli, R., Criado, M. F., D’souza-Schorey, C., Das, S., Datta Chaudhuri, A., De Candia, P., De Santana, E. F., De Wever, O., Del Portillo, H. A., Demaret, T., Deville, S., Devitt, A., Dhondt, B., Di Vizio, D., Dieterich, L. C., Dolo, V., Dominguez Rubio, A. P., Dominici, M., Dourado, M. R., Driedonks, T. A., Duarte, F. V., Duncan, H. M., Eichenberger, R. M., Ekstrom, K., El Andaloussi, S., Elie-Caille, C., Erdbrugger, U., Falcon-Perez, J. M., Fatima, F., Fish, J. E., Flores-Bellver, M., Forsonits, A., Frelet-Barrand, A., et al. 2018. Minimal information for studies of extracellular vesicles 2018 (Misev2018): a position statement of the International Society for Extracellular Vesicles and update of the Misev2014 guidelines. J Extracell Vesicles, 7, 1535750.

59. Thompson, A. G., Gray, E., Heman-Ackah, S. M., Mager, I., Talbot, K., Andaloussi, S. E., Wood, M. J. & Turner, M. R. 2016. Extracellular vesicles in neurodegenerative disease - pathogenesis to biomarkers. Nat Rev Neurol, 12, 346–57.

60. Tiscornia, G., Singer, O. & Verma, I. M. 2006. Production and purification of lentiviral vectors. Nat Protoc, 1, 241–5.

61. Tyanova, S., Temu, T., Sinitcyn, P., Carlson, A., Hein, M. Y., Geiger, T., Mann, M. & Cox, J. 2016. The Perseus computational platform for comprehensive analysis of (prote)omics data. Nat Methods, 13, 731–40.

62. Van Niel, G., D’angelo, G. & Raposo, G. 2018. Shedding light on the cell biology of extracellular vesicles. Nat Rev Mol Cell Biol, 19, 213–228.

63. Vietri, M., Radulovic, M. & Stenmark, H. 2020. The many functions of Escrts. Nat Rev Mol Cell Biol, 21, 25–42.

64. Von Mering, C., Jensen, L. J., Snel, B., Hooper, S. D., Krupp, M., Foglierini, M., Jouffre, N., Huynen, M. A. & Bork, P. 2005. String: known and predicted protein-protein associations, integrated and transferred across organisms. Nucleic Acids Res, 33, D433–7.

65. Wagenaar, D. A., Pine, J. & Potter, S. M. 2004. Effective parameters for stimulation of dissociated cultures using multi-electrode arrays. J Neurosci Methods, 138, 27–37.

66. Willms, E., Johansson, H. J., Mager, I., Lee, Y., Blomberg, K. E., Sadik, M., Alaarg, A., Smith, C. I., Lehtio, J., El Andaloussi, S., Wood, M. J. & Vader, P. 2016. Cells release subpopulations of exosomes with distinct molecular and biological properties. Sci Rep, 6, 22519.

67. You, Y., Borgmann, K., Edara, V. V., Stacy, S., Ghorpade, A. & Ikezu, T. 2020. Activated human astrocyte-derived extracellular vesicles modulate neuronal uptake, differentiation and firing. J Extracell Vesicles, 9, 1706801.

68. You, Y. & Ikezu, T. 2019. Emerging roles of extracellular vesicles in neurodegenerative disorders. Neurobiol Dis, 130, 104512.

69. Zhang, H., Freitas, D., Kim, H. S., Fabijanic, K., Li, Z., Chen, H., Mark, M. T., Molina, H., Martin, A. B., Bojmar, L., Fang, J., Rampersaud, S., Hoshino, A., Matei, I., Kenific, C. M., Nakajima, M., Mutvei, A. P., Sansone, P., Buehring, W., Wang, H., Jimenez, J. P., Cohen-Gould, L., Paknejad, N., Brendel, M., Manova-Todorova, K., Magalhaes, A., Ferreira, J. A., Osorio, H., Silva, A. M., Massey, A., Cubillos-Ruiz, J. R., Galletti, G., Giannakakou, P., Cuervo, A. M., Blenis, J., Schwartz, R., Brady, M. S., Peinado, H., Bromberg, J., Matsui, H., Reis, C. A. & Lyden, D. 2018. Identification of distinct nanoparticles and subsets of extracellular vesicles by asymmetric flow field-flow fractionation. Nat Cell Biol, 20, 332–343.

