## Supplementary figures for "Ectosomes and exosomes are distinct proteomic entities that modulate spontaneous activity in neuronal cells"

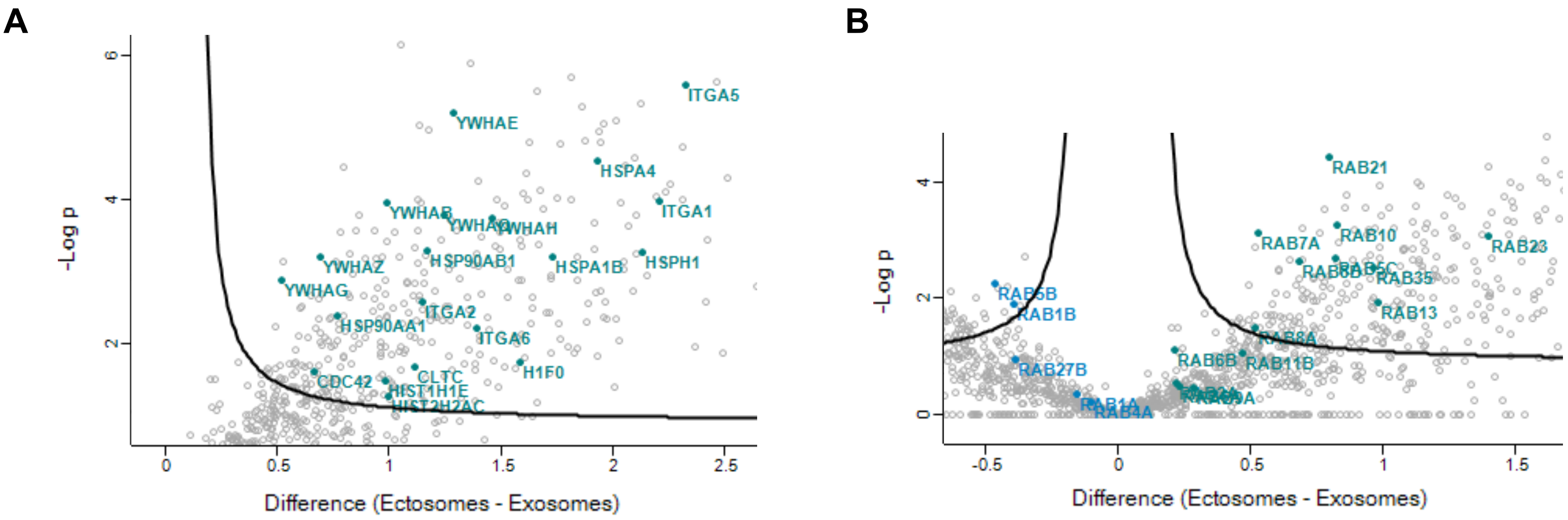


**Supplementary Figure 1. Protein hits enriched in ectosomes and exosomes. Volcano plots of quantitative differences in proteins in EVs fractions.**

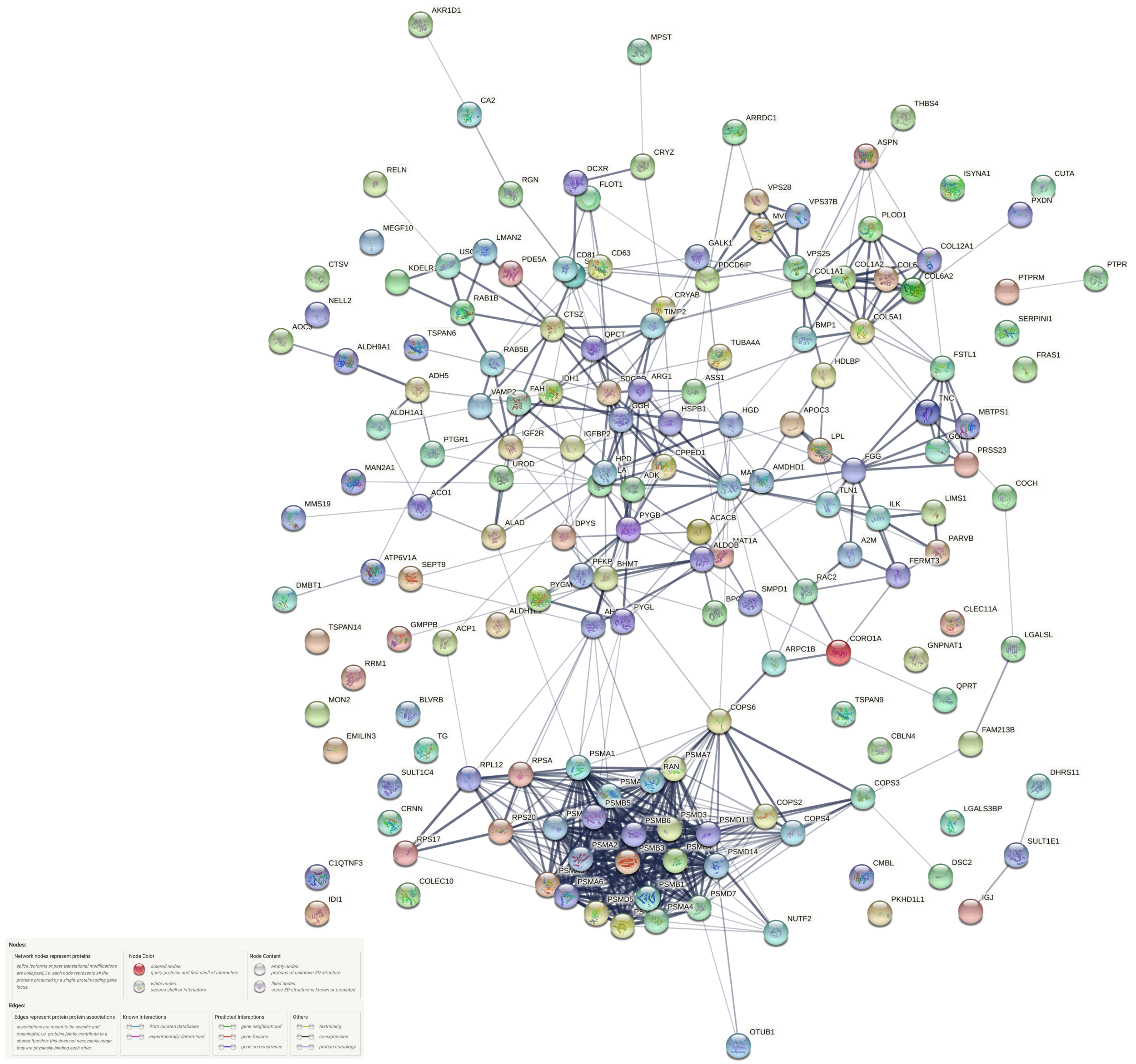


**Supplementary Figure 2. Functional protein association networks were plotted by using the STRING database.**

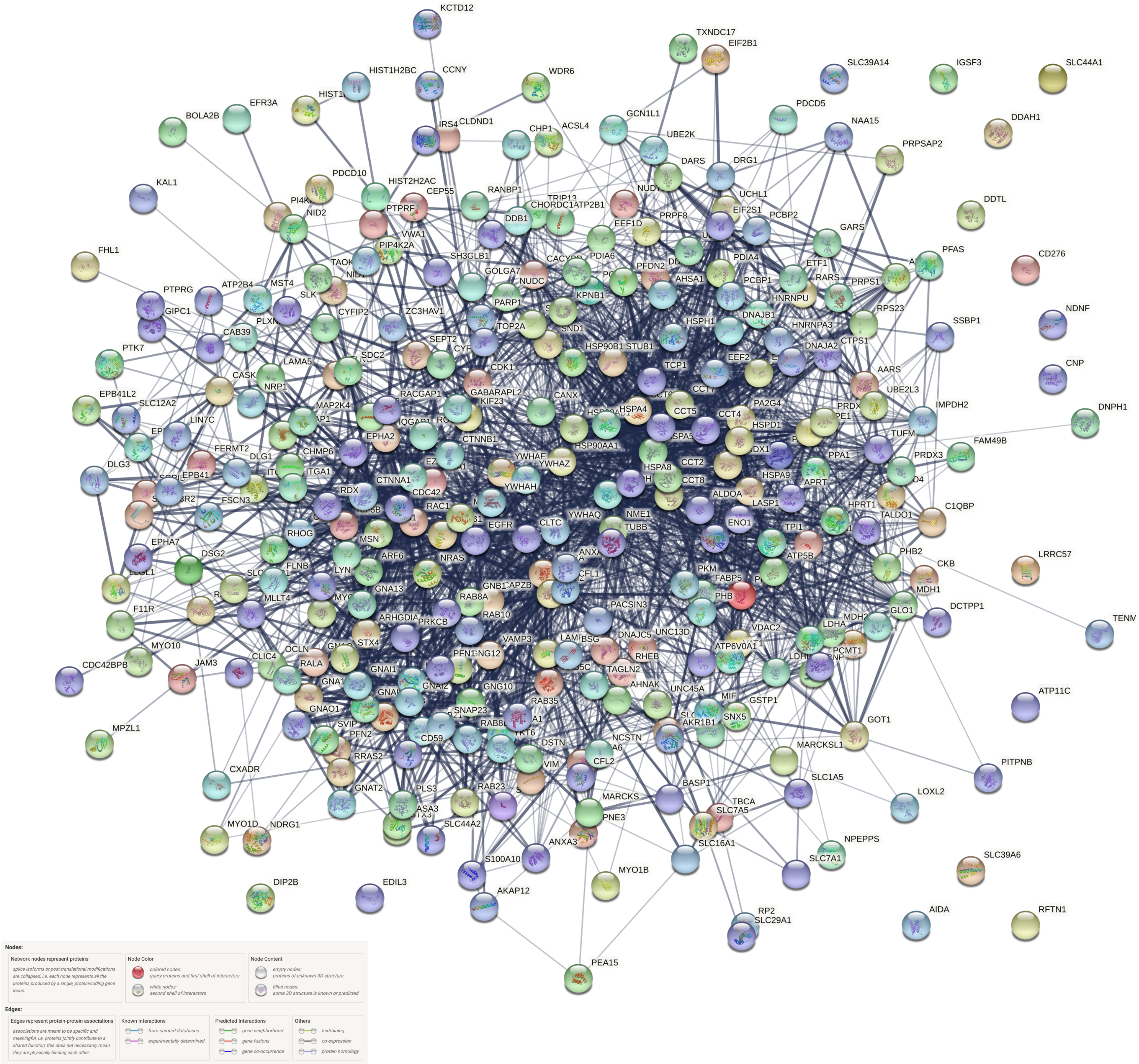


**Supplementary Figure 3. Functional protein association networks were plotted by using the STRING database.**

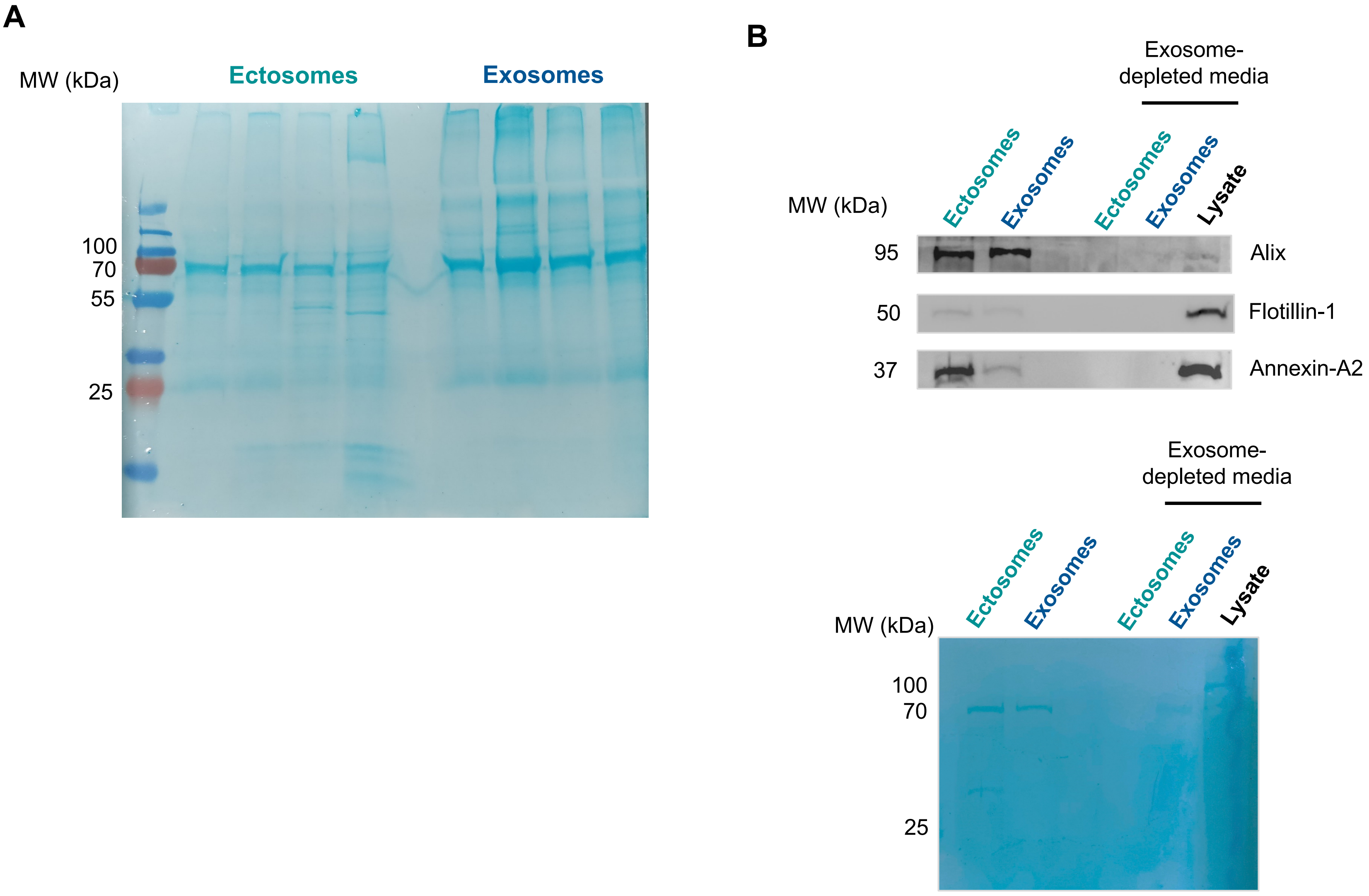


**Supplementary Figure 4. Conditioned media used for the experiments does not contain residual EVs.**

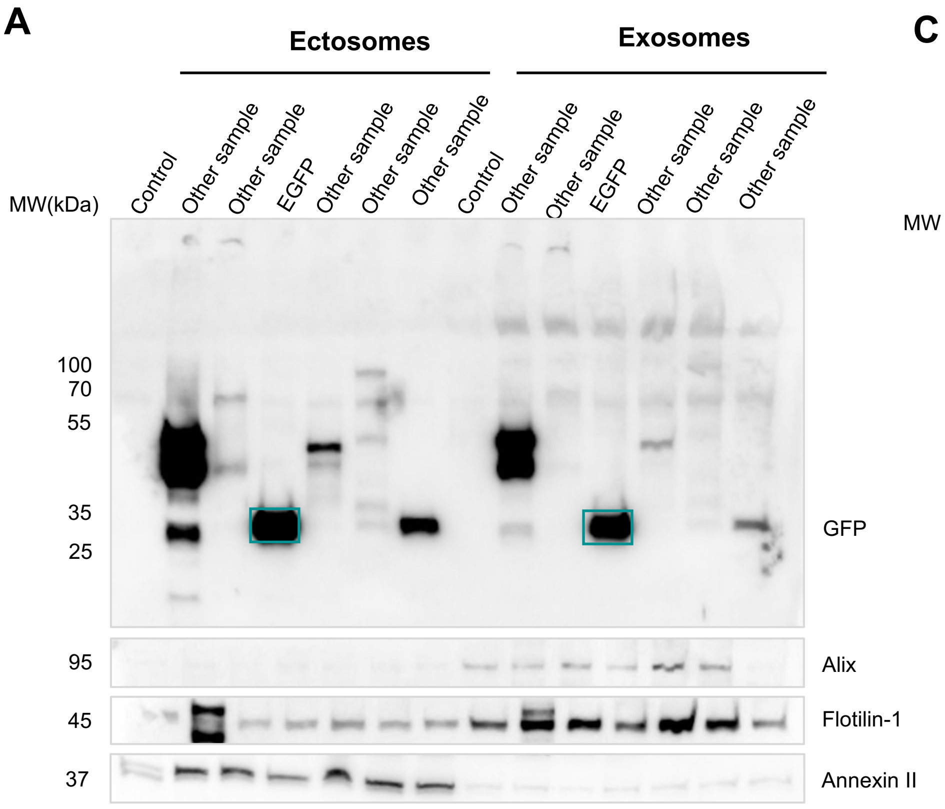


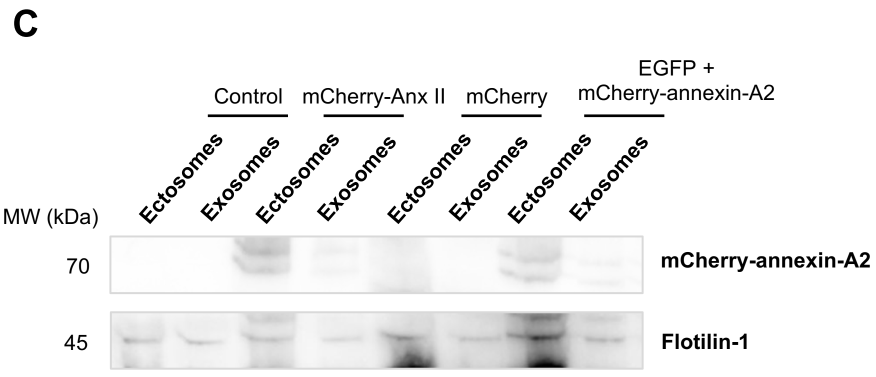


**Supplementary Figure 5. Characterisation of EGFP and annexin-A2 levels in EVs.**

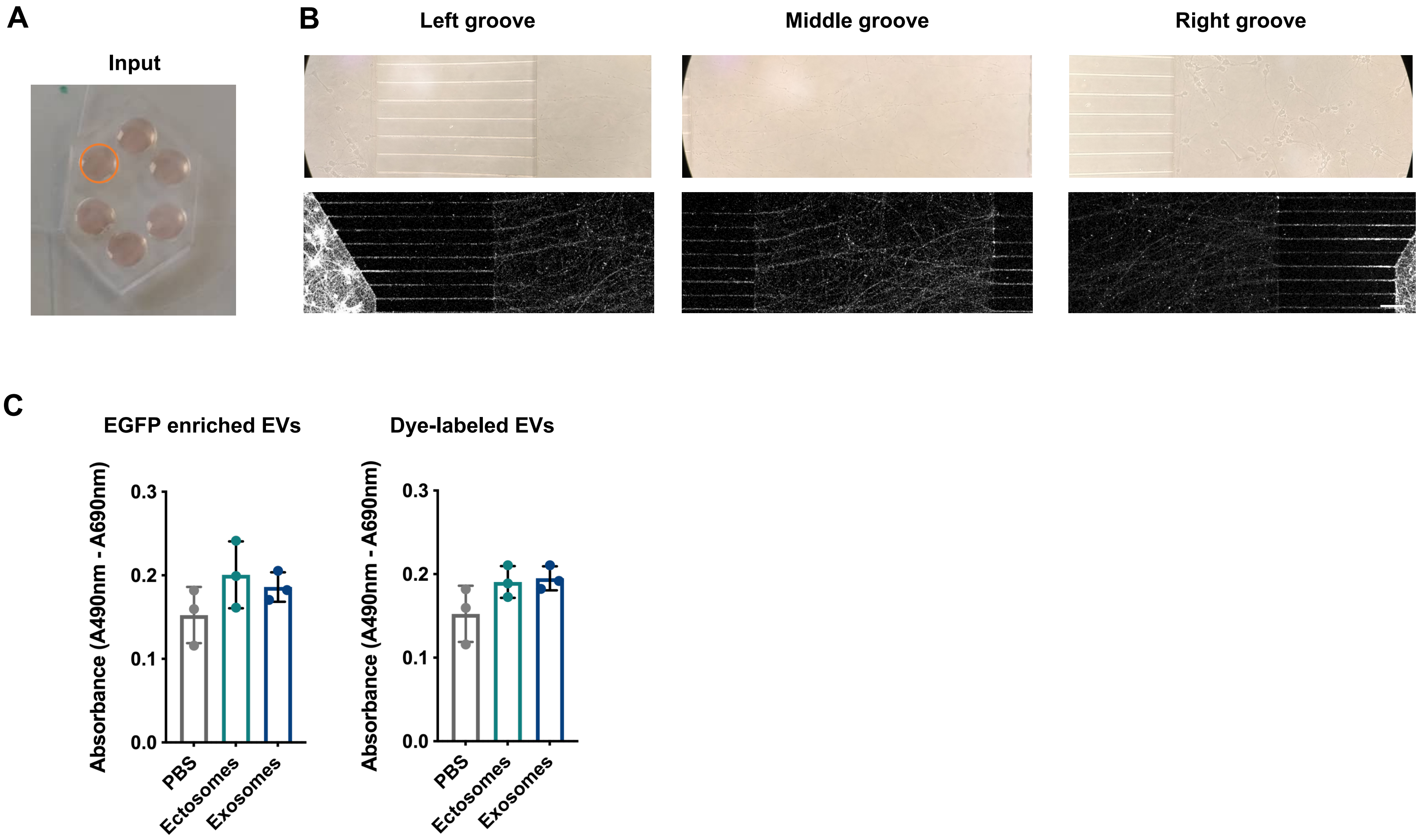


**Supplementary Figure 6. Cellular uptake of EVs by primary cortical neurons.**

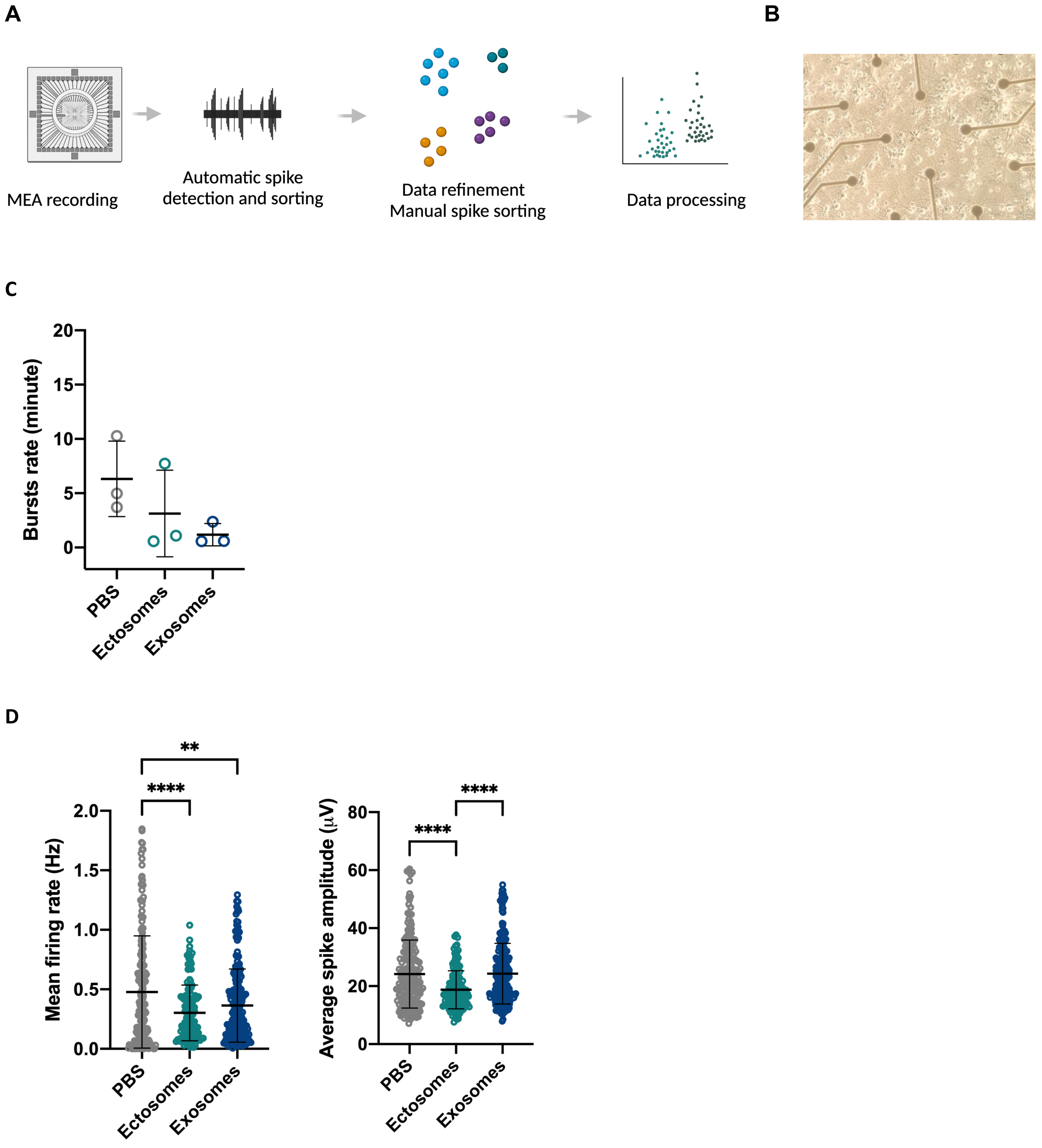


**Supplementary Figure 7. EVs modulate synaptic function in primary cortical neurons.**
