## Supplementary tables for "Ectosomes and exosomes are distinct proteomic entities that modulate spontaneous activity in neuronal cells"

**Supplementary Table 1.** List of significantly altered proteins identified in exosomes. Common proteins (11) in the list of Top 100 proteins often identified in EVs are highlighted in bold (see Supplementary Table 3).

| Protein names | Gene names | -log10(p-value) |
|---|---|---|
| **Alix** | ***PDCD6IP*** | **4,836** |
| Glycogen phosphorylase, muscle form | *PYGM* | 4,300 |
| Proteasome subunit beta type-1 | *PSMB1* | 4,098 |
| Glycogen phosphorylase, liver form | *PYGL* | 4,097 |
| Proteasome subunit alpha type-7 | *PSMA7* | 3,984 |
| Proteasome subunit beta type-7 | *PSMB7* | 3,752 |
| Asporin | *ASPN* | 3,734 |
| Proteasome subunit alpha type-6 | *PSMA6* | 3,695 |
| Proteasome subunit beta type-5 | *PSMB5* | 3,652 |
| Homogentisate 1,2-dioxygenase | *HGD* | 3,611 |
| Receptor-type tyrosine-protein phosphatase | *PTPRM* | 3,610 |
| Vacuolar protein sorting-associated protein 28 homolog | *VPS28* | 3,525 |
| Proteasome subunit alpha type-2 | *PSMA2* | 3,511 |
| Proteasome subunit alpha type-4 | *PSMA4* | 3,460 |
| Proteasome subunit beta type-4 | *PSMB4* | 3,443 |
| Cation-independent mannose-6-phosphate receptor | *IGF2R* | 3,377 |
| Eukaryotic initiation factor 4A-II | *EIF4A2* | 3,345 |
| **Protein kinase C-binding protein NELL2** | *NELL2* | 3,340 |
| 3-mercaptopyruvate sulfurtransferase | *MPST* | 3,327 |
| Mannose-1-phosphate guanyltransferase beta | *GMPPB* | 3,264 |
| Cytosolic 10-formyltetrahydrofolate dehydrogenase | *ALDH1L1* | 3,226 |
| Cytoplasmic aconitate hydratase | *ACO1* | 3,172 |
| S-adenosylmethionine synthase isoform type-1 | *MAT1A* | 3,097 |
| Delta-aminolevulinic acid dehydratase | *ALAD* | 3,039 |
| Tubulin alpha-4A chain | *TUBA4A* | 3,033 |
| Vacuolar protein sorting-associated protein 37B | *VPS37B* | 3,012 |
| **Adenosylhomocysteinase** | ***AHCY*** | **2,989** |
| Regucalcin | *RGN* | 2,989 |
| Fructose-bisphosphate aldolase B | *ALDOB* | 2,989 |
| 4-trimethylaminobutyraldehyde dehydrogenase | *ALDH9A1* | 2,960 |
| Microtubule-associated protein RP/EB family member 2 | *MAPRE2* | 2,933 |
| Uroporphyrinogen decarboxylase | *UROD* | 2,927 |
| Coronin-1A | *CORO1A* | 2,918 |
| Serine protease 23 | *PRSS23* | 2,896 |
| Proteasome subunit beta type-2 | *PSMB2* | 2,860 |
| Haptoglobin | *HP* | 2,859 |
| Collagen alpha-3(VI) chain | *COL6A3* | 2,814 |
| Argininosuccinate synthase | *ASS1* | 2,791 |
| **Galectin-3-binding protein** | ***LGALS3BP*** | **2,774** |
| **von Willebrand factor;von Willebrand antigen 2** | *VWF* | 2,736 |
| Lipoprotein lipase | *LPL* | 2,730 |
| Protein MON2 homolog | *MON2* | 2,712 |
| Coagulation factor X | *F10* | 2,700 |

| | | |
|---|---|---|
| 3-oxo-5-beta-steroid 4-dehydrogenase | *AKR1D1* | 2,696 |
| Proteasome subunit alpha type-5 | *PSMA5* | 2,660 |
| Reelin | *RELN* | 2,654 |
| Proteasome subunit alpha type-3 | *PSMA3* | 2,636 |
| Prostaglandin reductase 1 | *PTGR1* | 2,592 |
| Complement C4-A | *C4A;C4B* | 2,588 |
| Prostamide/prostaglandin F synthase | *FAM213B* | 2,553 |
| Collectin-10 | *COLEC10* | 2,547 |
| 26S proteasome non-ATPase regulatory subunit 5 | *PSMD5* | 2,523 |
| Carbonic anhydrase 2 | *CA2* | 2,519 |
| Ferritin heavy chain | *FTH1* | 2,515 |
| COP9 signalosome complex subunit 7b | *COPS7B* | 2,509 |
| Clusterin | *CLU* | 2,506 |
| COP9 signalosome complex subunit 3 | *COPS3* | 2,467 |
| **Syntenin-1** | ***SDCBP*** | **2,464** |
| Immunoglobulin J chain | *IGJ* | 2,463 |
| Retinal dehydrogenase 1 | *ALDH1A1* | 2,440 |
| Ectonucleotide pyrophosphatase/phosphodiesterase family member 2 | *ENPP2* | 2,412 |
| Alpha-crystallin B chain | *CRYAB* | 2,407 |
| Integrin-linked protein kinase | *ILK* | 2,402 |
| Ubiquitin thioesterase OTUB1 | *OTUB1* | 2,375 |
| Tenascin | *TNC* | 2,365 |
| Neuroserpin | *SERPINI1* | 2,360 |
| Quinone oxidoreductase | *CRYZ* | 2,330 |
| Prolow-density lipoprotein receptor-related protein 1 | *LRP1* | 2,328 |
| Proteasome subunit beta type-6 | *PSMB6* | 2,324 |
| Membrane-bound transcription factor site-1 protease | *MBTPS1* | 2,321 |
| Proteasome subunit alpha type-1 | *PSMA1* | 2,316 |
| General vesicular transport factor p115 | *USO1* | 2,311 |
| Alcohol dehydrogenase class-3 | *ADH5* | 2,301 |
| COP9 signalosome complex subunit 4 | *COPS4* | 2,294 |
| Sulfotransferase 1C2 | *SULT1C2* | 2,293 |
| Thyroglobulin | *TG* | 2,282 |
| Ras-related protein Rab-5B | *RAB5B* | 2,269 |
| Collagen alpha-1(II) chain | *COL2A1* | 2,246 |
| Tetraspanin-14 | *TSPAN14* | 2,231 |
| Betaine--homocysteine S-methyltransferase 1 | *BHMT* | 2,218 |
| Collagen alpha-2(V) chain | *COL5A2* | 2,204 |
| Thrombospondin-4 | *THBS4* | 2,203 |
| Proteasome subunit beta type-3 | *PSMB3* | 2,197 |
| Nicotinate-nucleotide pyrophosphorylase [carboxylating] | *QPRT* | 2,184 |
| Golgi membrane protein 1 | *GOLM1* | 2,177 |
| Flavin reductase (NADPH) | *BLVRB* | 2,156 |
| 26S protease regulatory subunit 6A | *PSMC3* | 2,154 |
| Phenazine biosynthesis-like domain-containing protein | *PBLD* | 2,146 |
| Glycogen phosphorylase, brain form | *PYGB* | 2,135 |
| Galactokinase | *GALK1* | 2,130 |
| Follistatin-related protein 1 | *FSTL1* | 2,130 |

| | | |
|---|---|---|
| Aflatoxin B1 aldehyde reductase member 4 | *AKR7L* | 2,122 |
| Actin-related protein 2/3 complex subunit 1B | *ARPC1B* | 2,120 |
| Ribonucleoside-diphosphate reductase large subunit | *RRM1* | 2,083 |
| Protocadherin Fat 1 | *FAT1* | 2,074 |
| Septin-9 | *09-Sep* | 2,074 |
| 4-hydroxyphenylpyruvate dioxygenase | *HPD* | 2,068 |
| Complement C1q tumor necrosis factor-related protein 3 | *C1QTNF3* | 2,056 |
| Vesicular integral-membrane protein VIP36 | *LMAN2* | 2,020 |
| Procollagen-lysine,2-oxoglutarate 5-dioxygenase 1 | *PLOD1* | 1,988 |
| Collagen alpha-1(I) chain | *COL1A1* | 1,949 |
| Fumarylacetoacetate hydrolase domain-containing protein 2A | *FAHD2A;FAHD2B* | 1,929 |
| Ras-related protein Rab-1B | *RAB1B* | 1,910 |
| Dehydrogenase/reductase SDR family member 11 | *DHRS11* | 1,893 |
| Collagen alpha-2(I) chain | *COL1A2* | 1,880 |
| **GTP-binding nuclear protein Ran** | ***RAN*** | **1,877** |
| Collagen alpha-1(V) chain | *COL5A1* | 1,874 |
| Metalloproteinase inhibitor 2 | *TIMP2* | 1,870 |
| Tetraspanin-6 | *TSPAN6* | 1,852 |
| Serine/threonine-protein phosphatase 2A catalytic subunit alpha isoform | *PPP2CA* | 1,843 |
| Mitogen-activated protein kinase 1 | *MAPK1* | 1,822 |
| Endothelial protein C receptor | *PROCR* | 1,815 |
| Isocitrate dehydrogenase [NADP] cytoplasmic | *IDH1* | 1,811 |
| Membrane primary amine oxidase | *AOC3* | 1,810 |
| Low molecular weight phosphotyrosine protein phosphatase | *ACP1* | 1,803 |
| Bone morphogenetic protein 1 | *BMP1* | 1,796 |
| 60S ribosomal protein L12 | *RPL12* | 1,784 |
| Adenosine kinase | *ADK* | 1,783 |
| Beta-parvin | *PARVB* | 1,779 |
| Peroxidasin homolog | *PXDN* | 1,775 |
| Arginase-1 | *ARG1* | 1,764 |
| Dihydropyrimidinase | *DPYS* | 1,763 |
| Fibrinogen gamma chain | *FGG* | 1,745 |
| Tetraspanin-9 | *TSPAN9* | 1,742 |
| Transforming growth factor beta-1 | *TGFB1* | 1,740 |
| Fibrocystin-L | *PKHD1L1* | 1,737 |
| Glutaminyl-peptide cyclotransferase | *QPCT* | 1,720 |
| Fumarylacetoacetase | *FAH* | 1,715 |
| Arrestin domain-containing protein 1 | *ARRDC1* | 1,706 |
| Probable maltase-glucoamylase-like protein | *LOC93432* | 1,703 |
| V-type proton ATPase catalytic subunit A | *ATP6V1A* | 1,700 |
| Alpha-galactosidase A | *GLA* | 1,698 |
| Probable imidazolonepropionase | *AMDHD1* | 1,690 |
| ProSAAS | *PCSK1N* | 1,685 |
| Cerebellin-4 | *CBLN4* | 1,685 |
| Receptor-type tyrosine-protein phosphatase kappa | *PTPRK* | 1,671 |
| Collagen alpha-2(VI) chain | *COL6A2* | 1,671 |
| Lysosomal alpha-mannosidase | *MAN2B1* | 1,666 |
| Cathepsin L2 | *CTSV* | 1,659 |

| | | |
|---|---|---|
| Glucosamine 6-phosphate N-acetyltransferase | *GNPNAT1* | 1,649 |
| Isopentenyl-diphosphate Delta-isomerase 1 | *IDI1* | 1,636 |
| Vesicle-associated membrane protein 2 | *VAMP2* | 1,635 |
| cGMP-specific 3,5-cyclic phosphodiesterase | *PDE5A* | 1,630 |
| cAMP-dependent protein kinase type I-alpha regulatory subunit | *PRKAR1A* | 1,624 |
| Acetyl-CoA carboxylase 1 | *ACACA* | 1,620 |
| Alpha-mannosidase 2 | *MAN2A1* | 1,619 |
| Heat shock protein beta-1 | *HSPB1* | 1,616 |
| Extracellular matrix protein FRAS1 | *FRAS1* | 1,607 |
| **CD63 antigen** | ***CD63*** | **1,589** |
| MMS19 nucleotide excision repair protein homolog | *MMS19* | 1,583 |
| Apolipoprotein C-III | *APOC3* | 1,580 |
| 40S ribosomal protein S20 | *RPS20* | 1,578 |
| L-xylulose reductase | *DCXR* | 1,556 |
| Fermitin family homolog 3 | *FERMT3* | 1,552 |
| Prolyl endopeptidase FAP | *FAP* | 1,545 |
| Estrogen sulfotransferase | *SULT1E1* | 1,544 |
| Sphingomyelin phosphodiesterase | *SMPD1* | 1,538 |
| Ras-related C3 botulinum toxin substrate 2 | *RAC2* | 1,536 |
| 40S ribosomal protein SA | *RPSA* | 1,529 |
| Sorbitol dehydrogenase | *SORD* | 1,516 |
| Vacuolar protein-sorting-associated protein 25 | *VPS25* | 1,496 |
| Protein CutA | *CUTA* | 1,485 |
| 26S proteasome non-ATPase regulatory subunit 14 | *PSMD14* | 1,483 |
| C-type lectin domain family 11 member A | *CLEC11A* | 1,483 |
| **Talin-1** | ***TLN1*** | **1,479** |
| **Gelsolin** | ***GSN*** | **1,477** |
| COP9 signalosome complex subunit 2 | *COPS2* | 1,474 |
| Putative RNA-binding protein Luc7-like 2 | *LUC7L2* | 1,472 |
| 40S ribosomal protein S17 | *RPS17* | 1,470 |
| **CD81 antigen** | ***CD81*** | **1,454** |
| Transmembrane protein 132C | *TMEM132C* | 1,444 |
| Nuclear transport factor 2 | *NUTF2* | 1,441 |
| Inositol-3-phosphate synthase 1 | *ISYNA1* | 1,428 |
| 26S proteasome non-ATPase regulatory subunit 7 | *PSMD7* | 1,421 |
| Cochlin | *COCH* | 1,410 |
| ER lumen protein-retaining receptor 1 | *KDELR1* | 1,410 |
| ATP-dependent 6-phosphofructokinase, platelet type | *PFKP* | 1,398 |
| Carboxymethylenebutenolidase homolog | *CMBL* | 1,391 |
| 26S proteasome non-ATPase regulatory subunit 11 | *PSMD11* | 1,388 |
| Deleted in malignant brain tumors 1 protein | *DMBT1* | 1,387 |
| Pleckstrin homology domain-containing family B member 2 | *PLEKHB2* | 1,360 |
| Gamma-glutamyl hydrolase | *GGH* | 1,349 |
| **Complement C3** | ***C3*** | **1,344** |
| LIM and senescent cell antigen-like-containing domain protein 1 | *LIMS1* | 1,338 |
| Complement C1r subcomponent | *C1R* | 1,324 |
| Desmocollin-3 | *DSC3* | 1,321 |
| COP9 signalosome complex subunit 6 | *COPS6* | 1,316 |

| | | |
|---|---|---|
| 26S proteasome non-ATPase regulatory subunit 3 | *PSMD3* | 1,312 |
| Bisphosphoglycerate mutase | *BPGM* | 1,301 |
| Galectin-related protein | *LGALSL* | 1,282 |
| Cathepsin Z | *CTSZ* | 1,275 |
| Tetraspanin-7 | *TSPAN7* | 1,272 |
| Multiple epidermal growth factor-like domains protein 10 | *MEGF10* | 1,242 |
| EMILIN-3 | *EMILIN3* | 1,232 |
| Vigilin | *HDLBP* | 1,225 |
| Serine/threonine-protein phosphatase CPPED1 | *CPPED1* | 1,189 |
| Collagen alpha-1(XII) chain | *COL12A1* | 1,181 |
| **Alpha-2-macroglobulin** | ***A2M*** | **1,112** |
| **Ig gamma-4 chain C region** | *IGHG4* | 1,089 |
| Cornulin | *CRNN* | 1,037 |
| Hemoglobin subunit beta | *HBB* | 1,016 |

**Supplementary Table 2.** List of significantly altered proteins identified in ectosomes. Common proteins (52) in the list of Top 100 proteins often identified in EVs are highlighted in bold (see Supplementary Table 3).

| **Protein names** | **Gene names** | **-log10(p-value)** |
|---|---|---:|
| Band 4.1-like protein 2 | *EPB41L2* | 9,852 |
| Band 4.1-like protein 3 | *EPB41L3* | 8,468 |
| **Plasma membrane calcium-transporting ATPase 1** | *ATP2B1* | 7,567 |
| Phosphatidylinositol 4-kinase alpha | *PI4KA* | 7,239 |
| Tubulin-specific chaperone A | *TBCA* | 6,731 |
| **Profilin-1** | ***PFN1*** | **6,708** |
| MARCKS-related protein | *MARCKSL1* | 6,644 |
| **4F2 cell-surface antigen heavy chain** | ***SLC3A2*** | **6,412** |
| Cytoplasmic FMR1-interacting protein 1 | *CYFIP1* | 6,209 |
| Myelin protein zero-like protein 1 | *MPZL1* | 6,167 |
| Na(+)/H(+) exchange regulatory cofactor NHE-RF1 | *SLC9A3R1* | 6,151 |
| Unconventional myosin-Ic | *MYO1C* | 5,986 |
| Protein scribble homolog | *SCRIB* | 5,985 |
| N(G),N(G)-dimethylarginine dimethylaminohydrolase 1 | *DDAH1* | 5,900 |
| 2,3-cyclic-nucleotide 3-phosphodiesterase | *CNP* | 5,696 |
| **Sodium/potassium-transporting ATPase subunit alpha-1** | ***ATP1A1*** | **5,640** |
| Lethal(2) giant larvae protein homolog 1 | *LLGL1* | 5,620 |
| FERM, RhoGEF and pleckstrin domain-containing protein 1 | *FARP1* | 5,604 |
| Integrin alpha-5 | *ITGA5* | 5,589 |
| Catenin delta-1 | *CTNND1* | 5,556 |
| Charged multivesicular body protein 6 | *CHMP6* | 5,507 |
| Basigin | *BSG* | 5,349 |
| PDZ domain-containing protein GIPC1 | *GIPC1* | 5,304 |
| Plastin-3 | *PLS3* | 5,244 |
| Peripheral plasma membrane protein CASK | *CASK* | 5,235 |
| Trifunctional purine biosynthetic protein adenosine-3 | *GART* | 5,214 |
| **14-3-3 protein epsilon** | ***YWHAE*** | **5,213** |
| **Moesin** | ***MSN*** | **5,169** |

| | | |
|---|---|---|
| **Annexin A6** | ***ANXA6*** | **5,150** |
| Myristoylated alanine-rich C-kinase substrate | *MARCKS* | 5,105 |
| Protein lin-7 homolog C | *LIN7C* | 5,092 |
| Guanine nucleotide-binding protein subunit alpha-13 | *GNA13* | 5,054 |
| **Ubiquitin-40S ribosomal protein S27a** | *RPS27A;UBB/C* | 5,027 |
| Immunoglobulin superfamily member 3 | *IGSF3* | 4,988 |
| EGF-like repeat and discoidin I-like domain-containing protein 3 | *EDIL3* | 4,974 |
| Sodium/potassium-transporting ATPase subunit beta-1 | *ATP1B1* | 4,960 |
| Unconventional myosin-Ib | *MYO1B* | 4,885 |
| WD repeat-containing protein 6 | *WDR6* | 4,826 |
| Guanine nucleotide-binding protein G(i) subunit alpha-1 | *GNAI1* | 4,814 |
| Inactive tyrosine-protein kinase 7 | *PTK7* | 4,790 |
| CD276 antigen | *CD276* | 4,781 |
| Sodium/potassium-transporting ATPase subunit beta-3 | *ATP1B3* | 4,738 |
| Catenin beta-1 | *CTNNB1* | 4,699 |
| Unconventional myosin-Id | *MYO1D* | 4,582 |
| Calcyclin-binding protein | *CACYBP* | 4,579 |
| Heat shock 70 kDa protein 4 | *HSPA4* | 4,550 |
| Guanine nucleotide-binding protein G(I)/G(S)/G(O) subunit gamma-12 | *GNG12* | 4,474 |
| Afadin | *MLLT4* | 4,449 |
| Catenin alpha-1 | *CTNNA1* | 4,449 |
| Ras-related protein Rab-21 | *RAB21* | 4,444 |
| Kinesin-like protein KIF23 | *KIF23* | 4,376 |
| A-kinase anchor protein 12 | *AKAP12* | 4,302 |
| Complement component 1 Q subcomponent-binding protein | *C1QBP* | 4,296 |
| **Peroxiredoxin-1** | ***PRDX1*** | **4,263** |
| Neutral amino acid transporter B(0) | *SLC1A5* | 4,221 |
| **Fatty acid synthase** | ***FASN*** | **4,138** |
| Pachytene checkpoint protein 2 homolog | *TRIP13* | 4,113 |
| Radixin | *RDX* | 4,089 |
| Phospholipid-transporting ATPase IG | *ATP11C* | 4,077 |
| **Guanine nucleotide-binding protein G(s) subunit alpha isoforms** | ***GNAS*** | **4,055** |
| Nck-associated protein 1 | *NCKAP1* | 4,054 |
| C-1-tetrahydrofolate synthase | *MTHFD1* | 4,019 |
| Disks large homolog 1 | *DLG1* | 4,010 |
| Glutathione S-transferase P | *GSTP1* | 4,008 |
| Creatine kinase B-type | *CKB* | 4,007 |
| Ectonucleotide pyrophosphatase/phosphodiesterase family member 1 | *ENPP1* | 3,995 |
| Collagen alpha-1(XVIII) chain | *COL18A1* | 3,995 |
| Integrin alpha-1 | *ITGA1* | 3,987 |
| **14-3-3 protein beta/alpha** | ***YWHAB*** | **3,953** |
| Cyclin-dependent kinase 1 | *CDK1* | 3,950 |
| Zinc transporter ZIP6 | *SLC39A6* | 3,947 |
| Microtubule-associated protein RP/EB family member 1 | *MAPRE1* | 3,914 |
| Large neutral amino acids transporter small subunit 1 | *SLC7A5* | 3,865 |
| Translation initiation factor eIF-2B subunit alpha | *EIF2B1* | 3,790 |
| Monocarboxylate transporter 1 | *SLC16A1* | 3,788 |
| **14-3-3 protein theta** | ***YWHAQ*** | **3,784** |

| | | |
|---|---|---|
| **Ras GTPase-activating-like protein** | ***IQGAP1*** | **3,751** |
| **14-3-3 protein eta** | ***YWHAH*** | **3,743** |
| Peroxiredoxin-6 | *PRDX6* | 3,729 |
| Cytoplasmic FMR1-interacting protein 2 | *CYFIP2* | 3,687 |
| Kinesin-1 heavy chain | *KIF5B* | 3,680 |
| Nicastrin | *NCSTN* | 3,677 |
| Guanine nucleotide-binding protein G(k) subunit alpha | *GNAI3* | 3,644 |
| **Annexin A2** | ***ANXA2*** | **3,628** |
| dCTP pyrophosphatase 1 | *DCTPP1* | 3,580 |
| **Integrin beta-1** | ***ITGB1*** | **3,579** |
| DnaJ homolog subfamily C member 5 | *DNAJC5* | 3,575 |
| Four and a half LIM domains protein 1 | *FHL1* | 3,567 |
| Actin, cytoplasmic 2 | *ACTG1* | 3,555 |
| Endoplasmin | *HSP90B1* | 3,554 |
| BolA-like protein 2 | *BOLA2* | 3,550 |
| Purine nucleoside phosphorylase | *PNP* | 3,538 |
| Basement membrane-specific heparan sulfate proteoglycan core protein | *HSPG2* | 3,487 |
| Receptor-type tyrosine-protein phosphatase F | *PTPRF* | 3,433 |
| D-3-phosphoglycerate dehydrogenase | *PHGDH* | 3,433 |
| Protein unc-45 homolog A | *UNC45A* | 3,432 |
| Insulin receptor substrate 4 | *IRS4* | 3,430 |
| **Guanine nucleotide-binding protein G(i) subunit alpha-2** | ***GNAI2*** | **3,417** |
| Brain acid soluble protein 1 | *BASP1* | 3,383 |
| **Ezrin** | ***EZR*** | **3,375** |
| Cofilin-2 | *CFL2* | 3,371 |
| 10 kDa heat shock protein, mitochondrial | *HSPE1* | 3,363 |
| Protein NDRG1 | *NDRG1* | 3,352 |
| Coxsackievirus and adenovirus receptor | *CXADR* | 3,347 |
| Poly(rC)-binding protein 2 | *PCBP2* | 3,321 |
| **Ubiquitin carboxyl-terminal hydrolase isozyme L1** | *UCHL1* | 3,321 |
| Guanine nucleotide-binding protein subunit alpha-11 | *GNA11* | 3,303 |
| **Heat shock protein HSP 90-beta** | ***HSP90AB1*** | **3,294** |
| Heat shock protein 105 kDa | *HSPH1* | 3,259 |
| **Ras-related protein Rab-10** | ***RAB10*** | **3,255** |
| GTP-binding protein Rheb | *RHEB* | 3,252 |
| Protein XRP2 | *RP2* | 3,245 |
| Equilibrative nucleoside transporter 1 | *SLC29A1* | 3,219 |
| Protein unc-13 homolog D | *UNC13D* | 3,212 |
| **14-3-3 protein zeta/delta** | ***YWHAZ*** | **3,209** |
| Poly(rC)-binding protein 1 | *PCBP1* | 3,208 |
| Ephrin type-A receptor 2 | *EPHA2* | 3,206 |
| Plexin-A1 | *PLXNA1* | 3,205 |
| Heat shock 70 kDa protein 1B;Heat shock 70 kDa protein 1A | *HSPA1B;HSPA1A* | 3,197 |
| Hypoxanthine-guanine phosphoribosyltransferase | *HPRT1* | 3,188 |
| Programmed cell death protein 5 | *PDCD5* | 3,183 |
| **T-complex protein 1 subunit alpha** | ***TCP1*** | **3,167** |
| **Annexin A1** | ***ANXA1*** | **3,162** |
| Prohibitin | *PHB* | 3,152 |

| | | |
|---|---|---|
| Chloride intracellular channel protein 4 | *CLIC4* | 3,144 |
| Leucine-rich repeat-containing protein 57 | *LRRC57* | 3,142 |
| Neurogenic locus notch homolog protein 2 | *NOTCH2* | 3,139 |
| **Ras-related protein Rab-7a** | ***RAB7A*** | **3,137** |
| Junctional adhesion molecule C | *JAM3* | 3,108 |
| Nidogen-2 | *NID2* | 3,103 |
| Small VCP/p97-interacting protein | *SVIP* | 3,093 |
| MOB kinase activator 1A;MOB kinase activator 1B | *MOB1A;MOB1B* | 3,090 |
| Inorganic pyrophosphatase | *PPA1* | 3,088 |
| Ras-related protein Rab-23 | *RAB23* | 3,071 |
| Zinc finger CCCH-type antiviral protein 1 | *ZC3HAV1* | 3,039 |
| HLA class I histocompatibility antigen, A-2 alpha chain | *HLA-A* | 3,034 |
| Glypican-4;Secreted glypican-4 | *GPC4* | 3,033 |
| Phosphoribosylformylglycinamidine synthase | *PFAS* | 2,977 |
| **L-lactate dehydrogenase A chain** | ***LDHA*** | **2,965** |
| T-complex protein 1 subunit epsilon | *CCT5* | 2,944 |
| Single-stranded DNA-binding protein, mitochondrial | *SSBP1* | 2,938 |
| HLA class I histocompatibility antigen, B-7 alpha chain | *HLA-B* | 2,909 |
| Syntaxin-4 | *STX4* | 2,907 |
| Integrin alpha-V | *ITGAV* | 2,885 |
| Synaptosomal-associated protein 23 | *SNAP23* | 2,884 |
| Fascin | *FSCN1* | 2,883 |
| **Peroxiredoxin-2** | ***PRDX2*** | **2,876** |
| **14-3-3 protein gamma** | ***YWHAG*** | **2,874** |
| **Guanine nucleotide-binding protein G(I)/G(S)/G(T) subunit beta-1** | ***GNB1*** | **2,860** |
| Rho GDP-dissociation inhibitor 1 | *ARHGDIA* | 2,851 |
| Copine-3 | *CPNE3* | 2,845 |
| **Triosephosphate isomerase** | ***TPI1*** | **2,842** |
| ADP-sugar pyrophosphatase | *NUDT5* | 2,839 |
| Solute carrier family 12 member 2 | *SLC12A2* | 2,802 |
| **Elongation factor 1-alpha 1;Putative elongation factor 1-alpha-like 3** | ***EEF1A1;EEF1A1P5*** | **2,761** |
| Synaptic vesicle membrane protein VAT-1 homolog | *VAT1* | 2,731 |
| T-complex protein 1 subunit eta | *CCT7* | 2,729 |
| **T-complex protein 1 subunit zeta** | ***CCT6A*** | **2,728** |
| Serine/threonine-protein kinase MRCK beta | *CDC42BPB* | 2,699 |
| Multifunctional protein ADE2 | *PAICS* | 2,688 |
| Protein FAM49B | *FAM49B* | 2,686 |
| **Ras-related protein Rab-5C** | ***RAB5C*** | **2,680** |
| Receptor-type tyrosine-protein phosphatase gamma | *PTPRG* | 2,677 |
| Myosin light polypeptide 6 | *MYL6* | 2,673 |
| Destrin | *DSTN* | 2,673 |
| **Fibronectin** | ***FN1*** | **2,672** |
| Gap junction alpha-1 protein | *GJA1* | 2,665 |
| LIM and SH3 domain protein 1 | *LASP1* | 2,647 |
| Aspartate aminotransferase, cytoplasmic | *GOT1* | 2,643 |
| CTP synthase 1 | *CTPS1* | 2,640 |
| **Guanine nucleotide-binding protein G(I)/G(S)/G(T) subunit beta-2** | ***GNB2*** | **2,631** |

| | | |
|---|---|---|
| Collagen alpha-1(IV) chain | *COL4A1* | 2,630 |
| Ras-related protein Rab-8B | *RAB8B* | 2,629 |
| Elongation factor 1-delta | *EEF1D* | 2,609 |
| **T-complex protein 1 subunit delta** | ***CCT4*** | **2,597** |
| Serine/threonine-protein kinase 26 | *STK26* | 2,584 |
| 60 kDa heat shock protein, mitochondrial | *HSPD1* | 2,582 |
| Integrin alpha-2 | *ITGA2* | 2,567 |
| Disks large homolog 3 | *DLG3* | 2,562 |
| Calcineurin B homologous protein 1 | *CHP1* | 2,549 |
| Ras-related protein Rab-35 | *RAB35* | 2,518 |
| Protein disulfide-isomerase A6 | *PDIA6* | 2,517 |
| **Ras-related protein Ral-A** | ***RALA*** | **2,509** |
| Elongation factor 1-beta | *EEF1B2* | 2,506 |
| Nuclear migration protein nudC | *NUDC* | 2,505 |
| Mitogen-activated protein kinase kinase kinase kinase 4 | *MAP4K4* | 2,487 |
| Choline transporter-like protein 2 | *SLC44A2* | 2,486 |
| Insulin-like growth factor 1 receptor | *IGF1R* | 2,486 |
| **Glyceraldehyde-3-phosphate dehydrogenase** | ***GAPDH*** | **2,478** |
| **Ras-related protein R-Ras2** | *RRAS2* | 2,478 |
| Protein deglycase DJ-1 | *PARK7* | 2,444 |
| Coatomer subunit alpha;Xenin;Proxenin | *COPA* | 2,433 |
| **Importin subunit beta-1** | ***KPNB1*** | **2,432** |
| **T-complex protein 1 subunit beta** | ***CCT2*** | **2,423** |
| Ras GTPase-activating protein 3 | *RASA3* | 2,405 |
| Importin-4 | *IPO4* | 2,393 |
| Presenilin-1 | *PSEN1* | 2,393 |
| 2-deoxynucleoside 5-phosphate N-hydrolase 1 | *DNPH1* | 2,385 |
| Prohibitin-2 | *PHB2* | 2,378 |
| **Heat shock protein HSP 90-alpha** | ***HSP90AA1*** | **2,376** |
| Elongation factor Tu, mitochondrial | *TUFM* | 2,363 |
| Glycine-tRNA ligase | *GARS* | 2,356 |
| GTPase HRas | *HRAS* | 2,355 |
| **Pyruvate kinase PKM** | ***PKM*** | **2,355** |
| Guanine nucleotide-binding protein G(o) subunit alpha | *GNAO1* | 2,351 |
| Rac GTPase-activating protein 1 | *RACGAP1* | 2,343 |
| ATP-dependent RNA helicase DDX3X/ DDX3Y | *DDX3X;DDX3Y* | 2,341 |
| Zinc transporter ZIP14 | *SLC39A14* | 2,336 |
| Sorting nexin-5 | *SNX5* | 2,330 |
| Ubiquitin-conjugating enzyme E2 K | *UBE2K* | 2,325 |
| Thioredoxin domain-containing protein 17 | *TXNDC17* | 2,317 |
| **Cofilin-1** | ***CFL1*** | **2,313** |
| Heterogeneous nuclear ribonucleoprotein A3 | *HNRNPA3* | 2,310 |
| DNA topoisomerase 2-alpha | *TOP2A* | 2,308 |
| Calcium-binding protein 39 | *CAB39* | 2,290 |
| Collagen alpha-2(IV) chain;Canstatin | *COL4A2* | 2,285 |
| Lactoylglutathione lyase | *GLO1* | 2,281 |
| Rho-associated protein kinase 1 | *ROCK1* | 2,271 |
| Centrosomal protein of 55 kDa | *CEP55* | 2,268 |

| | | |
|---|---|---|
| Nucleoside diphosphate kinase B;Putative nucleoside diphosphate kinase | *NME2;NME2P1* | 2,255 |
| **Ras-related C3 botulinum toxin substrate 1** | ***RAC1*** | **2,246** |
| Neuropilin-1 | *NRP1* | 2,227 |
| Phosphatidylethanolamine-binding protein 1; **Hippocampal cholinergic neurostimulating peptide** | *PEBP1* | 2,217 |
| Integrin alpha-6 | *ITGA6* | 2,203 |
| Protein kinase C beta type | *PRKCB* | 2,196 |
| Fatty acid-binding protein, epidermal | *FABP5* | 2,190 |
| **T-complex protein 1 subunit theta** | ***CCT8*** | **2,180** |
| Endophilin-B1 | *SH3GLB1* | 2,168 |
| AP-2 complex subunit beta | *AP2B1* | 2,147 |
| Melanoma-associated antigen D2 | *MAGED2* | 2,127 |
| Stathmin | *STMN1* | 2,123 |
| Annexin A3 | *ANXA3* | 2,100 |
| Fructose-bisphosphate aldolase A | *ALDOA* | 2,100 |
| **Heat shock cognate 71 kDa protein** | ***HSPA8*** | **2,096** |
| **Alpha-enolase** | ***ENO1*** | **2,093** |
| Guanine nucleotide-binding protein G(q) subunit alpha | *GNAQ* | 2,090 |
| Phosphatidylinositol transfer protein beta isoform | *PITPNB* | 2,089 |
| Junction plakoglobin | *JUP* | 2,075 |
| Filamin-B | *FLNB* | 2,072 |
| Golgin subfamily A member 7 | *GOLGA7* | 2,071 |
| Programmed cell death protein 10 | *PDCD10* | 2,062 |
| Neuroblast differentiation-associated protein AHNAK | *AHNAK* | 2,045 |
| Puromycin-sensitive aminopeptidase | *NPEPPS* | 2,044 |
| Lysyl oxidase homolog 2 | *LOXL2* | 2,043 |
| Transgelin-2 | *TAGLN2* | 2,032 |
| Anosmin-1 | *KAL1* | 2,029 |
| Cyclin-Y | *CCNY* | 2,016 |
| Laminin subunit alpha-5 | *LAMA5* | 2,015 |
| Nucleoside diphosphate kinase A | *NME1* | 2,011 |
| Rho-related GTP-binding protein RhoG | *RHOG* | 2,008 |
| Serine/threonine-protein kinase TAO1 | *TAOK1* | 1,998 |
| Proliferating cell nuclear antigen | *PCNA* | 1,991 |
| BTB/POZ domain-containing protein KCTD12 | *KCTD12* | 1,977 |
| Staphylococcal nuclease domain-containing protein 1 | *SND1* | 1,963 |
| CD59 glycoprotein | *CD59* | 1,961 |
| Junctional adhesion molecule A | *F11R* | 1,959 |
| **Annexin A5** | ***ANXA5*** | **1,957** |
| Protein NDNF | *NDNF* | 1,956 |
| NADH-cytochrome b5 reductase 3 | *CYB5R3* | 1,953 |
| Ras-related protein Rab-13 | *RAB13* | 1,943 |
| Occludin | *OCLN* | 1,938 |
| Glypican-6;Secreted glypican-6 | *GPC6* | 1,933 |
| CAD protein | *CAD* | 1,909 |
| Lysosome-associated membrane glycoprotein 1 | *LAMP1* | 1,899 |
| **Protein-L-isoaspartate(D-aspartate) O-methyltransferase** | *PCMT1* | 1,898 |
| Activator of 90 kDa heat shock protein ATPase homolog 1 | *AHSA1* | 1,886 |

| | | |
|---|---|---|
| Fermitin family homolog 2 | *FERMT2* | 1,875 |
| Band 4.1-like protein 5 | *EPB41L5* | 1,870 |
| STE20-like serine/threonine-protein kinase | *SLK* | 1,869 |
| Stress-70 protein, mitochondrial | *HSPA9* | 1,863 |
| DnaJ homolog subfamily B member 1 | *DNAJB1* | 1,858 |
| Malate dehydrogenase, mitochondrial | *MDH2* | 1,847 |
| Protein EFR3 homolog A | *EFR3A* | 1,842 |
| ATP synthase subunit alpha, mitochondrial | *ATP5A1* | 1,827 |
| Agrin | *AGRN* | 1,821 |
| Voltage-dependent anion-selective channel protein 2 | *VDAC2* | 1,821 |
| Hsc70-interacting protein | *ST13* | 1,816 |
| F-actin-capping protein subunit beta | *CAPZB* | 1,810 |
| POTE ankyrin domain family member E / F | *POTEE;POTEF* | 1,798 |
| Gamma-aminobutyric acid receptor-associated protein-like 2 | *GABARAPL2* | 1,775 |
| Teneurin-3 | *TENM3* | 1,767 |
| Histone H1.0 | *H1F0* | 1,727 |
| **Ubiquitin-like modifier-activating enzyme 1** | ***UBA1*** | **1,726** |
| Eukaryotic translation initiation factor 2 subunit 1 | *EIF2S1* | 1,722 |
| Desmoglein-2 | *DSG2* | 1,721 |
| **Elongation factor 2** | ***EEF2*** | **1,717** |
| GTPase NRas | *NRAS* | 1,710 |
| ATP synthase subunit beta, mitochondrial | *ATP5B* | 1,708 |
| Aspartate--tRNA ligase, cytoplasmic | *DARS* | 1,700 |
| E3 ubiquitin-protein ligase CHIP | *STUB1* | 1,697 |
| Developmentally-regulated GTP-binding protein 1 | *DRG1* | 1,697 |
| **Protein-glutamine gamma-glutamyltransferase E** | *TGM3* | 1,692 |
| Tyrosine-protein kinase Lyn | *LYN* | 1,668 |
| **Clathrin heavy chain 1** | ***CLTC*** | **1,666** |
| Epidermal growth factor receptor | *EGFR* | 1,664 |
| Serine/threonine-protein phosphatase PP1-alpha catalytic subunit | *PPP1CA* | 1,663 |
| Thioredoxin-dependent peroxide reductase, mitochondrial | *PRDX3* | 1,658 |
| Malate dehydrogenase, cytoplasmic | *MDH1* | 1,655 |
| Syntaxin-7 | *STX7* | 1,638 |
| Inosine-5-monophosphate dehydrogenase 2 | *IMPDH2* | 1,635 |
| Ran-specific GTPase-activating protein | *RANBP1* | 1,630 |
| **78 kDa glucose-regulated protein** | ***HSPA5*** | **1,628** |
| **Cell division control protein 42 homolog** | ***CDC42*** | **1,607** |
| DnaJ homolog subfamily A member 2 | *DNAJA2* | 1,605 |
| **L-lactate dehydrogenase B chain** | ***LDHB*** | **1,599** |
| ADP/ATP translocase 2 | *SLC25A5* | 1,585 |
| Profilin-2 | *PFN2* | 1,580 |
| Phosphatidylinositol 5-phosphate 4-kinase type-2 alpha | *PIP4K2A* | 1,580 |
| Alanine--tRNA ligase, cytoplasmic | *AARS* | 1,560 |
| D-dopachrome decarboxylase;D-dopachrome decarboxylase-like protein | *DDT;DDTL* | 1,555 |
| Protein 4.1 | *EPB41* | 1,553 |
| Histone H2B type 1-C/E/F/G/I | *HIST1H2BC* | 1,537 |
| Long-chain-fatty-acid--CoA ligase 4 | *ACSL4* | 1,535 |
| **Rab GDP dissociation inhibitor beta** | ***GDI2*** | **1,524** |

| | | |
|---|---|---|
| Phosphoribosyl pyrophosphate synthase-associated protein 2 | *PRPSAP2* | 1,523 |
| Ras-related protein Rap-2b | *RAP2B* | 1,517 |
| Choline transporter-like protein 1 | *SLC44A1* | 1,516 |
| NudC domain-containing protein 2 | *NUDCD2* | 1,513 |
| Plasma membrane calcium-transporting ATPase 4 | *ATP2B4* | 1,504 |
| Ras-related protein Rab-8A | *RAB8A* | 1,503 |
| Axin interactor, dorsalization-associated protein | *AIDA* | 1,492 |
| High affinity cationic amino acid transporter 1 | *SLC7A1* | 1,492 |
| N-alpha-acetyltransferase 15, NatA auxiliary subunit | *NAA15* | 1,488 |
| Histone H1.4 | *HIST1H1E* | 1,469 |
| Translational activator GCN1 | *GCN1L1* | 1,462 |
| Caspase-3;Caspase-3 subunit p17;Caspase-3 subunit p12 | *CASP3* | 1,458 |
| Proliferation-associated protein 2G4 | *PA2G4* | 1,458 |
| Histone H2A type 1C;Histone H2A type 3;Histone H2A type 1-B/E | *HIST1H2AC;HIST 3H2A;HIST1H2AB* | 1,454 |
| Eukaryotic peptide chain release factor subunit 1 | *ETF1* | 1,436 |
| Ephrin type-A receptor 7 | *EPHA7* | 1,434 |
| AP-2 complex subunit mu | *AP2M1* | 1,407 |
| Raftlin | *RFTN1* | 1,400 |
| ADP-ribosylation factor 6 | *ARF6* | 1,400 |
| Macrophage migration inhibitory factor | *MIF* | 1,392 |
| Na(+)/H(+) exchange regulatory cofactor NHE-RF2 | *SLC9A3R2* | 1,391 |
| Unconventional myosin-X | *MYO10* | 1,386 |
| Aldose reductase | *AKR1B1* | 1,384 |
| Ubiquitin-conjugating enzyme E2 L3 | *UBE2L3* | 1,381 |
| Synaptobrevin homolog YKT6 | *YKT6* | 1,379 |
| Transaldolase | *TALDO1* | 1,377 |
| Ras-related protein Ral-B | *RALB* | 1,377 |
| Barrier-to-autointegration factor | *BANF1* | 1,373 |
| Nidogen-1 | *NID1* | 1,361 |
| Protein kinase C and casein kinase substrate in neurons protein 3 | *PACSIN3* | 1,360 |
| V-type proton ATPase 116 kDa subunit a isoform 1 | *ATP6V0A1* | 1,345 |
| Tubulin beta chain | *TUBB* | 1,343 |
| Protein S100-A10 | *S100A10* | 1,340 |
| Ribose-phosphate pyrophosphokinase 1 | *PRPS1* | 1,337 |
| Adenine phosphoribosyltransferase | *APRT* | 1,333 |
| Guanine nucleotide-binding protein G(I)/G(S)/G(O) subunit gamma-10 | *GNG10* | 1,333 |
| von Willebrand factor A domain-containing protein 1 | *VWA1* | 1,328 |
| Arginine--tRNA ligase, cytoplasmic | *RARS* | 1,308 |
| Pre-mRNA-processing-splicing factor 8 | *PRPF8* | 1,298 |
| Polyadenylate-binding protein 1;Polyadenylate-binding protein 3 | *PABPC1;PABPC3* | 1,286 |
| Prefoldin subunit 2 | *PFDN2* | 1,286 |
| Syntaxin-3 | *STX3* | 1,282 |
| Vesicle-associated membrane protein 3 | *VAMP3* | 1,276 |
| Astrocytic phosphoprotein PEA-15 | *PEA15* | 1,276 |
| Cysteine and histidine-rich domain-containing protein 1 | *CHORDC1* | 1,275 |
| Histone H2A type 2-C;Histone H2A type 2-A | *HIST2H2AC;HIST 2H2AA3* | 1,251 |

| | | |
|---|---|---|
| Vimentin | *VIM* | 1,249 |
| ATP-binding **cassette sub-family E member 1** | *ABCE1* | 1,243 |
| Filamin-C | *FLNC* | 1,234 |
| Syndecan-2 | *SDC2* | 1,224 |
| Disco-interacting protein 2 homolog B | *DIP2B* | 1,211 |
| Calnexin | *CANX* | 1,201 |
| Heterogeneous nuclear ribonucleoprotein U | *HNRNPU* | 1,192 |
| Probable ATP-dependent RNA **helicase DDX5** | *DDX5* | 1,176 |
| DNA damage-binding protein 1 | *DDB1* | 1,170 |
| 40S ribosomal protein S23 | *RPS23* | 1,166 |
| Protein disulfide-isomerase A4 | *PDIA4* | 1,165 |
| Poly [ADP-ribose] polymerase 1 | *PARP1* | 1,114 |

**Supplementary Table 3.** Table of the Top 100 proteins often identified in EVs (source: vesiclepedia, http://microvesicles.org/extracellular_vesicle_markers ).

| GENE | NUMBER OF TIMES IDENTIFIED |
|---|---|
| ***PDCD6IP*** | 399 |
| ***GAPDH*** | 377 |
| ***HSPA8*** | 363 |
| ***ACTB*** | 350 |
| ***ANXA2*** | 337 |
| ***CD9*** | 328 |
| ***PKM*** | 327 |
| ***HSP90AA1*** | 327 |
| ***ENO1*** | 327 |
| ***ANXA5*** | 313 |
| ***HSP90AB1*** | 306 |
| ***CD63*** | 306 |
| ***YWHAZ*** | 301 |
| ***YWHAE*** | 300 |
| ***EEF1A1*** | 295 |
| ***PGK1*** | 291 |
| ***CLTC*** | 283 |
| ***PPIA*** | 278 |
| ***SDCBP*** | 277 |
| ***ALDOA*** | 275 |
| ***EEF2*** | 274 |
| ***ALB*** | 274 |
| ***TPI1*** | 270 |
| ***VCP*** | 269 |
| ***CFL1*** | 268 |
| ***MSN*** | 266 |
| ***ATP1A1*** | 266 |
| ***PRDX1*** | 263 |
| ***MYH9*** | 262 |
| ***EZR*** | 262 |
| ***CD81*** | 262 |
| ***ANXA6*** | 260 |
| ***FLOT1*** | 259 |

| | |
|---|---|
| ***YWHAB*** | 258 |
| ***LDHB*** | 258 |
| ***SLC3A2*** | 257 |
| ***GNB1*** | 257 |
| ***PFN1*** | 256 |
| ***TSG101*** | 255 |
| ***YWHAQ*** | 254 |
| ***GNAI2*** | 252 |
| ***CLIC1*** | 251 |
| ***ANXA1*** | 251 |
| ***ITGB1*** | 250 |
| ***LDHA*** | 249 |
| ***FASN*** | 248 |
| ***CDC42*** | 248 |
| ***RAP1B*** | 242 |
| ***CCT2*** | 242 |
| ***YWHAG*** | 240 |
| ***GNB2*** | 240 |
| ***ACTN4*** | 240 |
| ***RAB5C*** | 239 |
| ***C3*** | 239 |
| ***RAB10*** | 236 |
| ***HIST1H4A*** | 234 |
| ***KRT1*** | 233 |
| ***FN1*** | 233 |
| ***AHCY*** | 233 |
| ***A2M*** | 232 |
| ***BSG*** | 230 |
| ***ACTN1*** | 229 |
| ***ANXA7*** | 228 |
| ***ACLY*** | 228 |
| ***HIST1H4B*** | 227 |
| ***GDI2*** | 227 |
| ***FLNA*** | 227 |
| ***UBA1*** | 226 |
| ***GNAS*** | 226 |
| ***GSN*** | 225 |
| ***CCT4*** | 225 |
| ***RAN*** | 222 |
| ***PRDX2*** | 222 |
| ***RHOA*** | 220 |
| ***CCT3*** | 220 |
| ***RAC1*** | 219 |
| ***LGALS3BP*** | 219 |
| ***TCP1*** | 218 |
| ***KRT10*** | 218 |
| ***CAP1*** | 218 |
| ***RAB7A*** | 217 |
| ***TUBB4B*** | 216 |
| ***HSPA5*** | 215 |
| ***IQGAP1*** | 214 |

| | |
|---|---|
| ***GPI*** | 214 |
| ***RALA*** | 213 |
| ***KPNB1*** | 212 |
| ***HIST1H4I*** | 212 |
| ***TFRC*** | 211 |
| ***EIF4A1*** | 211 |
| ***HIST4H4*** | 210 |
| ***CCT8*** | 210 |
| ***TLN1*** | 209 |
| ***HIST1H4K*** | 209 |
| ***HIST1H4H*** | 209 |
| ***CCT6A*** | 209 |
| ***ANXA11*** | 209 |
| ***HIST1H4J*** | 208 |
| ***HIST1H4F*** | 208 |
| ***HIST1H4D*** | 208 |
